# Mammalian Retinal Bipolar Cells: Morphological Identification and Systematic Classification in Rabbit Retina with a Comparative Perspective

**DOI:** 10.1101/2024.09.19.613998

**Authors:** Edward V. Famiglietti

## Abstract

Retinal bipolar cells (BCs) convey visual signals from photoreceptors to more than 50 types of rabbit retinal ganglion cells (Famiglietti, 2020). More than 40 years ago, 10-11 types of bipolar cell were recognized in rabbit and cat retinas (Famiglietti, 1981). Twenty years later 10 were identified in mouse, rat, and monkey, while recent molecular genetic studies indicate that there are 15 types of bipolar cell in mouse retina (Shekhar et al., 2016). The present detailed study of more than 800 bipolar cells in ten Golgi-impregnated rabbit retinas indicates that there are 14-16 types of cone bipolar cell and one type of rod bipolar cell in rabbit retina. These have been carefully analyzed in terms of dendritic and axonal morphology, and axon terminal stratification with respect to fiducial starburst amacrine cells. In fortuitous proximity, several types of bipolar cell can be related to identified ganglion cells by stratification and by contacts suggestive of synaptic connection. These results are compared with other studies of rabbit bipolar cells. Homologies with bipolar cells of mouse and monkey are considered in functional terms.

## 1.0 INTRODUCTION

As Santiago Ramon y Cajal noted, in his seminal work on the retina (Cajal, 1893), Tartuferi (1887) first appreciated the key role of bipolar cells in connecting photoreceptors to retinal ganglion cells, using Golgi’s method of silver impregnation (Golgi, 1875). Applying Golgi’s method widely among the vertebrates, in sectioned material and in retinal whole-mounts, Cajal (1893) understood that bipolar cells were diverse, consisting of rod and cone bipolar cells. Among the latter, he found that their axon terminals branched variously in different strata of the inner plexiform layer (IPL), in order to contact ganglion cells branching.at these same levels.

Functional significance, attributable to such pairings of bipolar and ganglion cells in different strata of the IPL, escaped scientific understanding for eighty-three years (Famiglietti et al., 1977; Famiglietti & Kolb, 1976; Nelson et al., 1978). This gap in understanding may have been due in part to the fact that individual bipolar cells, beautifully illustrated by Cajal in his drawings from retinas of non-mammalian vertebrates, may contribute to one, two, and even four different strata of the IPL (Cajal, 1893). Such variation in form and stratification of non-mammalian bipolar cells, is largely absent in Cajal’s illustrations of mammalian bipolar cells, the diversity of which became apparent only much later in Golgi studies first of primate retina (Polyak, 1941) and then of rabbit retina (Famiglietti, 1981).

Recently, the application of several powerful techniques, including immunocytochemistry, 3D-reconstruction of cells from serial thin sections imaged by electron microscopy, and single cell RNA sequencing, show that there are at least 14 types of cone bipolar cell in mouse retina (Helmstaedter et al., 2013; Shekhar et al., 2016; Wassle et al., 2009). Nevertheless, 136 years after Tartuferi’s characterization of retinal bipolar cells, the silver impregnation method retains its power to delineate the entire, membrane-bound shape of a nerve cell with unsurpassed clarity at the light microscopic level of examination. The present Golgi study extends work begun in 1981 (Famiglietti, 1981), and more recent additions to that work (Bordt et al., 2019; Famiglietti, 2002, 2008), to show that the number of cone bipolar cells types in rabbit is comparable to and may exceed that demonstrated in mouse.

## 2.0 METHODS

The tissue preparation, Golgi impregnation, and analytical methods for studying retinal bipolar cells have been laid out in detail elsewhere (Famiglietti, 1992a, 1992b). The methods used conformed to institutional and NIH guidelines. A synopsis of these methods is presented below.

### 2.1 Golgi-impregnation

Golgi-impregnated rabbit retinas were prepared according to a Golgi-Kopsch-Colonnier method described elsewhere (Famiglietti, 1985). Briefly, pigmented rabbits (Dutch-belted, New Zealand red or black, and indeterminate strains with brown or black iris) were surgically enucleated, painlessly, under deep halothane anesthesia, and were euthanized by halothane overdose, followed by thoracotomy and exsanguination. The retinas were isolated from the posterior half of the eye under dim light in buffer, and were flattened by making radial cuts from the periphery toward the center. They were then mounted in a sandwich of filter paper, which was held flat by a coverslip and rubber bands on a glass slide and were then fixed in aldehydes. The slide was immersed in an aldehyde-dichromate mixture, and then in 1% silver nitrate, for a few days each. Retinas were dehydrated and mounted whole on glass slides either in DPX or in sheets of Epon/Araldite.

### 2.2 Retinal location of bipolar cells

Details of the methods for determining the location of mid-visual streak in each Golgi-impregnated retina and for accurate mapping of the retinal location of each cell are given in a previous communication (Famiglietti, 1992a). Thousands of bipolar cells were observed in 26 retinas; 794 Golgi-impregnated cone bipolar cells and numerous rod bipolar cells were closely examined. These were located to the nearest 50 microns, taking X and Y coordinates from the microscope stage micrometer and mapping them onto 10X or for the five most productive retinas onto 100X camera lucida drawings of the retinas. Most of the bipolar cells studied were in the region from 2 mm dorsal to 8 mm ventral to mid-visual streak (−2.0 < *dvs* < +8.0).

### 2.3 Measurement and analysis of dendritic field and axon terminal field sizes

Camera lucida drawings were made of all cells with a Zeiss drawing tube at magnifications ranging from 575X to 2350X, the great majority at 2000X. Detailed drawings at high magnification were made of 708 cells and 772 cells were suitable for measurement of axon terminal field and dendritic trees.. Methods of determining field areas using best-fitting convex polygons are described elsewhere (Famiglietti, 1992a). Areas were converted to “equivalent dendritic field diameters”, according to the equation d_eq_ = 2√(A/π), where A = dendritic field area.

Corrections were not made for tissue shrinkage in Golgi preparations due to fixation and dehydration, when the sizes of Golgi-impregnated cells were graphed. Shrinkage has been estimated in this material to range up to 30% in all dimensions (Famiglietti, 1985), and is typically 15-25%.

### 2.4 Graphic analysis

Graphs were constructed in Sygraph (Systat^TM^). Field sizes were plotted as a function of distance from the visual streak (*dvs*). Curves were fitted to the distributions usually by the linear method of least squares and sometimes by distance weighted least squares (DWLS).

### 2.5 Microscopic analysis of dendritic stratification

Analysis of dendritic stratification was performed for the most part in flattened, whole-mounted retinas, as described elsewhere in detail (Famiglietti, 1992b), using high magnification, high numerical aperture oil-immersion lenses and an accurate, gear-driven (Zeiss WL) microscope stage elevator. Dendritic stratification in the retina must be analyzed against a reference layer or layers.

The most valuable material for the analysis of dendritic stratification, which can lead to a useful representation of stratification, substratification, and co-stratification of dendritic trees in the inner plexiform layer (IPL) of rabbit, is the whole, unsectioned, flat-mounted retina containing numerous instances of overlapping dendritic trees. In these instances, “marker” cells constitute fiducials against which other less well-known cells can be studied. Consistency of relationships in level of stratification can thus be represented against a standard. The most useful marker cells in the IPL of the rabbit for this study are the type a and type b starburst amacrine cells, the type 1, bistratified (BS1) or ON-OFF DS ganglion cell, class I and class II cells (Famiglietti, 2004b) and the class IV, type 1 small tufted (ST1) (local edge detector) (Famiglietti, 2005). For graphic analysis and demonstration of stratification, except in a few cases, only cells referenced to the fiducials of starburst amacrine cells or type 1 bistratified (ON-OFF directionally selective) ganglion cells were used. Those exceptions were generally based upon mutual overlap with wide-field, unistratified radiate amacrine cells. When available in proximity to bipolar cells, AII amacrine cells were useful both in delimiting the depth of the IPL, and determining the position of the a/b sublaminar border.

### 2.6 Camera lucida drawings, digital photography and reproduction

Camera lucida drawings were traced using a Zeiss RA microscope fitted with a Zeiss drawing tube and oil-immersion 40X or 100X NA = 1.0 long-working-distance microscope objectives. Drawings were digitized at 600 dpi on a flat-bed scanner and imported into Adobe Photoshop 6, where contrast and brightness were adjusted.

Photographs were taken through a 25X, 40X oil, or a 100X oil objective lens with a Canon G6 digital microscope camera, using Canon Zoombrowser software, and assembled and processed in Adobe Photoshop. Serial optical sections of Golgi preparations were photographed at 0.5 micron intervals on a Zeiss WL gear-driven microscope stage, using a Zeiss Neofluar 100X oil immersion objective. In many cases, the retinal whole-mounts required the use of a long-working distance lens (WD = 600 µm), NA 1.0, with a theoretical resolution of 0.32 µm. In some cases, particularly for photography, it was possible to use a Leitz 100X oil immersion, NA = 1.32, with a 180 µm working distance and a theoretical resolution of 0.26 µm.

### 2.7 Immunocytochemistry and analysis of neuronal stratification in the IPL

Well-characterized primary antibodies used in this study were as follows: 1) rabbit anti-chick choline acetyltransferase (ChAT), #1465, a gift of Myles Epstein, 2) monoclonal anti-Calbindin-D (28kD) made against chicken gut purchased from Sigma Immunochemicals, No. C8666, and 3) rabbit anti-recoverin, purified from bovine retina, a gift from A. M. Dizhoor to W. K. Stell (Dizhoor et al., 1991). The first of these, shown to stain cholinergic/starburst amacrine cells in rabbit retina (Famiglietti & Tumosa, 1987), has been characterized immunologically (Millar et al., 1985), and reveals a similar pattern of staining in all mammals studied. The second and third primary antibodies have been used previously (Massey & Mills, 1996; Sharpe et al., 1993; Zhang et al., 2005) to mark the same bipolar cells studied here.

Eyes were obtained from adult female New Zealand red and New Zealand white rabbits. Adult rabbits were premedicated prior to tracheal intubation and then anesthetized with halothane. Eyes were enucleated successively under anesthesia, after which the animal was euthanized according to approved protocols. The anterior segment of the eye was removed in phosphate-buffered saline (PBS), and the posterior eyecup placed in fixative, consisting of 4% paraformaldehyde and 0.1% glutaraldehyde in 0.1M Sorensen’s Phosphate buffer at pH= 7.2 for 15 minutes at room temperature. Enucleation and tissue fixation were carried out under ambient room light. After an hour in the same fixative at 4°C, the eyecups were left overnight in 4% paraformaldehyde buffered at pH= 10.4 by carbonate-bicarbonate buffer. The next day, after two 30 minute rinses with and without sodium borohydride, the retina was dissected from the eyecup and cut into the required pieces. Pieces intended for frozen-sectioning were placed in 30% sucrose with 0.1% sodium azide overnight. Material for sectioning was mounted in low-temperature gelling agarose and sectioned at −20°C, generally at 12 μm. Sections were mounted on Fols subbed slides and stored at −20° C until processing.

Thawed, slide-mounted sections were incubated in 10% normal goat serum for 30 minutes, rinsed in PBS, and then incubated overnight, within a humid chamber at room temperature, in primary antibody diluted in PBS containing sodium azide and Triton-X100, typically at 0.03%. Primary antibody dilutions were 1:2500 (calbindin), 1:1000 (anti-recoverin), and 1:500 (anti-ChAT). Double labeling was achieved by using secondary antibodies conjugated to different fluorophores. After two PBS washes, sections were incubated in secondary antibodies appropriate to primary antibodies, conjugated either to FITC or to Cy3 for up to one and one-half hours, followed by two washes in PBS and mounting in glycerin or Fluorosave R. Slides were checked and then stored at −20° C for up to two weeks prior to photomicrographic documentation.

Slides of sectioned material were photographed in a Zeiss Universal fluorescence photomicroscope, using band-pass filter sets (Omega) for FITC (ex.: 485DF22; em.: 530DF20 nm) and Texas Red (ex.: 555DF27; em.: 615DF45) to adequately separate signals from FITC and Cy3 fluorescence in the same double-labeled section.

Water immersion Zeiss Neofluar 16X and 25X lenses were used to image the fluorescent labels, and 35mm film was exposed, in black and white (Kodak T-MAX 400). Digital images were made from film and were processed in Adobe Photoshop for contrast and brightness, and were pseudo-colored to facilitate viewing of immunoreactive cells and processes.

As a supplement to the analysis of stratification in the IPL, optical densitometry was used to quantify the positions of immunocytochemically demonstrated bands of label in transverse sections of retina. Photographs of individual sections were digitized and imported into Image (NIH Image), and densitometry performed using gel analysis software. Scans were compared in double-stained preparations. In comparisons of single stained preparations, scans were adjusted if necessary to equivalent thickness of the IPL. In the latter case, scans were aligned with reference to ChAT-IR to compare patterns of stratification for ChAT and recoverin.

### 2.8 EM immunocytochemistry

Eyes were obtained from a New Zealand white rabbit, as described above, the anterior segment and vitreous humor removed, and the eyecup fixed in 2.5% paraformaldehyde and 0.01% glutaraldehyde with 2% DMSO overnight at 4° C. Small squares of retinal were transferred to 20% DMSO and frozen on filter paper in liquid nitrogen, thawed, encapsulated in low-temperature gelling agarose, and sectioned on a Vibrotome at 75 µm.

Sections were incubated in anti-ChAT, #1465,a gift of Miles Epstein (Johnson & Epstein, 1986), at 1:500 dilution, with 2% DMSO in PBS-ovalbumin (OA) for two days, and then in goat anti-rabbit IgG, followed by peroxidase antiperoxidase, 1:100 with 2% DMSO in PBS-OA five hours, and then diaminobenzidine for 15 minutes. Post-fixation in1% glutaraldehyde was followed by osmication (1% OsO_4_, 1.5% K_3_Fe(CN)_6_), tannic acid incubation (1%), en bloc uranyl acetate staining (2%), methanol dehydration, immersion in dioxane, and epoxy infiltration in a dioxane solution. Epoxy embedding was carried out in an Epon/Araldite mixture. Thin sections were cut on an AO Ultracut microtome at 0.75 nm, stained in 5% uranyl acetate and Reynolds’ lead citrate, and viewed in a Phillips 400 electron microscope, in which electron micrographs were obtained.

## 3.0 RESULTS

### 3.1 Cone bipolar cells in rabbit retina

Sixteen types of cone bipolar cell have been identified in this study. Seven are type a bipolar cells with axon terminals branching in sublamina a, and nine are type b cells, branching in sublamina b of the inner plexiform layer (IPL) (Figure 1). Eleven are narrow field cells, “na” or “nb”, three are medium wide field cells, “ma” or “ mb”, and two are wide-field cone bipolar cells, “wa” or “wb”.

**FIGURE 1.**
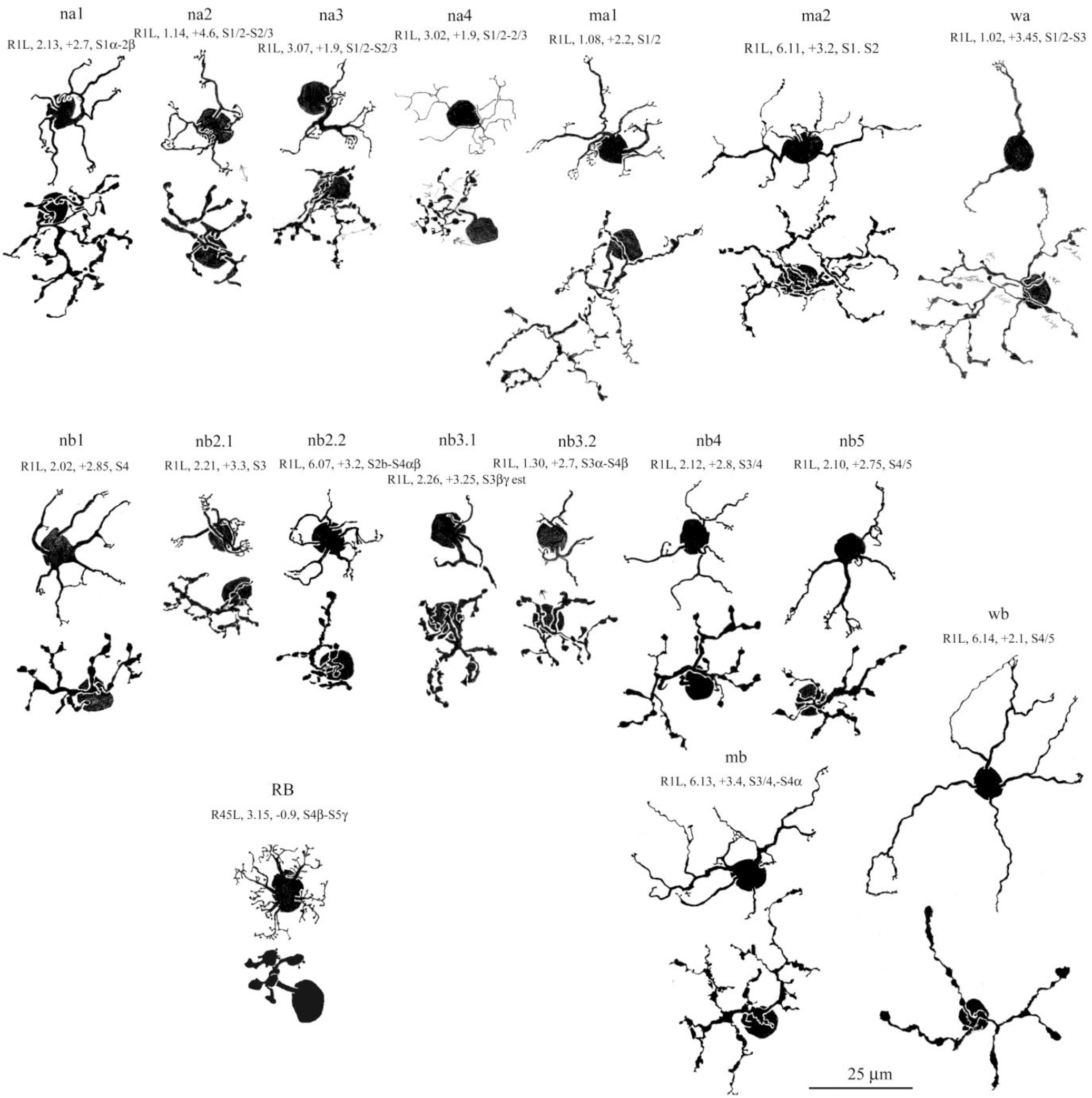
The bipolar cells of rabbit retina. Camera lucida drawings of 16 types of cone bipolar cell, and one type of rod bipolar cell, viewed *flat* (horizontally) in whole mounted retinas. For comparison, all cone bipolar cells taken from a single retina, R1L. The identifications listed over each cell specify, in order: retina #, reference #, *dvs*= distance (mm) dorsal (–) or ventral (+) to mid--visual streak, and stratum (S) of axon terminal branching. Classification: n = narrow-field, m = medium-wide field, w = wide-field; a, b = sublamina a, b of the inner plexiform layer.

### 3.2 Type a cone bipolar cells

In a previous study of rabbit bipolar cells (Famiglietti, 1981), several types of cone bipolar cell were identified that branch in sublamina a of the IPL. These include three narrow field types: na1, na2, and na3, a medium wide field type: ma, and a wide field type: wa. In that study, it was proposed that wa cone bipolar cells connect with blue (short wavelength) cones, exclusively, and “blue cone pathways” to ganglion cells were subsequently described (Famiglietti, 2008). Recently, a second type of medium wide field “type a” cone bipolar cell was discovered (Bordt et al., 2019), favoring a renaming of the first type “ma1”, and the new variety “ma2”. In the first study of rabbit bipolar cells (Famiglietti, 1981), a narrow field type of cone bipolar cell with an axon terminal that appeared to cross the a/b sublaminar border was designated “nab”. For reasons related to its axon terminal stratification (see below), the former nab cells are now designated “na4” cone bipolar cells. Thus it now appears that there is a total of seven kinds of type a cone bipolar cell in rabbit retina (Figure 1).

In the present effort to provide a complete survey of bipolar cell types in rabbit retina, a more detailed description of each is warranted. In addition to morphological features of their dendritic trees, detailed analysis of the level (s) of their axon terminal stratification in the IPL receives special attention here, and use is made of the standard 5-tiered stratification scheme shown in Figure 2b. Of particular value for the analysis of stratification are the two narrow bands of starburst amacrine cell dendrites, and starburst amacrine cells, whether immuno-stained with antibodies to choline acetyltransferase (ChAT) (Figure 2) or Golgi-impregnated, are the fiducial cells of choice for measuring the stratification of bipolar cells in this study.

**FIGURE 2.**
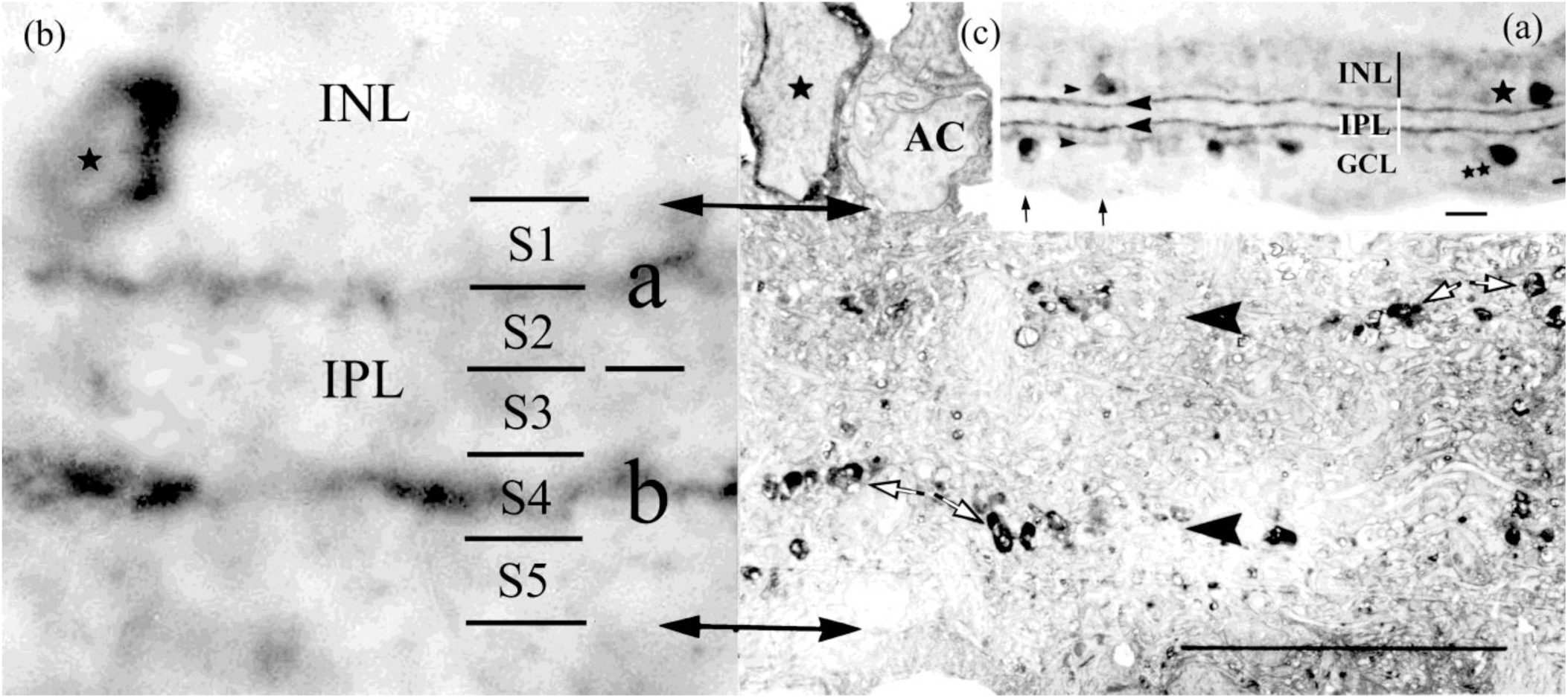
Choline acetyltransferase immunocytochemistry (ChAT-ICC) of retinal sections, staining starburst amacrine (SA) cells, principal fiducial markers for analysis and measurement of the axon terminal stratification of bipolar cells in the inner plexiform layer (IPL). The IPL, inner nuclear layer (INL), and ganglion cell layer (GCL) are viewed in transverse sections by light microscopy (a), (b) and electron microscopy (c), with alignment of strata at equal magnification in (b) and (c). Single star symbols indicate immunoreactive type a SA cells, and a double star marks a type b SA cell in (a). An unstained amacrine cell soma (AC) is marked in (c). (a) Rightward arrow heads: borders of the IPL; leftward arrowheads: two bands of ChAT-IR in the IPL. Vertical arrows: location of ICC-reaction-intensity scans graphed in Figure 29c. (b) Arrows: boundaries of the IPL. A standard 5-tier schema of stratification is superimposed. Strata 1 and 2 are in sublamina a; strata 3, 4, and 5 are in sublamina b. The ChAT-IR band in sublamina a, indicating type a starburst (SAa) cells, lies at the S1/S2 border; the band in sublamina b, indicating type b cells, lies in the middle of S4 in sublamina b. Calibration bars: 15 µm.

#### 3.2.1 na1 cone bipolar cells

In previous work (Famiglietti, 1981), a parallelism was drawn between particular narrow field type a and type b cone bipolar cells, especially in regard to their dendritic branching patterns, reflected in the nomenclature: na1 with nb1, and na2 with nb2. The n1 varieties were described as having “dendritic branches of medium thickness, each of which ends in a small solitary cluster of ascending terminal appendages (e.g. Figure 1). More thoroughgoing analysis shows that this description of dendritic appendages is more consistently seen in nb1 cells, however, as described below (cf. Figures A1 and A5).

The na1 cells do share with nb1 cells a radiating dendritic branching pattern. They also share a predominantly divergent branching pattern in contrast to the convergent pattern in na2 and nb2 cells as previously described (Famiglietti, 1981). As a consequence of this radiating-divergent pattern both na1 and nb1 cells have the largest dendritic trees at a given retinal location among narrow field type a and type b cells, respectively (Figures 3c and 3d). In this configuration, they seldom converge upon the same cone. In regard to terminal appendages of the distal most branches, na1 cells often have single, or a splayed pair of digitiform appendages, but may have an irregular flat array of as many as five such appendages, extending over as much as 4 µm in the plane of the retina (Figure A1).

**FIGURE 3.**
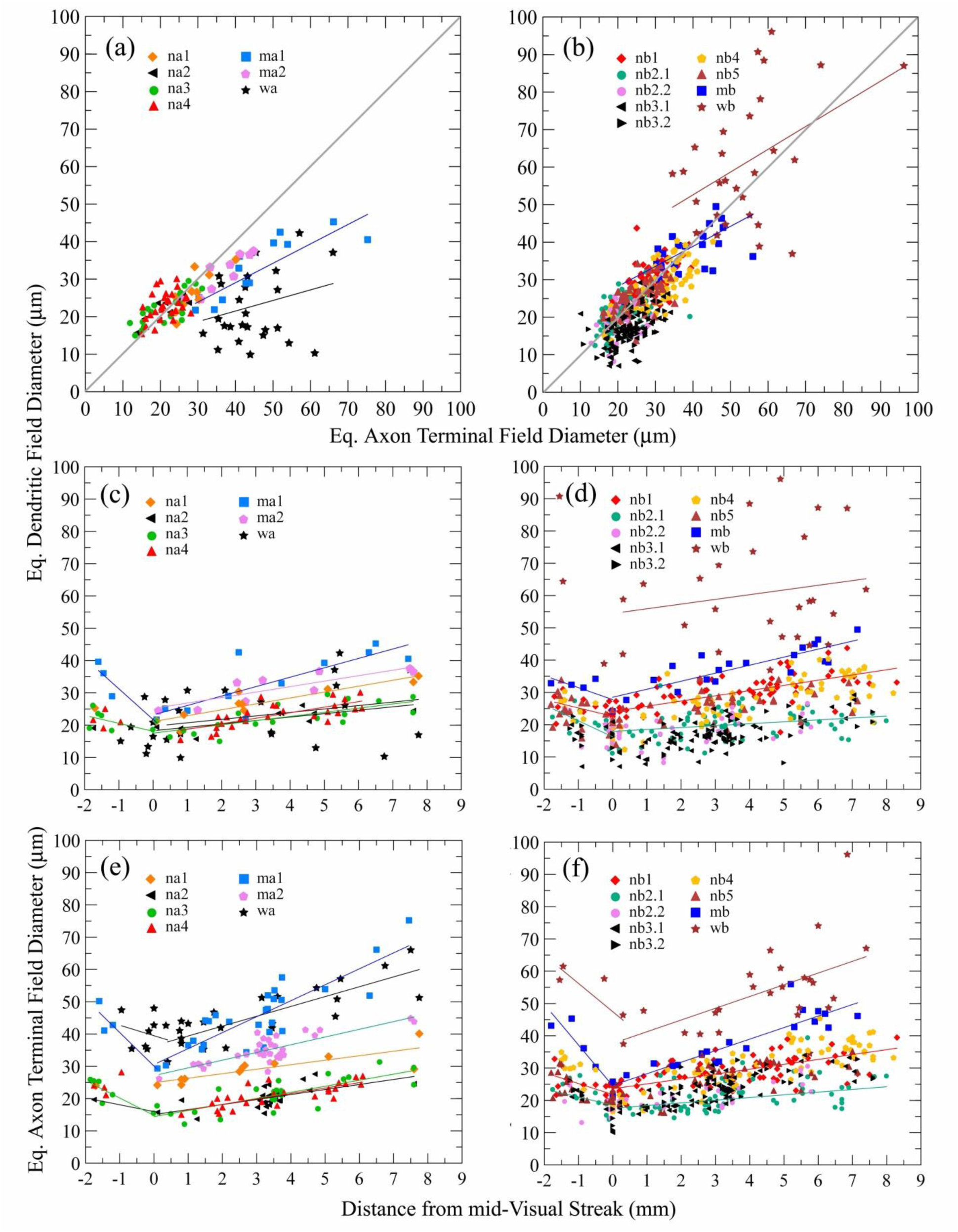
Graphs displaying measurements of bipolar cell dendritic tree and axon terminal field areas in the plane of the retina, converted to “equivalent diameters”. (a) and (b) The dendritic trees and axonal arborizations of narrow field cells are about the same size for both type a and type b bipolar cells. Wide-field cells differ significantly, wa cells having smaller dendritic trees on average, and wb cells having the largest dendritic trees. Medium-wide field cells differ less in this regard, but ma1 cells are notable for having larger axon terminal fields. Variation in dendritic field size of type a cells (c) and type b cells (d) is plotted with distance from mid-visual streak. Note that wb cells are much larger than all others, while wa cells are among the smallest. Variation in axon terminal field size of type a cells (e) and type b cells (f) shows that wa and wb cells are comparably large, and ma1 cells, although morphologically matched to mb cells, are larger than the latter and are comparable in axon terminal field size to wide field bipolar cells.

#### 3.2.2 na2 cone bipolar cells

The na2 cells were originally described as having “thicker, twisted dendrites which converge to form masses of small, ascending terminal digits” (Famiglietti, 1981). While this description was intended for the dendrites of both na2 and nb2 cells, the reference to “masses of… ascending terminal digits” more aptly applies to nb2.1 cells, described below. The somewhat more compact and convergent dendritic pattern of na2 cells, as compared to na1 cells, can be appreciated in camera lucida drawings (Figure 4 and A1). As a consequence of these features, na2 cells have smaller dendritic trees than na1 cells (Figure 3c).

**FIGURE 4.**
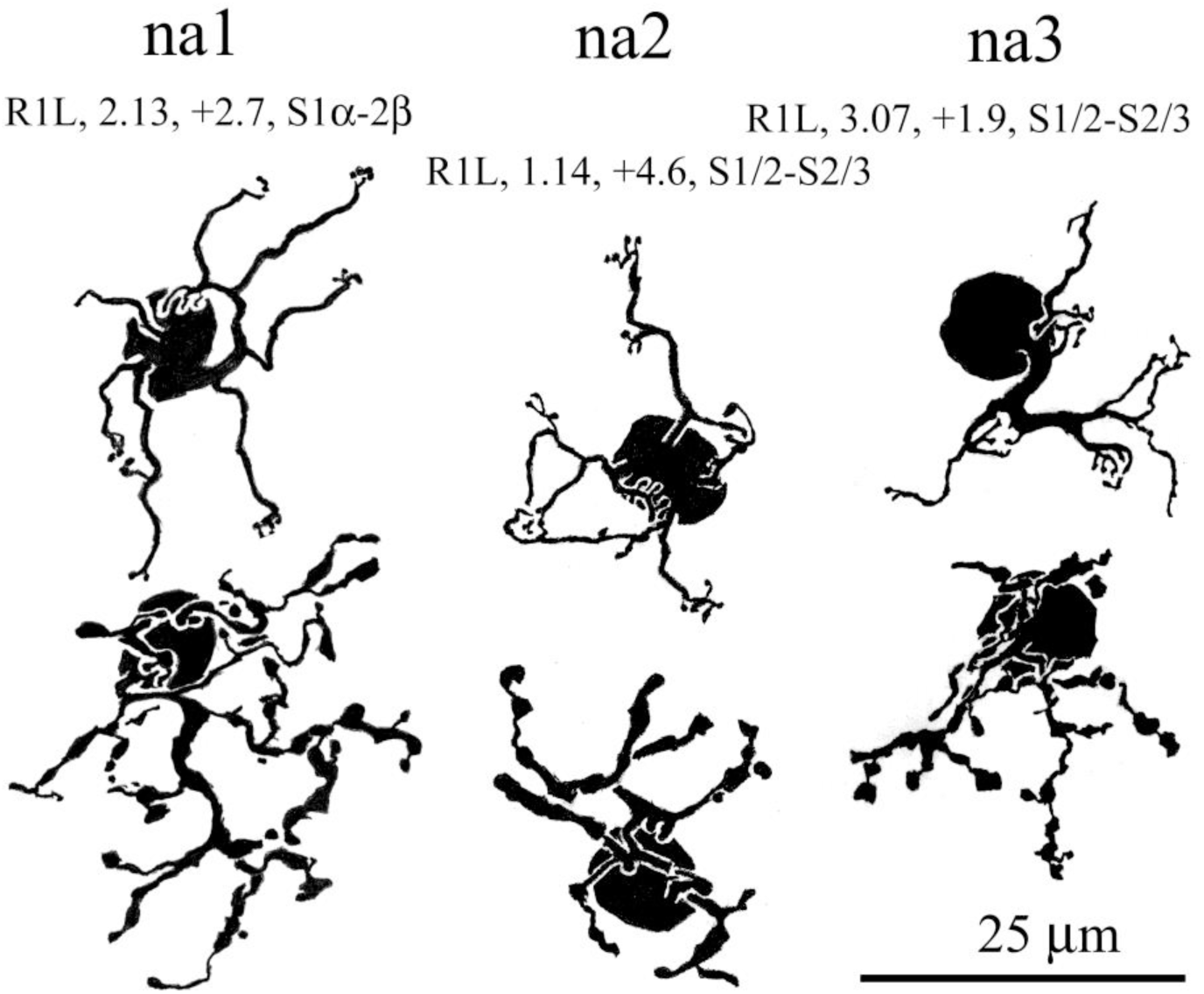
Three kinds of narrow-field (n), type a cone bipolar cells in rabbit retina. Each has distinctive dendritic branching, while axon terminal stratification of na1, na2, and na3 differs in more subtle ways (see text and Figures 5 and 6).

As is commonly the case for narrow field cone bipolar cells in rabbit, the size of the axon terminal is generally commensurate with the size of the dendritic tree (Figure 3a and b). Hence, in regard to the former, na1 cells are larger than na2 cells (Figure 3e). As well as being smaller, the axon terminals of na2 cells are less complex, and their terminal lobular appendages are slightly more robust when compared to those of na1 cells (Figure A1). The axon terminal stratification of na1 and na2 cells also differs: na1 cells branch predominantly in stratum (S)1 and na2 cells are biased toward S2 (Figure 5).

**FIGURE 5.**
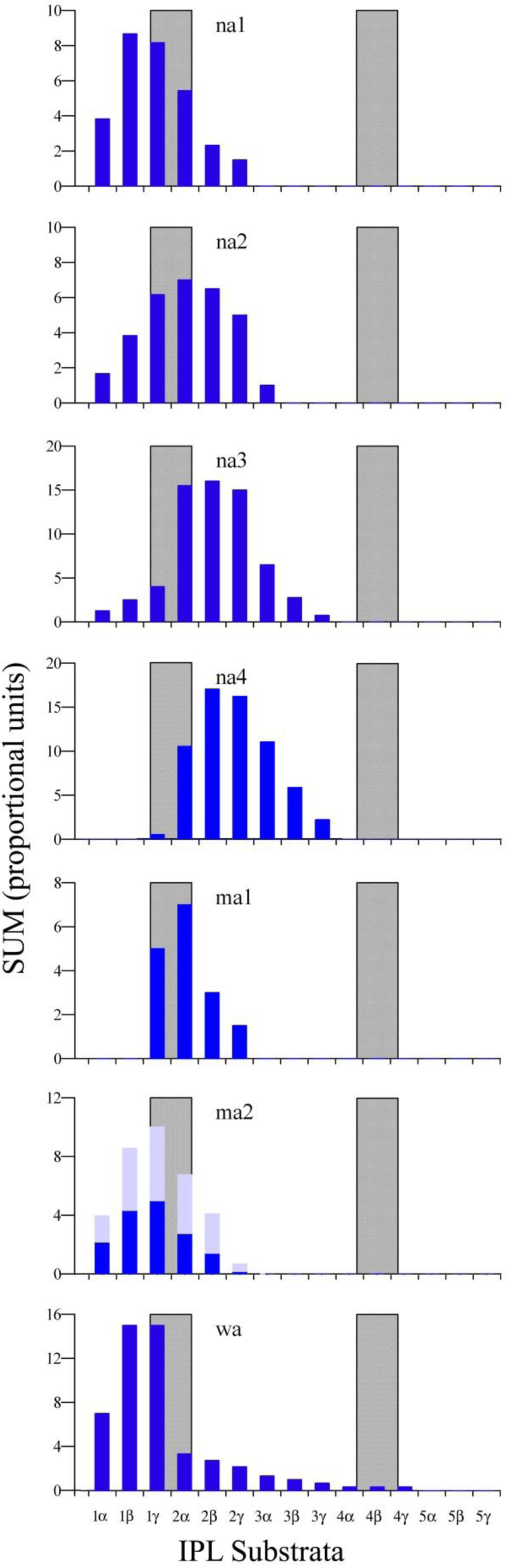
Axon terminal stratification of the seven kinds of type a cone bipolar cell in rabbit retina. Each stratum is subdivided into three substrata. The stratification of each cell is determined with reference to a fiducial cell (usually SAa), and the portion in each substratum is given a value. Then the values are summed across substrata and plotted. Gray bars represent the location and breadth of SAa and SAb dendritic laminae. Blue bars: stratification based upon fiducial cells; light blue bars: stratification estimated (see text)

#### 3.2.3 na3 cone bipolar cells

The third narrow field type a cone bipolar cell, na3, does not differ significantly from na2 cells in the size of their dendritic trees or axon terminal arbors (Figure 3a, c), but has distinctive and characteristic features of dendritic branching, and of axon terminal stratification. The na3 cell has converging dendrites in a compact dendritic tree with one to three foci, centered on cone pedicles, of a complex, flat array of digitiform appendages, typically forming ring-like structures (Figures 4 and 6), as noted in previous work (Famiglietti, 1981). These span the base of a typical cone pedicle, with which they are commensurate, as shown in Figure 6b.

**FIGURE 6.**
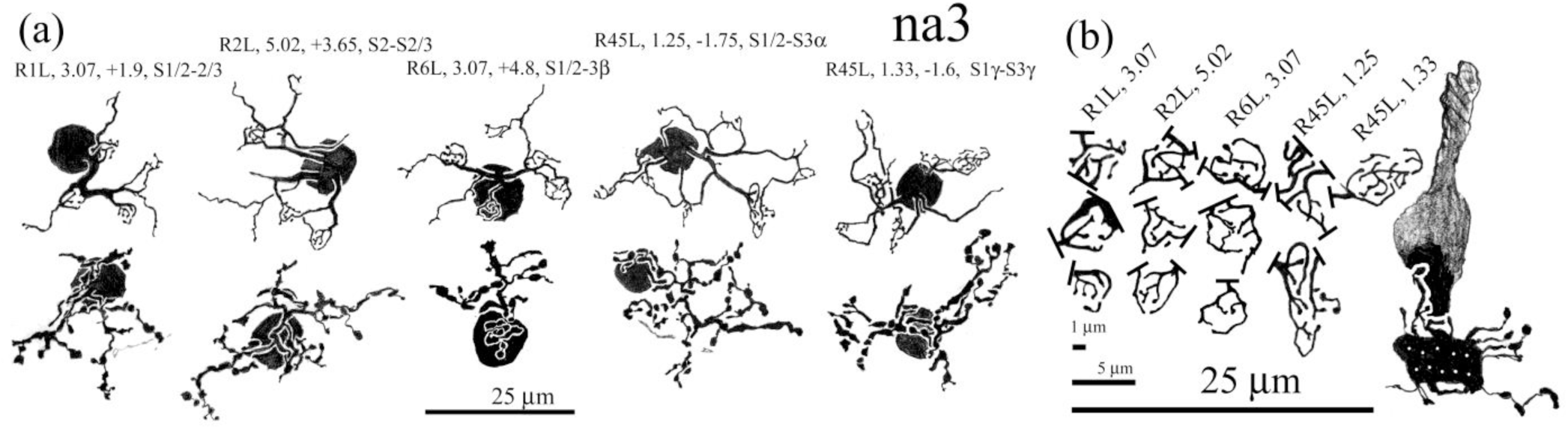
(a) na3 cone bipolar cells, taken from four retinas. (b) details of the nest-like dendritic terminal appendages of the bipolar cells in (a), showing that the individual areas of digitiform appendages are commensurate with the cone pedicle base (b). Dots in the pedicle are an estimate of the number and distribution of synaptic ribbons.

Morphologically, na3 axon terminals are usually complex with lobular appendages, both scalloped and rounded, varying in size and compact in the retinal plane, branching at a moderately high frequency (Figure A1). Their axon terminals resemble those of na1 cells more than those of na2 cells (Figure 4), except for their vertical rather than horizontal disposition with respect to the retinal plane. The axon terminals of na3 cells typically contribute little to S1, and branch mostly in S2 (Figure 5). In the previous study (Famiglietti, 1981), the axon terminals of na3 cells were thought to be confined to sublamina a of the IPL, like na1 and na2 cells. A more thorough analysis, with the aid of fiducial cells overlapping in the retinal plane, reveals, however, that na3 cells sometimes are not so confined. In some retinas, but not others, they extend into S3 (Figure A2).

#### 3.2.4 na4 cone bipolar cells

In the previous study (Famiglietti, 1981), some types of bipolar cells were described that straddle the a/b sublaminar border, including type “nab” bipolar cells. At the time, this was a matter of some concern, in the wake of the discovery that, in general, the a/b sublaminar border separates OFF/hyperpolarizing and ON/depolarizing inner retinal neurons forming synapses in the IPL, in both mammalian and teleost retinas (Famiglietti et al., 1977; Famiglietti & Kolb, 1976; Nelson et al., 1978). Furthermore, multistratified cone bipolar cells that cross such a border were known to be common in non-mammalian vertebrates (Cajal, 1893). Against this background, the axon terminals of rabbit nab cone bipolar cells were described as either confined to sublamina a or overlapping the a/b sublaminar border (Famiglietti, 1981).

More thoroughgoing analysis of nab cells shows that they vary from animal to animal in concert with na3 cells (Figure A2). Study of nearby na3 and nab (na4) cells, overlapped by the same fiducial type a starburst amacrine cell (Figure 7), viewed with a high resolution microscope objective, reveals that there is little difference in stratification between the two, and, in the latter case, justifies the change in nomenclature from nab to na4. Moreover, the axon terminal morphology of na3 and na4 cone bipolar cells is similar (cf. Figures A1 and A2). That cannot be said about the dendrites of these two types, which represent morphological extremes among bipolar cells of rabbit retina (Figures 1 and 7). In contrast to the rings of digitiform appendages, borne by the typically robust dendrites of na3 cells, na4 cells usually have slender dendrites branching in a skein of fine processes with few distinct terminal appendages (Figures 7 and A2). Their dendritic trees do not differ significantly in size, however (Figure 3c).

**FIGURE 7.**
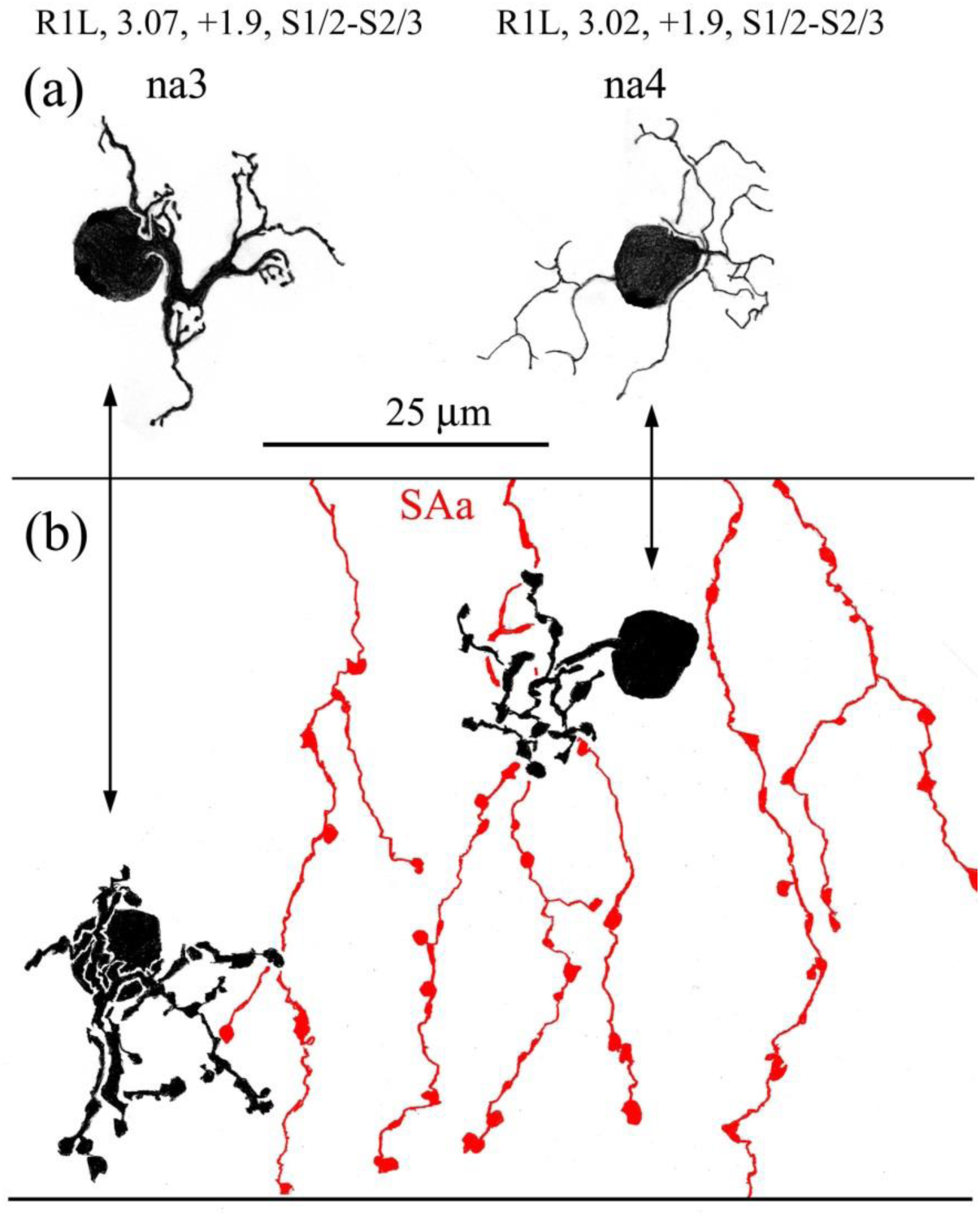
Comparing na3 and na4 cone bipolar cells. (a) dendritic branching, (b) axon terminal branching with overlapping fiducial type a starburst amacrine cells (SAa, red). Microscopic study shows that both have branches at approximately the same level.

#### 3.2.5 ma1 cone bipolar cells

Formerly, a single medium-wide field type a cone bipolar cell was described (Famiglietti, 1981), but recently, another medium-wide field type a cell was found (Bordt et al., 2019). Hence the name of the former is changed here to “ma1”, while the new find is “ma2”. On average, ma1 cells have the largest dendritic fields among type a bipolar cells (Figure 3c), and their axon terminal fields, significantly larger than their dendritic fields (Figure 3a), and much larger that the axon terminal fields of narrow field cells, are comparable in size only to wide field cells (Figures 3e and f).

The dendritic trees of ma1 cells are mostly radiating and only moderately convergent, forming in each case a few arrays of digitiform appendages (Figures 8 and A3). These are reminiscent of those formed by na3 cells, but not as complex, and are more often distributed peripherally at terminal branches. Some of the terminal dendrites end simply, unadorned by terminal appendages, and some of these have been reported to contact rod photoreceptor spherules (Li et al., 2004). The axon terminals of ma1 cells are narrowly stratified, most often at the S1/S2 border (Figure 5), and extensive in a radiating, rather sparsely branched distribution of lobulated processes. Their branches include segments of both varicose and beaded contour (Figure A3).

**FIGURE 8.**
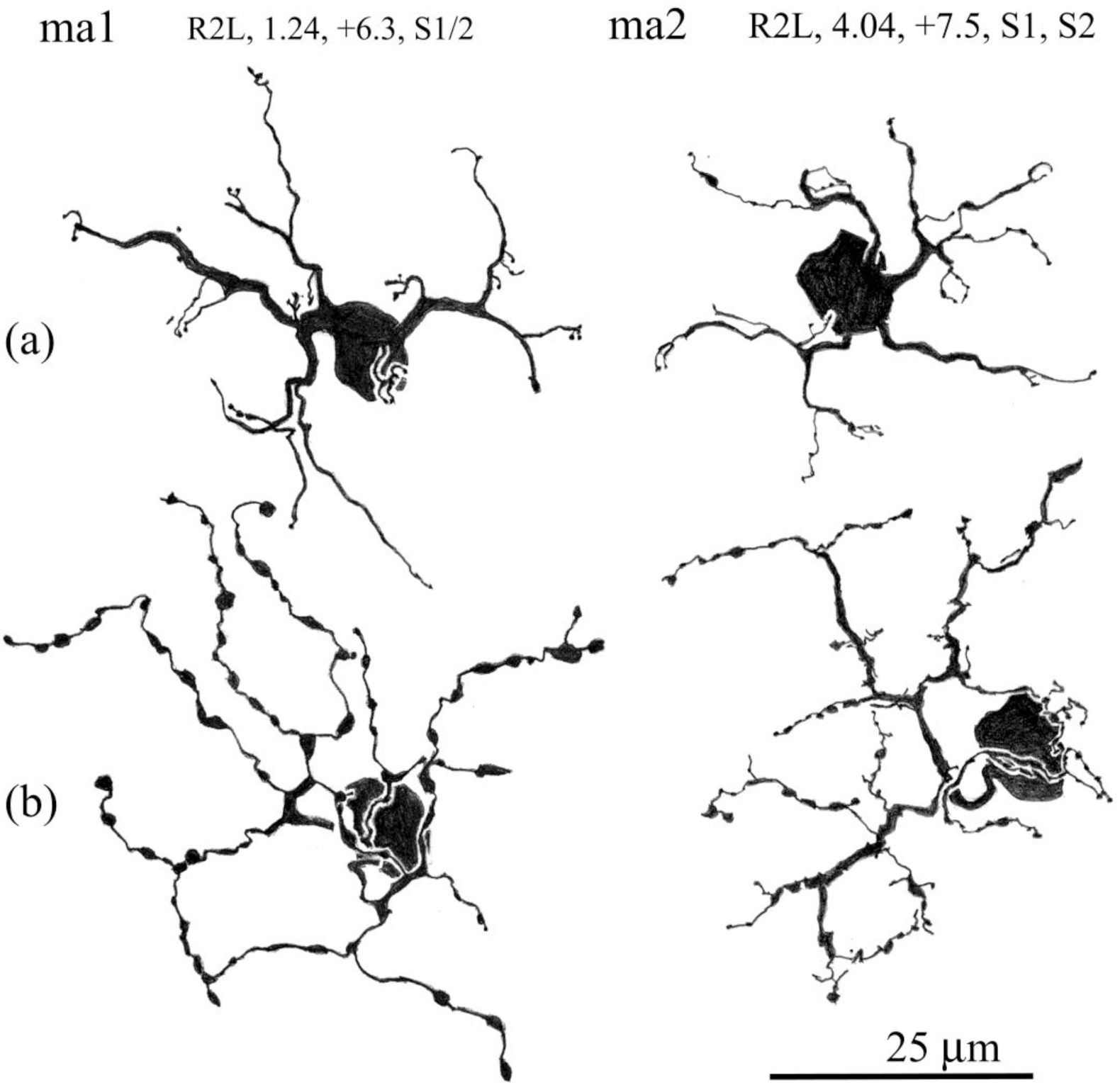
Comparing ma1 and ma2 cone bipolar cells. Both have similar extended dendritic and axon terminal branching patterns. (a) The dendrites of ma1 cells bear loose clusters of spines in irregular clusters of digitiform appendages at the ends of terminal dendrites. The dendrites of ma2 cells are more irregular, in comparison. (b) ma1 axon terminals have a radiating pattern and often a beaded appearance, as seen here. The axon terminals of ma2 cells are unique, exhibiting long stretches of relatively thick processes of uniform caliber, bearing much thinner offshoots (see text).

#### 3.2.6 ma2 cone bipolar cells

The ma2 cone bipolar cell was missed in the previous studies. It was rarely impregnated in all but one retina among those studied here. Moreover, at first glance it is also relatively nondescript. Its dendritic tree is generally extensive with little convergence. Closer examination reveals features not shared with other rabbit cone bipolar cells (Figures 8 and A3). The dendritic branching of most cone bipolar cells, like that of most neurons, gives rise to daughter branches that are notably smaller in diameter than the parent. The ma2 cells exhibit atypical bifurcations, one branch about the same size as the parent, and the other about the same size as the parent or else much smaller (Figure A3). Another atypical feature is lack of taper between branch points, and even enlargements along the course of a dendritic branch. Dendritic terminals are simple and often singular, unlike the more discrete clusters of appendages borne on the terminal dendrites of ma1 cells.

The axon terminals of ma2 cells are even more anomalous, with extended branching, in the inner portion of S1, of comparatively thick processes of uniform caliber. Axon terminal branching is similar to dendritic branching, giving rise to few daughter branches of similar caliber, and a scattering of much thinner, short branches. Short lobular appendages emerge from the main branches, and most of these are direct downward into S2 (Figure A3). One example of a peripheral ma2 cell was found in an array of three adjacent ma2 cells that form a network of connected axon terminals (Figure 9), suggesting homotypic coupling among ma2 cells.

**FIGURE 9.**
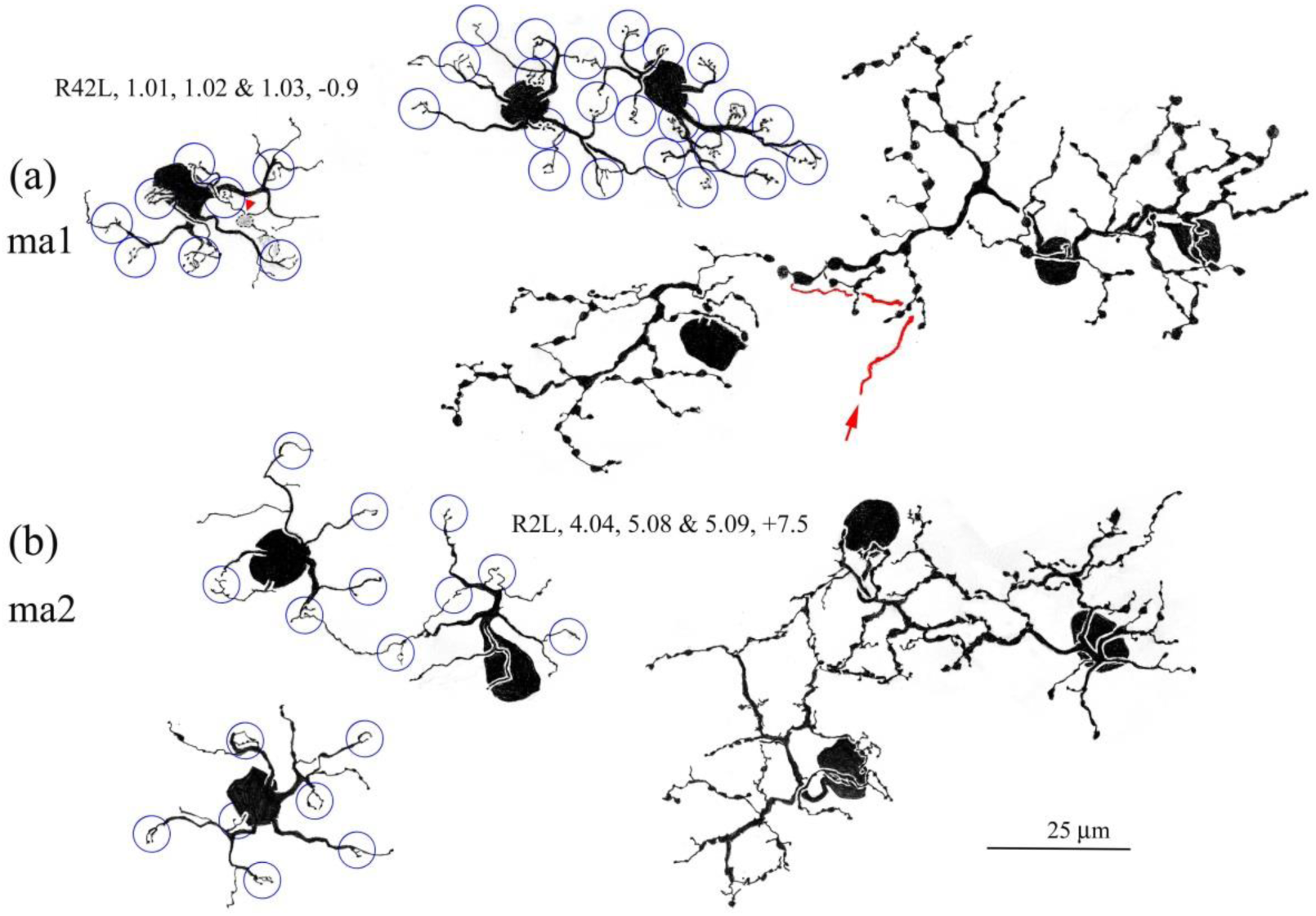
Camera lucida drawings of apparent nearest neighbors among ma1 (a.) and ma2 (b) cells.. Positions of cone photoreceptors (circles) appear to form part of continuous arrays: right pair in (a); top pair in (b). In the example of ma1 cells, the axon terminal of a more distant companion nearly touches the others. A near neighbor of the ma2 pair contributes to a network of axon terminal s suggestive of electrotonic coupling.

#### 3.2.7 wa cone bipolar cells

The wa cone bipolar cells have been described before (Famiglietti, 1981), and in detail elsewhere (Famiglietti, 2008). In 1981, the prediction was made that wa cone bipolar cells were selectively innervated by blue/short wavelength sensitive cones, a fact confirmed by the combination of immunocytochemistry and intracellular staining (Liu & Chiao, 2007). In peripheral, ventral retina, commonly 2-4 cones are innervated, although sometimes only a single cone is contacted (Figures 10, A4). Focal dendritic fields limited to the area directly above the cell body, like some seen in the visual streak (Figure A4), have not been observed In the periphery, perhaps due to an upward shift in the ratio of blue cones to (type a) blue cone bipolar cells.

**FIGURE 10.**
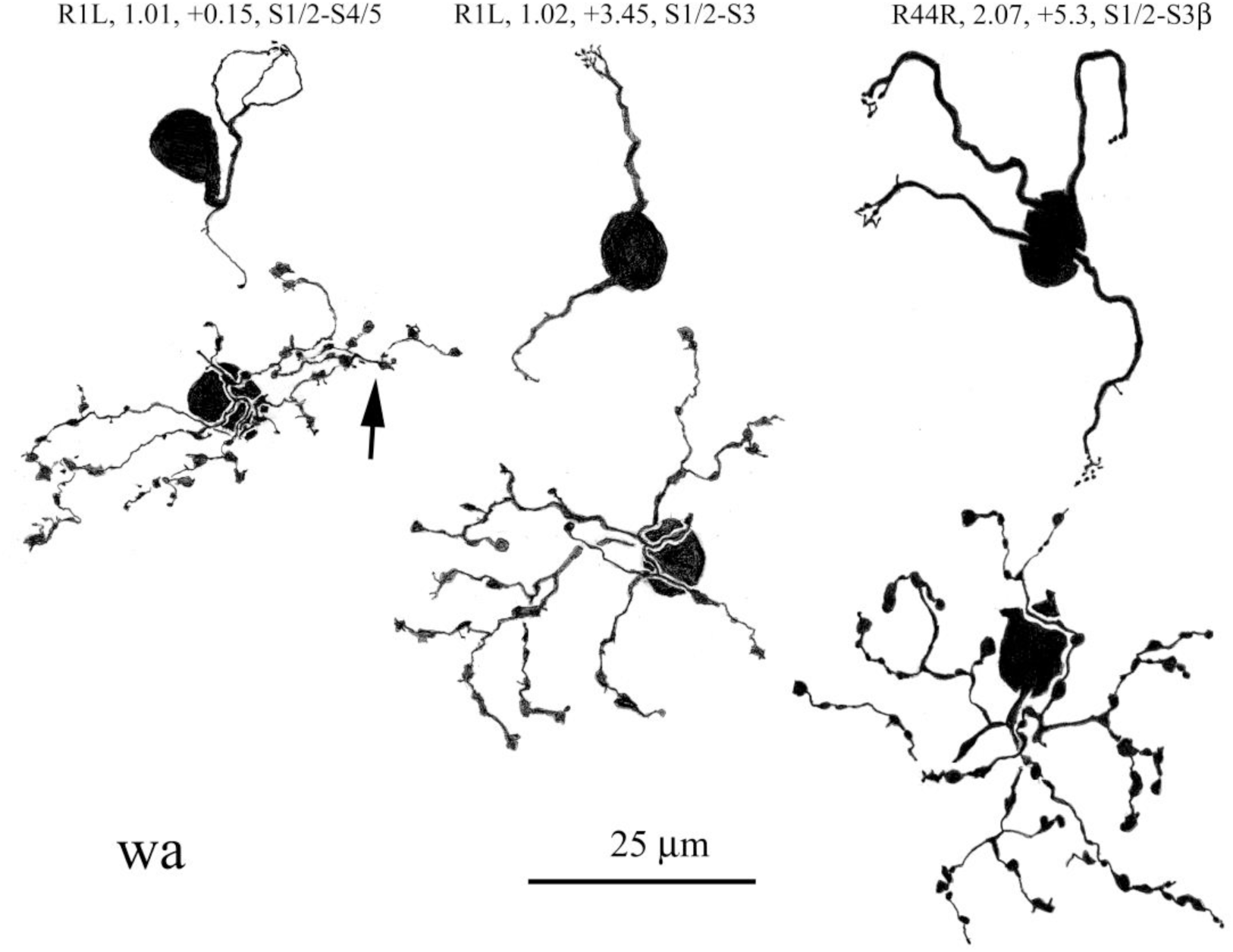
The morphology of wa cone bipolar cells. These select for the minority population of blue cones. Consequently, a number of dendritic branches may converge at a single locus (blue cone pedicle); wa bipolar cells may have only one or two, and sometimes as many as four unbranched dendrites.Axon terminals are slender and beaded. Although their main axon terminal branching is in S1 of the IPL, some branches descend (arrows) to S2, a few to S3, and occasionally to S4 (arrow).

The axon terminal fields of wa cells are not as variable in size as the dendritic fields (Figure 3c,e). They are comparable in size to those of ma1 cells, and are larger in the visual streak (Figure 3e). Their morphology is distinctive, consisting of a low density of slender branches, bearing periodic lobules, giving rise to a beaded appearance (Figures 10, A4). Their stratification is in the main confined to S1, but it is fairly common to see a small number of branches descending into S2, and one or two descending as far as the S3/S4 border (Figure 5). In one example, R1L, 1.01, located in mid-visual streak, one axon terminal branch descends through S4, apparently to the S4/S5 border. Unfortunately, however, no overlap with a fiducial type b starburst amacrine cell has been found in these examples.

### 3.3 Type b cone bipolar cells

In previous work (Famiglietti, 1981), only two kinds of narrow-field bipolar cell: nb1 and nb2, and one kind of wide-field cell: wb, were identified among type b cone bipolar cells. A third kind of narrow-field type b bipolar cell, referred to as “nb3”, with its axon terminal branching in S5, was recognized in our own Golgi-impregnated and immunocytochemical material (Famiglietti, 2002; Sharpe et al., 1993), now named “nb5” here. A complete description of type b cone bipolar cells in Golgi preparations of rabbit retina, documented in the present work, has produced more five more types, for a total of nine varieties of type b bipolar cells (Figure 1). The classification of type b cells has proved far more difficult than such efforts in the case of type a cells (Bordt et al., 2019; Famiglietti, 1981), requiring a three-fold larger sample than that supporting the classification of type a cells.

#### 3.3.1 nb1 cone bipolar cells

As noted above, nb1 and nb2 cells were initially defined in conjunction with na1 and na2 cells. The nb1 cells are the most distinctive of nb cells. They have a radiating array of moderately thick dendrites terminating in a compact cluster of 2-6 digitiform appendages that rise toward (and presumably into) the cone pedicles (Figure 11a,b,h,q). Terminal dendrites are often “zero-order” (unbranched), and are rarely third-order dendrites. These appear to contact all or nearly all cones in the field, amounting to 8-12 cones, whether in the visual streak or in the retinal periphery (Figure A5). As a consequence of their extended, radiating dendritic branching, they have the largest dendritic fields of narrow-field, type b cone bipolar cells (Figure 3d).

**FIGURE 11.**
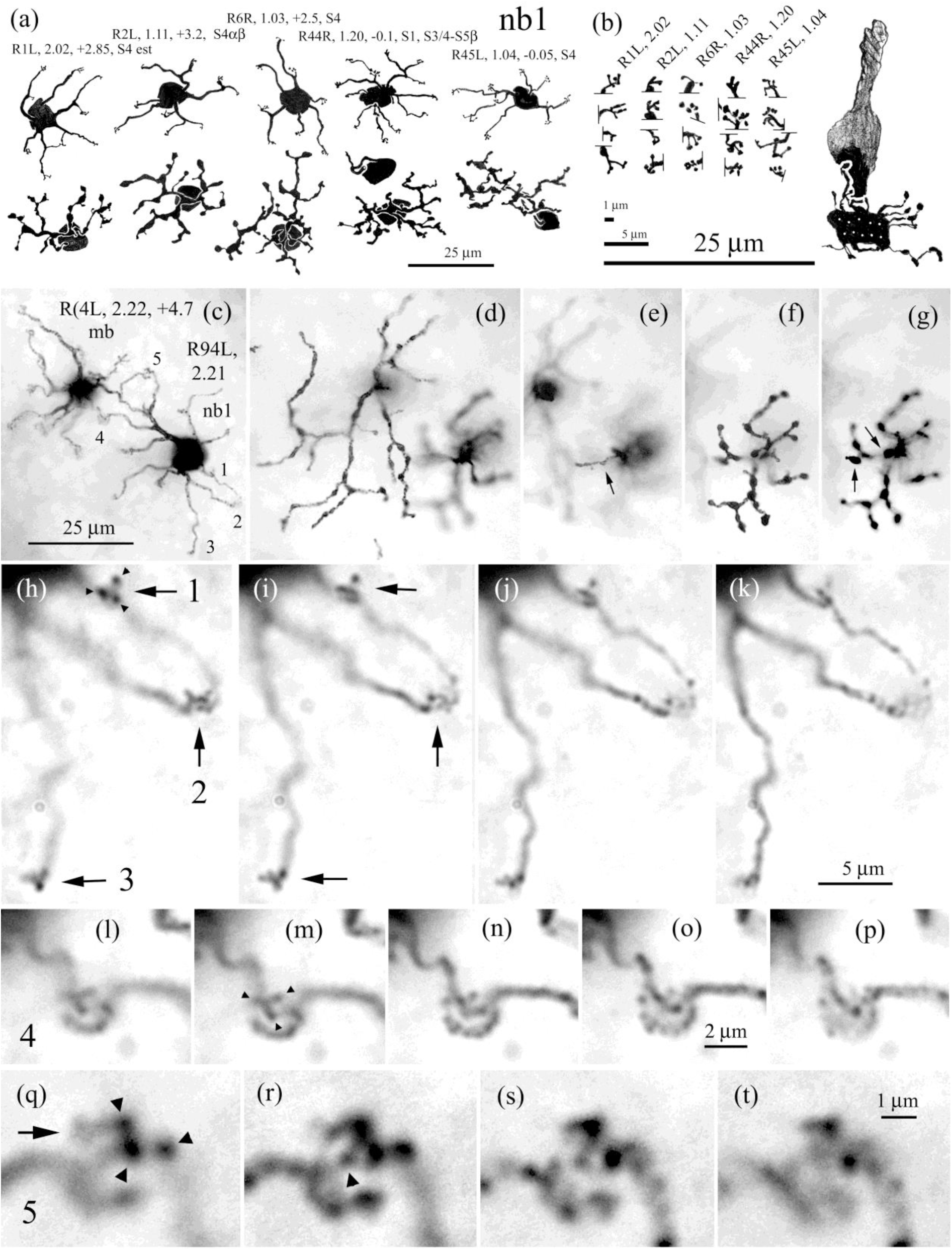
nb1 cone bipolar cells, and their dendritic terminal appendages. (a) nb1 bipolar cells from five different retinas, in and near the visual streak, easily recognized by the relatively uniform radiating dendritic branching pattern, with minimal convergence on single cones, and with 7 to 14 distal dendrites terminating compact clusters of digitiform appendages. Axon terminals are comparatively robust, consisting of branching, lobulated, terminal branches, sometimes scalloped and contour. (b) the terminal clusters of digitiform appendages from the cells in (a). These take various forms, consisting of 2 to 6 short spinous extensions. Cone drawn scale with white dots in cone pedicle, representing estimated 10 ribbon synapses, showing that one pedicle can accommodate two or more bipolar cell terminal branches of these digitiform clusters. (c) photomicrographs of nb1 and mb cone bipolar cells, their dendrites converging on three cones; two (4 and 5) are shown at higher magnification below [(l)-(p) and (q)-(t)]. Three additional terminals of the nb1 cell (1-3) are viewed in (h)-(k). (d) Focus on the mb axon terminal at the S3/S4 border of the IPL. (e) nb1 axon terminal branch in S1 (arrow). (f) Focus on the main portion of nb1 axon terminal in S4. (g) Deeper focus on lobular projections (arrows) near the S4/S5 border. (h)-(k) 0.5 µm step-focusing on vertically projecting digitiform appendages. At the highest level, (h), the apical termini of tri-digital clusters #1 and #2 (arrowheads) are in focus, presumably within invaginations of the cone pedicles. (l)-(p) 0.5 µm step-focusing through cluster #4. (m) Focus on tri-digital cluster from right-hand (nb1) branch. (o) & (p) Single terminal appendages of mb cell, 1-1.5 µm nearer in depth to dendritic branch of origin. (q)-(t) 0.5 µm steps, focusing on cluster #5. (q) Focus the highest level, on tips of tri-digital array (arrowheads), belonging to nb1 cell. Arrow indicates a fourth, more faintly impregnated digital terminal. (r)-(s) deeper focus on termini of both mb and nb1 cells

The nb1 cells have robust axon terminals, bearing bulbous terminal appendages and regions of irregular, varicose thickening (Figures 11a,f,g, A5). They are extensive, rather than compact, and are overall narrowly stratified mainly in S4 of sublamina b, although in some retinas, lobules may rise, or more often descend, from the main level of branching. In a typical example, overlapping the dendritic branching of a BS1 (ON-OFF DS) ganglion cell, the nb1 axon terminal is mostly in contact with the underside of its dendrites in sublamina b (Figure 12a,i,j). On average, their peak stratification lies in the inner portion of S4, just below the middle of the starburst band in sublamina b (Figure 13). In addition, about 1/3 of the present sample of nb1 cells has one or two axon terminal branches in S1α (Figures 11a, e, 13 and A5). Such branches apparently have a functional role (see Discussion).

**FIGURE 12.**
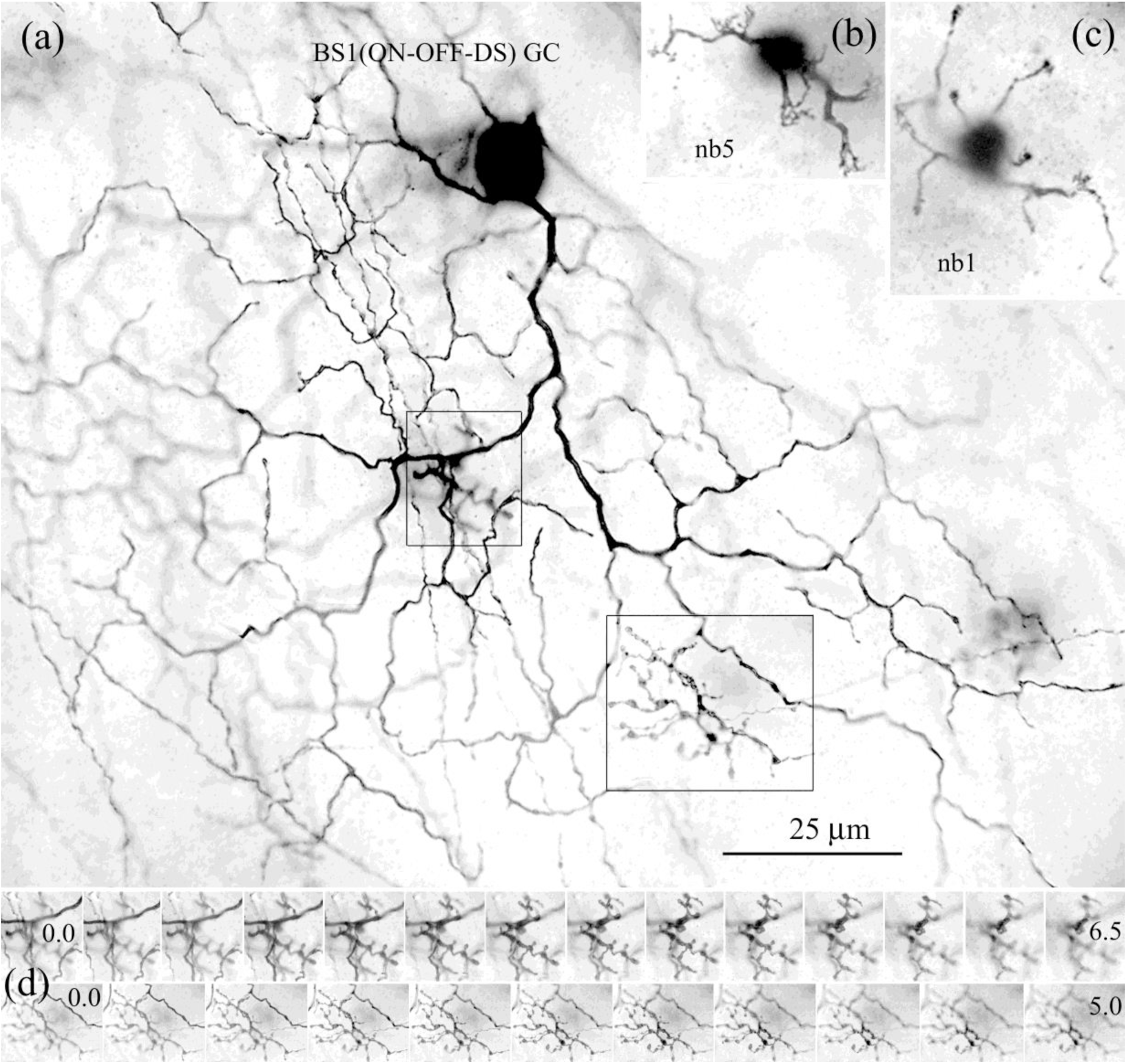
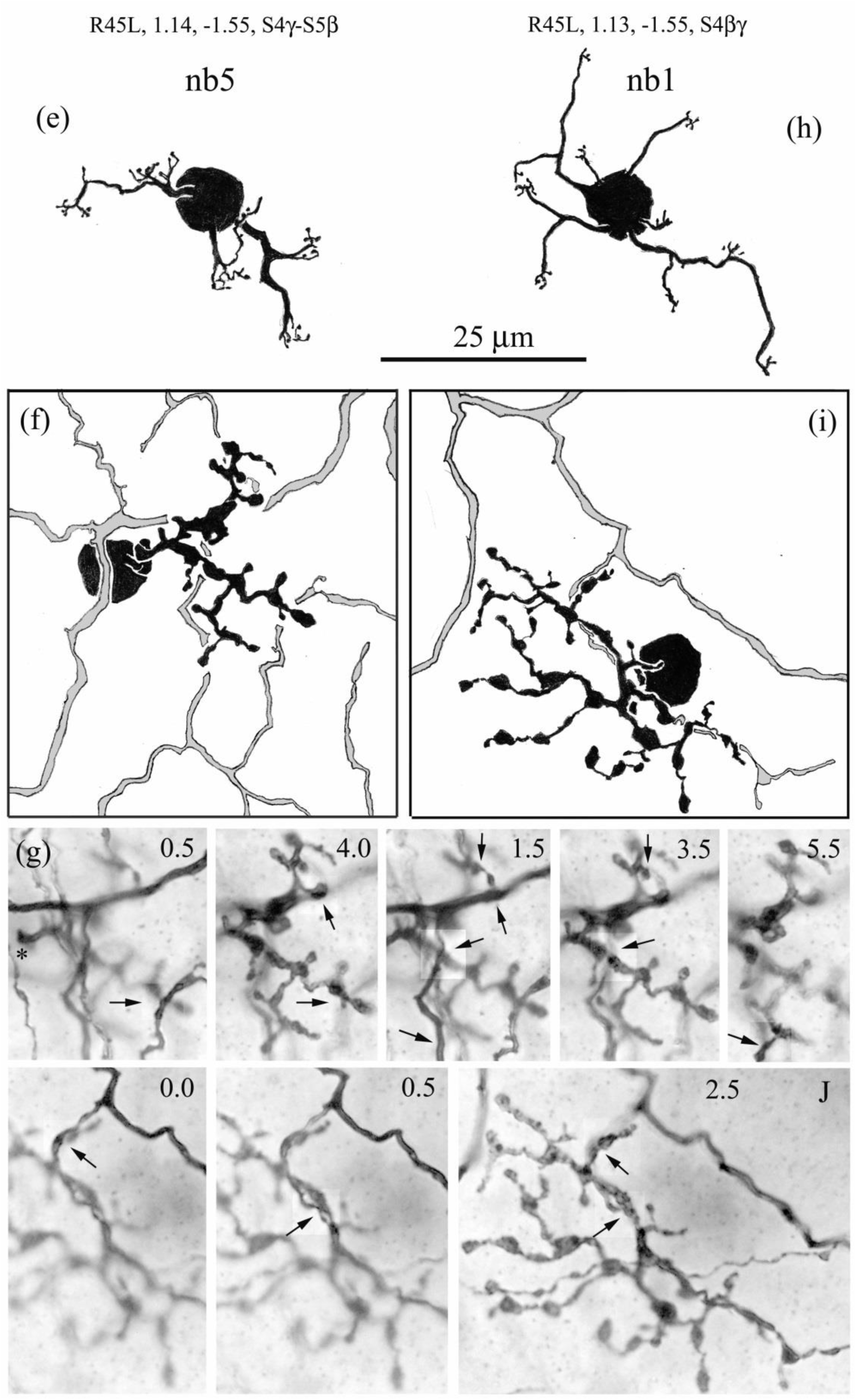
Photomicrographs and cameral lucida drawings of nb1 and nb5 cone bipolar cells with a fiducial BS1 (ON-OFF DS) ganglion cell. (a) dendritic tree of BS1 ganglion cell with overlapping bipolar cells (boxes); focus on branching in sublamina b. (b) and (e) dendritic tree of nb5 cell; note flattened arrays of terminal appendages. (c) and (h) dendritic tree of nb1 cell; note compact clusters of digitiform terminal appendages. (d) optical sections, 0.5 µm stepping, of nb5 axon terminal (14), and of more narrowly stratified nb1 axon terminal (11). (f) and (g) nb5 axon terminal; at five crossing points, laminar distance from BS1 fiducial dendrites to axon terminal varies from 2.0 to 4.0 µm deeper, in the direction of the ganglion cell layer; arrows point to comparable locations of axo-dendritic crossing in two focal planes. (i) and (j) nb1 axon terminal; laminar distance to BS1 fiduacial dendrites varies from 0.0 µm (co-stratified) to 2.0 µm deeper; arrow as in (g). Calibration in (a) applies to (a) – (c); calibration in (h) applies to (e) – (j).

**FIGURE 13.**
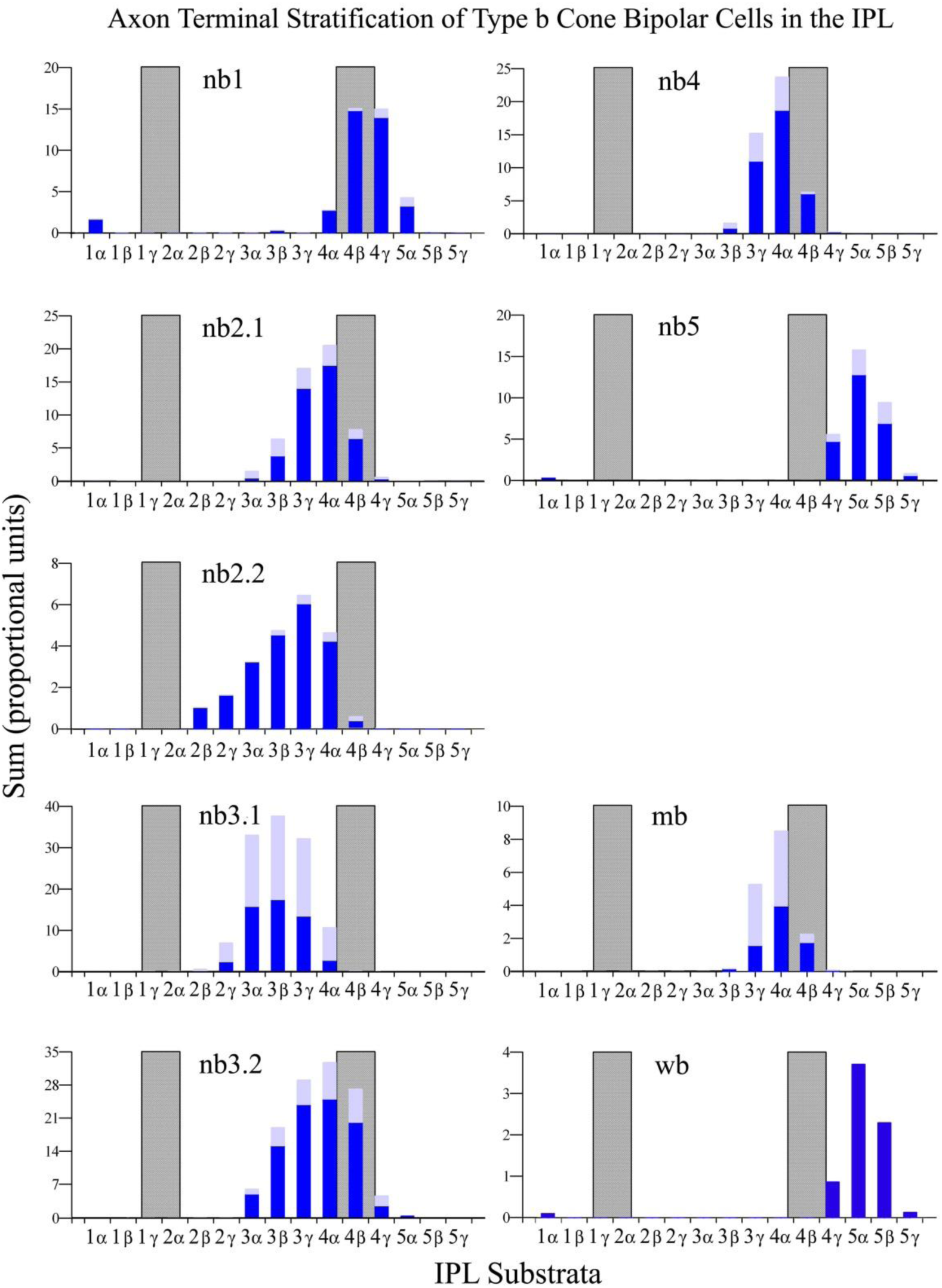
Axon terminal stratification of nine kinds of type b cone bipolar cell in rabbit retina (as for type a bipolar cells in Figure 5). Stacked bars represent sums of determinations of stratification for individual cells in each substratum with respect to fiducial SAb cells (dark bars), or estimates using other fiducials (light bars), depicted against the SAb band of dendrites (gray bar centered on S4β). Medium and wide field cells (mb and wb) have the narrowest stratification. Note that nb1, nb5, and wb bipolar cells may contribute small branches to S1α, although this is far more common for nb1 cells.

#### 3.3.2 nb2 cone bipolar cells

Two features, formerly cited as characterizing the dendritic trees of nb2 cone bipolar cells (Famiglietti, 1981), are: 1) a multitude of digitiform dendritic terminal appendages, and 2) convergence of dendritic terminal branches at the bases of cones. While convergence may be a prominent feature in some nb2 cells, further study reveals that the more conspicuous feature: the presence of many digitiform appendages justifies the subdivision of nb2 cells into two types: nb2.1 and nb2.2, according to the prevalence of each of the two features (Figure 14).

**FIGURE 14.**
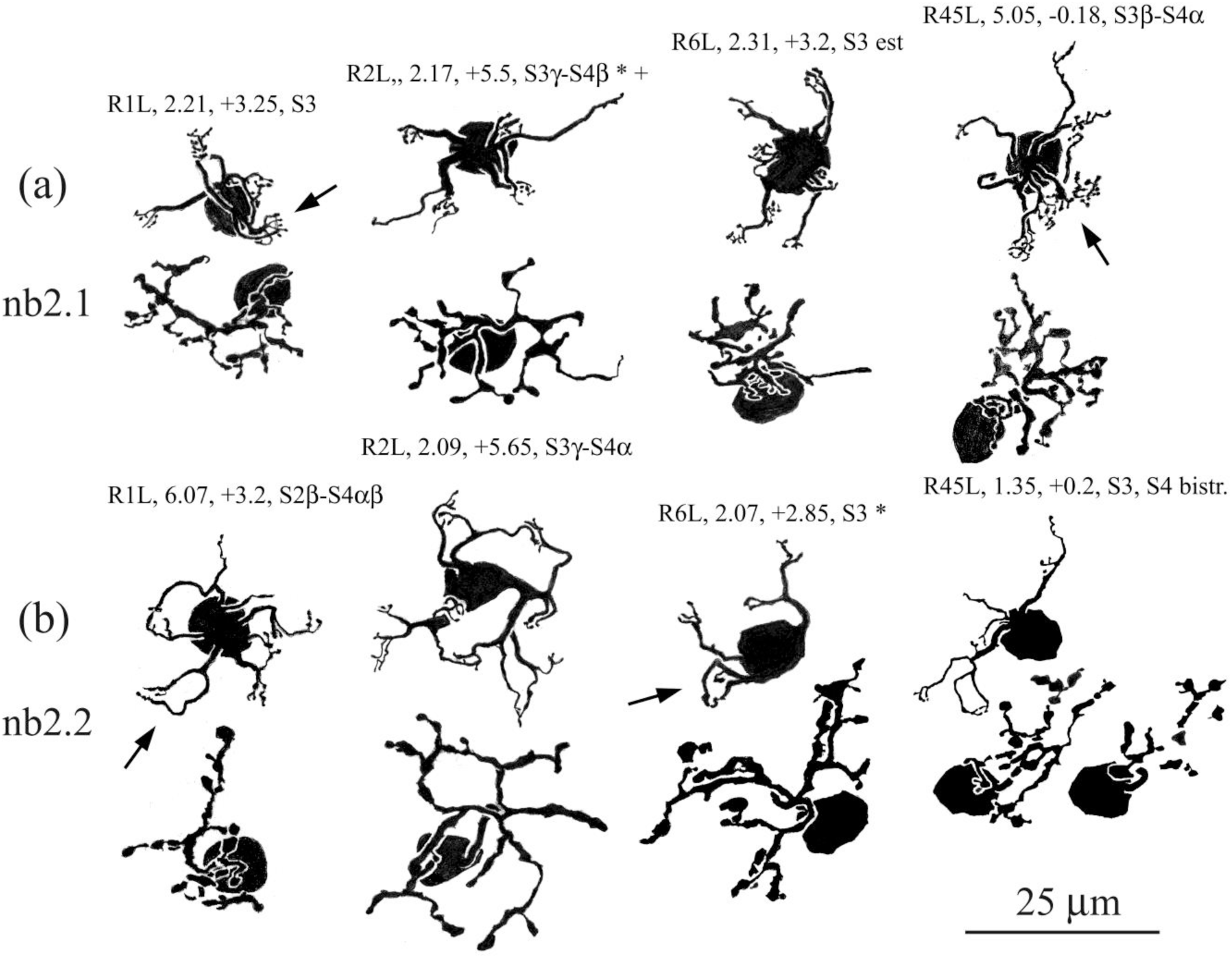
Camera lucida drawings, pairing nb2.1 and nb2.2 bipolar cells in the same retinas (above and below) at similar locations. The nb2.1 cells are characterized by relatively short, direct, and robust dendrites, often converging, giving rise to polydactylic clusters of terminal appendages (arrows). In contrast, the dendrites of nb2.2 cells typically have more meandering, curved trajectory, also commonly convergent, but giving rise to simple arrays of terminal appendages. Often, these curving and convergent, relatively robust terminal dendrites join to form a “claw - like” terminus (arrows).

#### nb2.1 and nb2.2 cone bipolar cells

The morphological feature characteristic of nb2.1 cells is a tight cluster of many digitiform terminal appendages, often borne by a single stout terminal dendrite, but occasionally the result of two dendrites converging (Figures 15a, A6). Usually the cluster of terminal appendages occurs at end of a stout branch that emerges proximal to the cell body with other branches in a radiating fashion. In a variety of examples, this profusion of terminal appendages appears sufficient to innervate the majority of ribbon synapses in a single cone (Figure 15b). Typically, nb2.1 cells have small dendritic fields, with terminal clusters appendages contacting 3 to 5 cones. These appear to be nearly all the cones in proximity to the cell body, at least in locations proximal to the visual streak (Figures 15a, A6).

**FIGURE 15.**
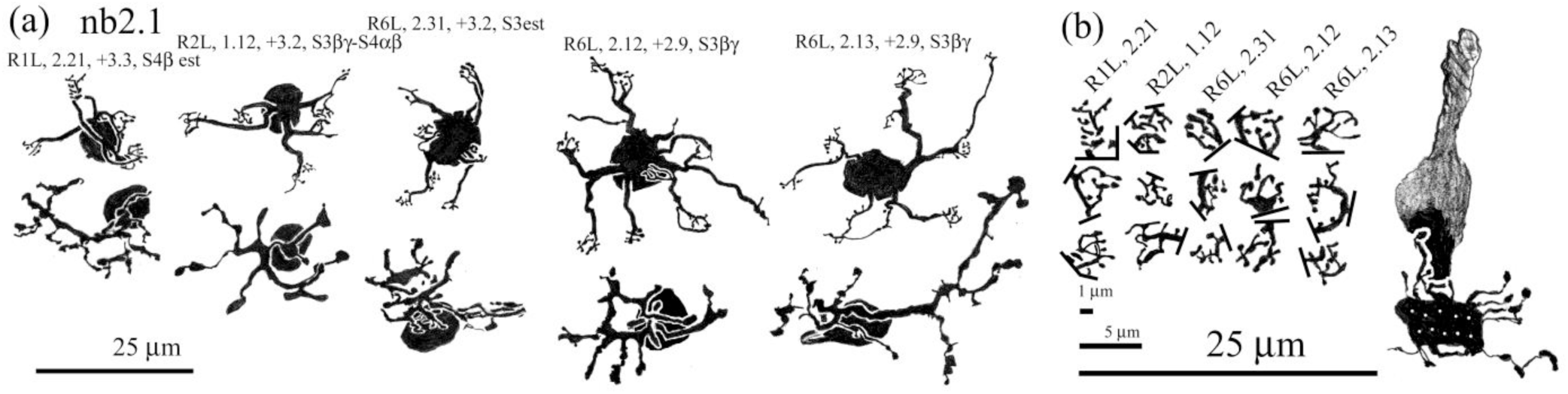
(a) Camera lucida drawings of a variety of nb2.1 bipolar cells from different retinas at about the same *dvs.* (b) Selected terminal polydactylic arrays from each cell are illustrated, with a cone photoreceptor, showing that each could occupy more than half the ribbon related sites in a single cone.

Distinguished from nb2.1 cells by the paucity of their digitiform dendritic appendages and their propensity for dendritic convergence and recurving dendritic branching, nb2.2 cells were identified in seven different retinas (Figures 14b and A7). In addition to dendritic convergence, a related morphological characteristic of nb2.2 dendrites is the formation of “claw-like” dendritic terminal configurations: two thicker short terminal dendritic branches enclosing a few digitiform appendages (arrows in Figure A7).

The nb2.2 bipolar cells are not clearly distinguishable from nb2.1 cells in their axon terminal morphology viewed in two dimensions. Their stratification is broader, however, and shifted towards sublamina a (Figure 13). Surprisingly, of the eight nb2.2 cells with adequate fiducial markers, four extended some axon terminal processes into S2.

#### nb2.1 cone bipolar cell subtypes?

One problem that arises in formulating a clear identity for nb2.1 cells, based primarily on the morphology of their dendritic trees, is that a small number of examples and similar dendritic terminal clusters have dendritic fields twice the size of typical cells at the same retinal location, with more widely spaced cone contacts. This was observed at least three different retinas (Figures 16a, b). As a rule, for narrow field cone bipolar cells, there is a good correspondence between the sizes of the dendritic and axonal trees in the plane of the retina, particularly for type b cells (Figure 3b). It is not always the case for nb2.1 cells with large dendritic trees that their axon terminal fields are also comparably large (Figure 16b).

**FIGURE 16.**
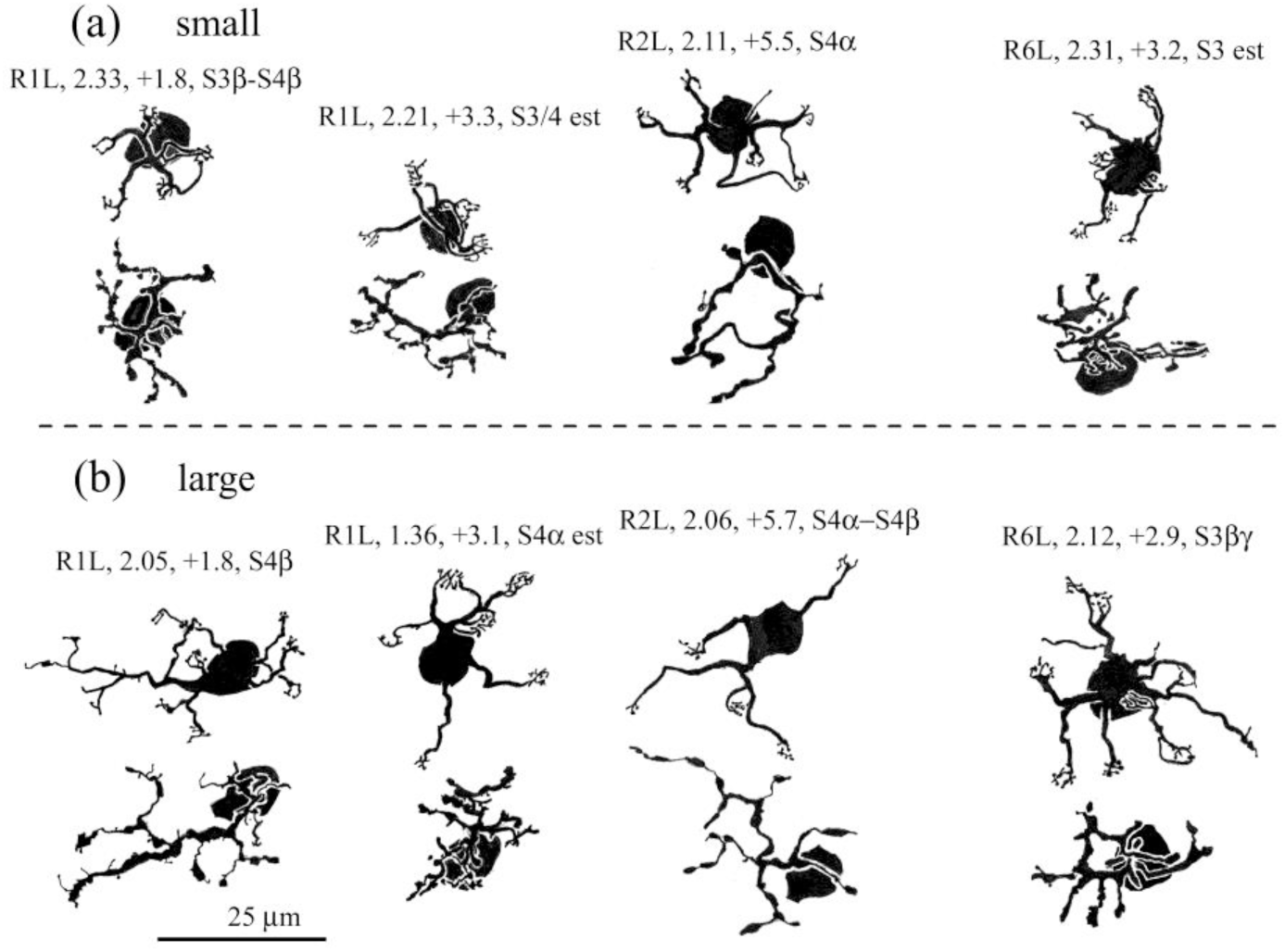
Camera lucida drawings of small (a) and large (b) varieties of nb2.1 cell, taken from the same retina at about the same *dvs* (above and below). (a) The small examples appear to contact all the cones near the cell body. (b) The large examples, in contrast, may bypass some nearby cones, resulting in more extensive dendritic trees.

When accepting cells with large and small dendritic trees as examples of the same type in this case, it is helpful to consider cell R2L, 2.06 (Figure 16b) together with the closely adjacent cell R2L, 2.07 (Figure 17). The dendritic trees of the two cells, in contrast to their axonal arborizations, are highly overlapped. The dendrites of each cell contact cones overlying the cell body of the other. Yet the distribution of cones overlying both forms a more or less regular array. Insofar as these two nb2.1 cell bodies are likely closer together than average for this retinal location, they may provide an instance highlighting the different sampling problems that confront bipolar cells in the outer plexiform layer (OPL), where the photoreceptor targets are discrete, discontinuous, and sometimes relatively distant, and in the IPL where postsynaptic targets are more proximate and continuous.

**FIGURE 17.**
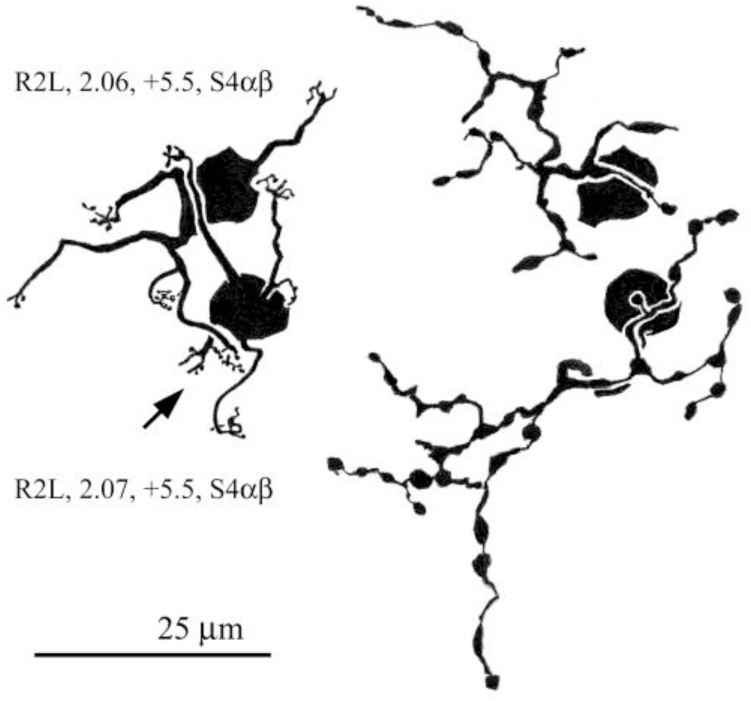
Cell R2L, 2.06 from Figure 14 and it’s very near neighbor 2.07 illustrate the fact that the “larger” nb2.1 cell can arise from mismatch of cone photoreceptor and nb2.1 bipolar cell body arrays, such that a putative competition for the more regular array of cones can result in deep extension of the dendrites of inordinately proximate nb2.1 cell bodies to reach cones overlying the neighboring bipolar cell. Where convergence of dendrites occurs between neighbors (arrow), the more proximate one predominates here. Note that there is no overlap of axon terminal arbors, directed by a different form of competition in the IPL.

A second problem, pertaining to axon terminal stratification, and independent of the first, also requires consideration of another possible subdivision of nb2.1 cone bipolar cells into two types. This problem illustrates a more general and unexpected issue of some variability in the axon terminal stratification of rabbit cone bipolar cells.

In the sample of 79 nb2.1 cells, it became evident that most were broadly stratified in the IPL, while a smaller number exhibited narrow stratification at the margin of the same range. In order to resolve this problem with a sufficient degree of confidence, it was necessary to take into account a degree of individual variation between experimental subjects, and to restrict the subsample to examples with overlapping fiducial cells. Thus, seven cells of the broadly stratified group were compared with eight cells of the narrowly stratified group, all from one retina, R2L (Figure 18a, b). The two groups, nb2.1.1 and nb2.1.2, were indistinguishable in terms of their dendritic morphology. The extent of axon terminal stratification, summed over the set of cells, for each substratum, in the case of the broadly stratified set, nb2.1.1, extended from S3α barely to S4γ, centered on the S3/S4 border (Figure 18c). For the narrowly stratified set, nb2.1.2, stratification extended from S3γ to S4β and was centered on S4α.

**FIGURE 18.**
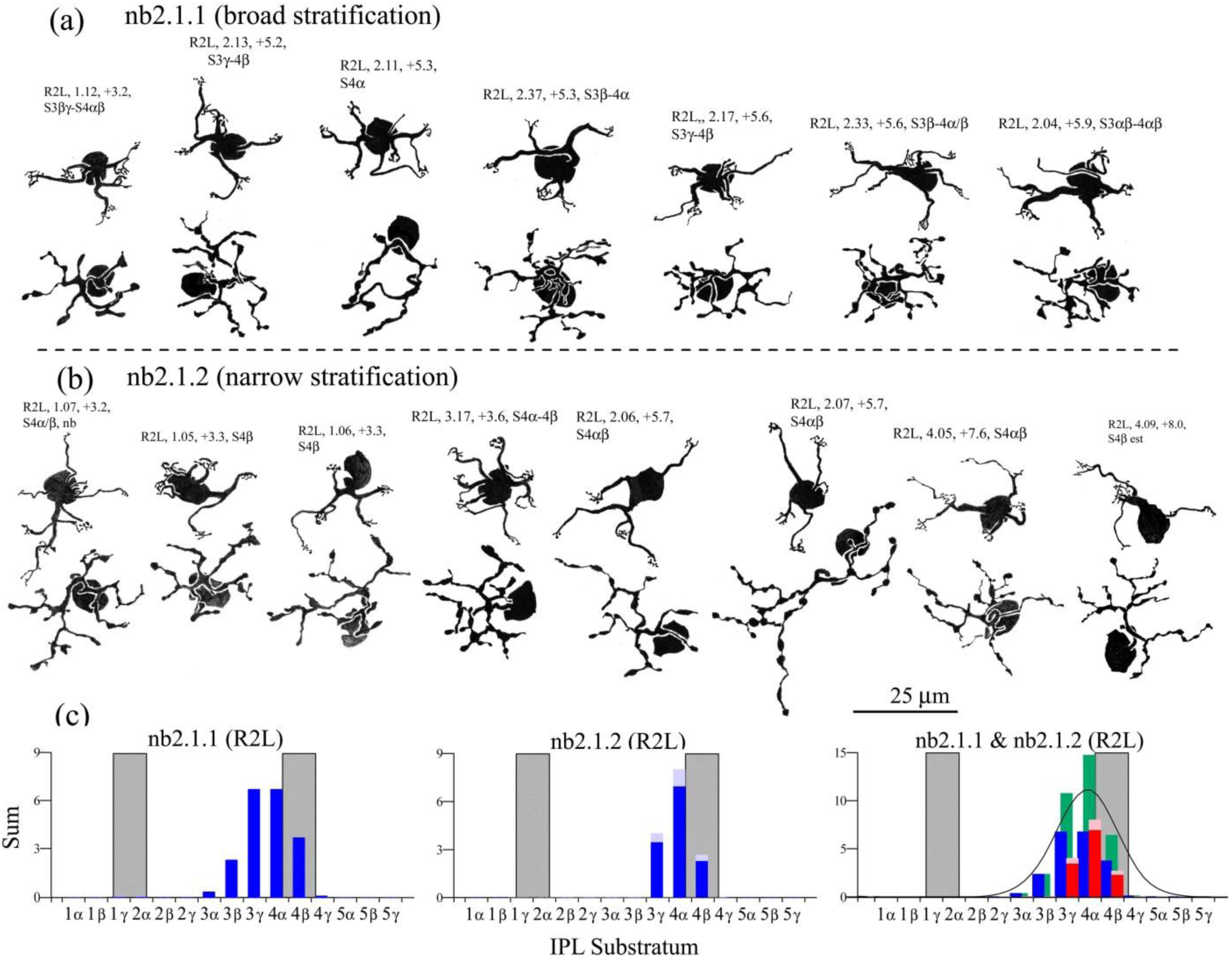
Comparison of possible (a) nb2.1.1 and (b) nb2.1.2 subtypes, differing in axon terminal stratification, all in the same retina, to control for individual variation. (c) Graphs show that the latter are encompassed in the broader stratification of the former. When the two (blue and red in right-hand graph) are summed (green) and fitted by a DWLS function, the curve is symmetrical, suggesting that the null hypothesis should not be rejected.

When the two are compared in combination (green bars, Figure 18c), it is clear that, in their axon terminal stratification, narrowly stratified nb2.1.2 cells do not extend outside the range of the broadly stratified nb2.1.1 cells. While the narrow stratification of nb2.1.2 cells could reflect more selective innervation of a narrowly stratified ganglion cell type, there are so many types of ganglion cell branching at or very near this level (Famiglietti, 2020), that such substratification would be uncertain grounds for distinguishing between nb2.1.1 and nb2.1.2 cells, particularly in view of the significant mismatch in numbers between the relatively few types of bipolar cell and the many types of ganglion cell requiring bipolar cell input.

In addition to differences in axon terminal stratification, nb2.1.1 and nb2.1.2 cells differ in average axon terminal field size. On average, narrowly stratified nb2.1.2 cells are 30% larger than nb2.1.1 cells (data not shown). The nb2.1 cells, numerically weighted by a preponderance of nb2.1.1 cells in the sample (84%), have the smallest axon terminal diameters of all type b cells (Figure 3f).

#### 3.3.3 nb3 cone bipolar cells

In two prior publications (Famiglietti, 2002; Sharpe et al., 1993), a third narrow-field, type b cone bipolar cell was referred to as “na3”, although it was not described in detail. That cone bipolar cell, now well characterized here, is henceforth termed “na5” (see below).

The nb3 cone bipolar cells of the present study have an easily recognized, simple dendritic branching pattern, with moderately robust proximal branches, shorter tapering lengths, and few orders of branching (Figures 19 & 20). Occasional solitary spines are observed along the course of dendritic branches. Typically, about half of their terminal branches are adorned with appendages (Figure 19a, b). On average, their dendritic field diameters are the smallest of type b cone bipolar cells (Figure 3e). In view of the spare architecture of their dendritic trees, a large sample of nb3 cells was studied in all retinas to ensure that these were not simply under-impregnated bipolar cells belonging to one or more other types. The morphology of nb3 axon terminals is generally that of beaded processes (Figures A8 and A9), relatively compact in the retinal plane, and of intermediate size among nb cone bipolar cells (Figure 3f).

**FIGURE 19.**
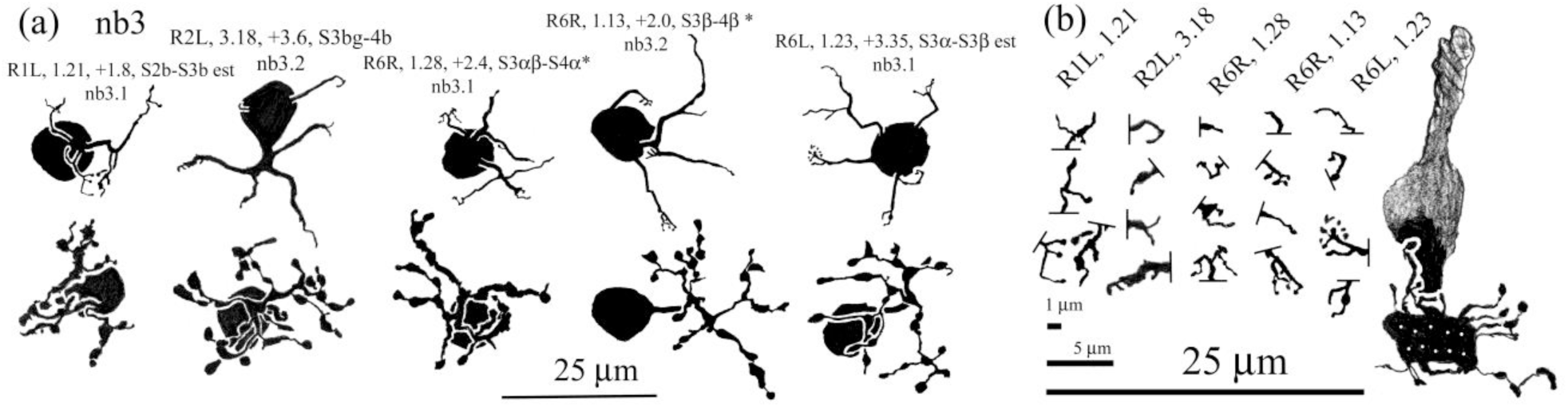
(a) Camera lucida drawings of nb3 cone bipolar cells from several different retinas. Dendritic branching is simple, and dendrites bear few appendages. Axon terminal branches are typically beaded. (b) dendritic terminals of these cells are usually simple and even solitary

A total of 88 nb3 cells were studied. Of these 43 were accompanied by satisfactory fiducial cells for the analysis of axon terminal stratification, and in a further 22 cases good estimates could be made. About 15% exhibited a somewhat bistratified distribution of axon terminal processes. The depth of stratification of the axon terminals of individual cells was not exceptionally broad. It was problematic, however, to observe the number of levels over which whole sample ranged.

The sample of nb3 cells could be provisionally divided in two (Figure 20a, b): those the axon terminals of which touched or breached the a/b sublaminar border into S2, and were otherwise confined to S3 (nb3.1), and those that branched from inner S3, across the S3/4 border (nb3.2), to at least partially overlap type b starburst amacrine cells (Figure 21a-d). The sample obtained with accompanying fiducial cells is unequal for the two groups, but they were also examined retina by retina, and consistent results were obtained in that analysis (Figure 21e, f). When the axon terminal stratification of the sample of nb3.1 and nb3.2 cone bipolar cells from all retinas is compared, two distinct, partly overlapping distributions are seen (red and blue bars, Figure 21g). When the two are summed, however, the result appears to be consistent with a normal distribution (green bars, Figure 21g), favoring a unitary grouping of nb3.1 and nb3.2 cells.

**FIGURE 20.**
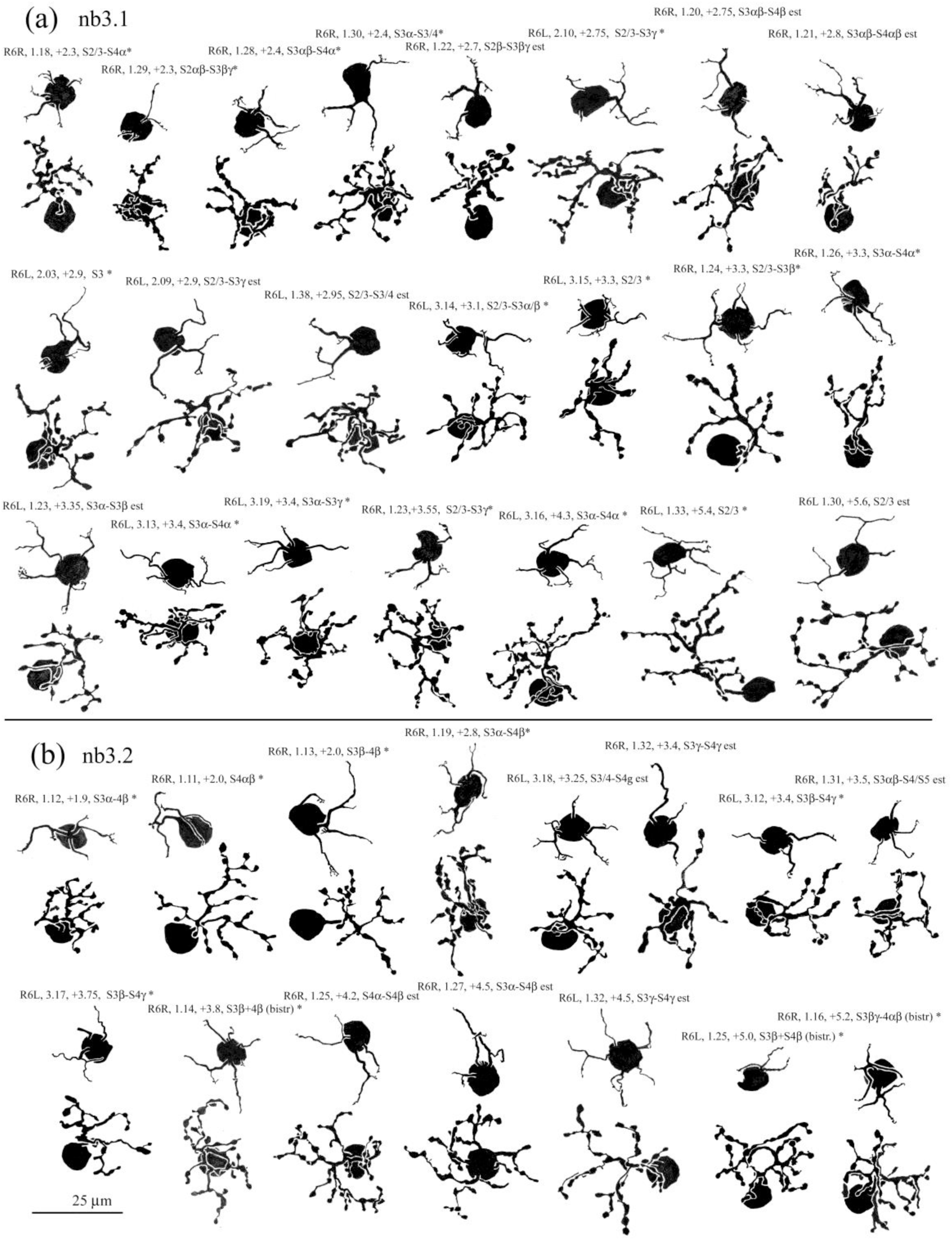
Camera lucida drawings of nb3 cells from a single animal, divided into two subtypes, nb3.1 (a) and nb3.2 (a). The dendrites are moderately thick, uniformly bereft of dendritic appendages. Their axon terminals are lobulated and of moderate complexity. In flat view, as seen here, nb3.1 and nb3.2 cells are indistinguishable.

**FIGURE 21.**
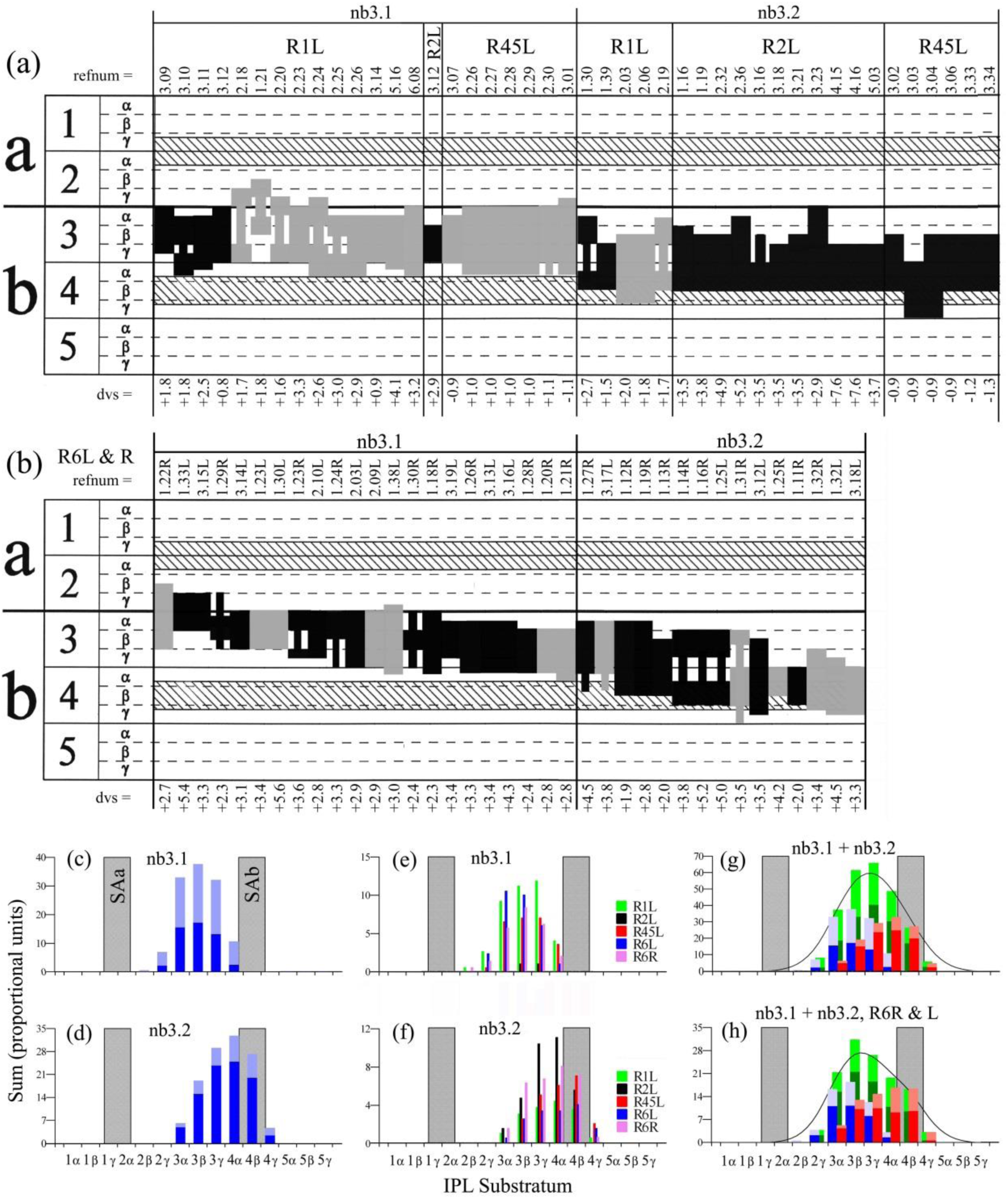
Analysis of the axon terminal stratification of nb3 cells in the IPL. (a) Examples of nb3.1 (left) and nb3.2 (right) cells from three retinas are registered in a stratification diagram, either accompanied by fiducial type b starburst amacrine cells (SAb) (black) or by other fiducials (gray). The stratification covers a broad range; nb3 cells are divided into those making no contact with SAb cells (nb3.1), and those that do make contact (nb3.2). (b) Thirty-seven nb3 cells from a single animal arranged in descending order of stratification level. (c) and (d). Summary graphs show only partial overlap of stratification: S2γ –S4α (nb3.1) and S3α – S4γ (nb3.2). (e) and (f) Results are consistent for all retinas studied. (g) When all retinas are summed, the result is fitted by a DWLS curve that resembles a normal distribution, suggesting that nb3.1 and nb3.2 cells are drawn from a single population. (h) When a sum of examples from a single animal is considered, however, the dwls curve is skewed, suggesting that the two populations may be distinct.

With an eye to the possibility that the variation in axon terminal stratification of nb3.1 and nb3.2 cells represents a continuum, rather than a marker of two types, examples restricted to a single pair of retinas of the same animal (R6, Figure 20) were subjected to separate evaluation (Figure 21b, h). Thirty-seven cells were sorted in a sequence representing increasing depth of stratification in the IPL (Figure 21b). Bisecting the sample into “nb3.1” cells branching almost exclusively in S3 (N = 22), and “nb3.2” branching equally in S3 and S4, or largely in S4, and also overlapping type b starburst amacrine cells (N = 15), yielded two partly overlapping distributions (red and blue bars, Figure 21h). To equalize the two groups, the per-substratum counts in respect to the latter were multiplied by 1.47, and these values summed with the R6, nb3.1 values (green bars, Figure 21h). The curve fitted to this sum is skewed, suggesting that nb3.1 cells and nb3.2 cells may belong to separate populations. From a functional standpoint, the range of stratification of nb3.1 cells and nb3.2 cells taken together seems expansive for a single type of cone bipolar cell, when considering potential connectivity with narrowly stratified postsynaptic targets.

#### 3.3.4 nb4 cone bipolar cells

A practiced eye is required to distinguish between nb4 and nb1 cone bipolar cells, and a focus on details is important. The two types have similar extended dendritic branching patterns (Figure 22a and b), and they are about the same size (Figures 3e and 3f). Two features are especially useful in making the distinction. The first of these is the compact cluster of digitiform appendages at terminal dendrites of nb1 cells and described above (Figure 11). Instead, a typical feature of nb4 cell dendrites is the termination of distal dendrites in pairs of short, slender branches that often form small, open rings (arrows, Figure 22a; see also Figure A10).

**FIGURE 22.**
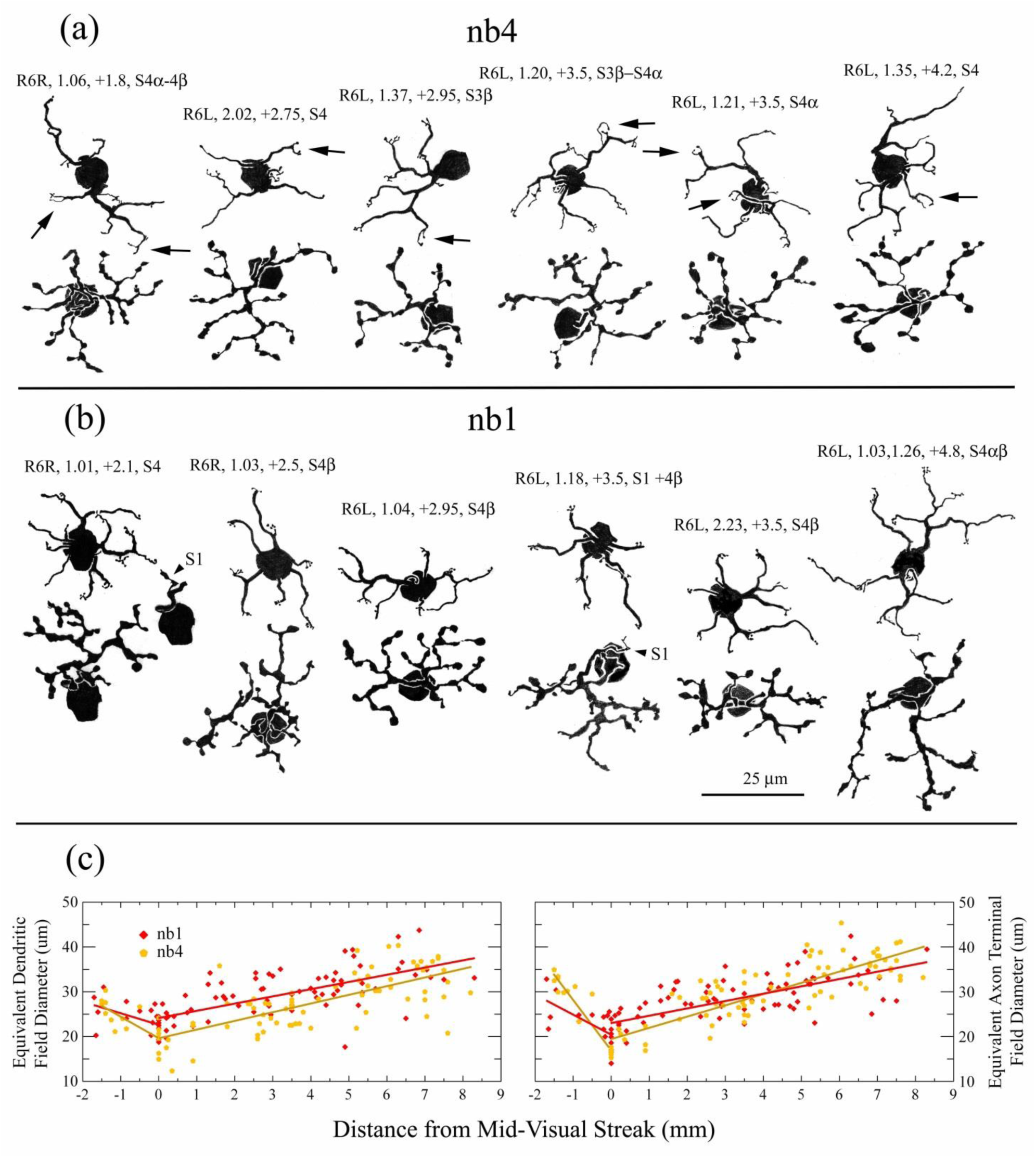
Camera lucida drawings in a morphological comparison, distinguishing between nb4 (a) and nb1 (b) bipolar cells, all from the same animal. Extended dendritic branching patterns are similar, with regular dichotomous branching and some long, unbranched dendrites, mostly of moderate caliber. Conspicuous differences occur at the terminals, however, branches of nb4 cells often ending in small circlets [arrows in (a)], in contrast to the tight clusters of digitiform appendages exhibited by nb1 cells (Figure 11). A consistent difference in the morphology of axon terminals is the characteristic “string of beads” appearance in the case of nb4 cells. S1 axonal branches have not been observed in a sample of more than 50 nb4 cells, whereas about 1/3 of nb1 cells have S1 branches. (c) Quantitative comparison of axon terminal field size supports the distinction of nb4 from nb1 cells: in the visual streak, nb1 cells are larger than nb4 cells, whereas the reverse is true in peripheral retina

The second notable and characteristic difference between nb4 and nb1 cells is a conspicuous feature of nb4 axon terminals, which invariably exhibit a beaded appearance (Figure A10). The nb1 cells do exhibit some beaded segments in their axon terminal branching, which is nevertheless dominated by more robust, lobular processes, sometimes exhibiting scalloped contours (Figure A5). A detailed comparison of the sizes of 90 nb1 and 84 nb4 cells reveals that on average the former have slightly larger dendritic trees at all locations, but more telling, the axon terminal field sizes of nb1 and nb4 have different distributions with respect to retinal location. The nb1 cells are larger in and near the visual streak, whereas nb4 cells are larger in the retinal periphery (Figure 22c).

In a few instances, pairs of nb1 and pairs of nb4 cells have been found in “nearest-neighbor” proximity, usually innervating some of the same cones (Figure 23). In regard to axon terminal proximity, three patterns are observed in the case of nb4 cells: close apposition, bypass with possible contact, and no contact (Figure 23a). Absence of contact between nb1 cells is also seen in one example, but in several others, close contact is observed (e.g. Figure 23b). It is significant that in the more distanced pair, even when no cones are shared, two close contacts are formed, suggesting that nb1 cells may form a continuous network in the IPL (see Discussion).

**FIGURE 23.**
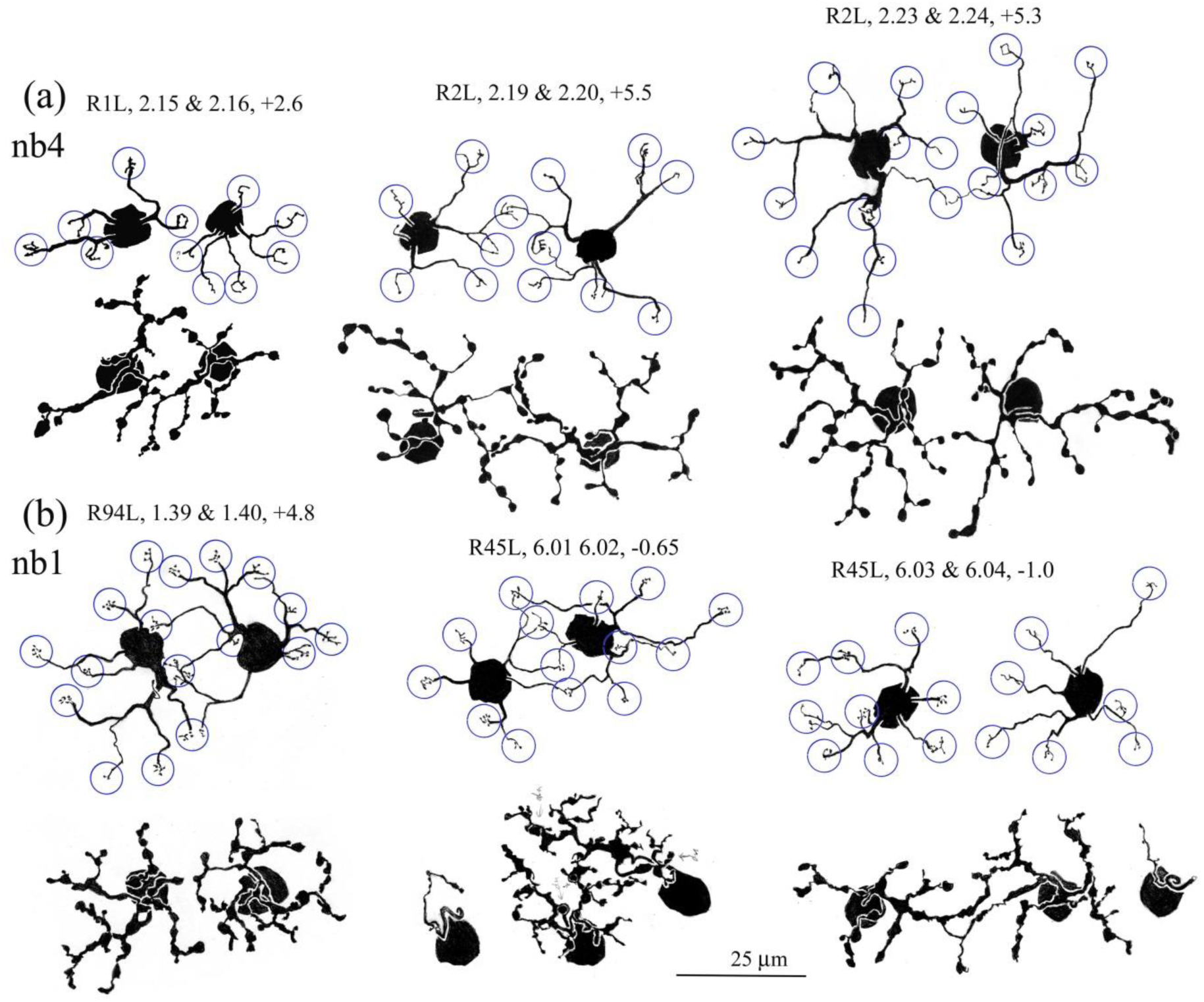
Camera lucida drawings of nearest neighbors among nb4 (a) and nb1 (b) cells, apparently contacting some of the same cones (circles). (a) In the three examples of nb4 pairs, one shows a close contact between axon terminals (arrow); one shows no contact. At a third, close overlap occurs, but there is no clear site of apposition. (b) In three examples of nb1 cells, two with S1 branches, one shows no contact, but the others show close contact, and the right-hand example of somewhat more distant cells exhibits multiple contacts

The axon terminals of nb4 cells are narrowly stratified like those of nb1 cells, but with fewer lobules ascending or descending from the principal plane of branching. The levels of their branching are rather close. The main level for nb4 cells, however, lies near the boundary of S3γ and S4α, whereas that of nb1 cells is at the boundary of S4β and S4γ (Figure 24). Unlike nb1 cells, no nb4 cells in the sample formed axonal branches in S1.

**FIGURE 24.**
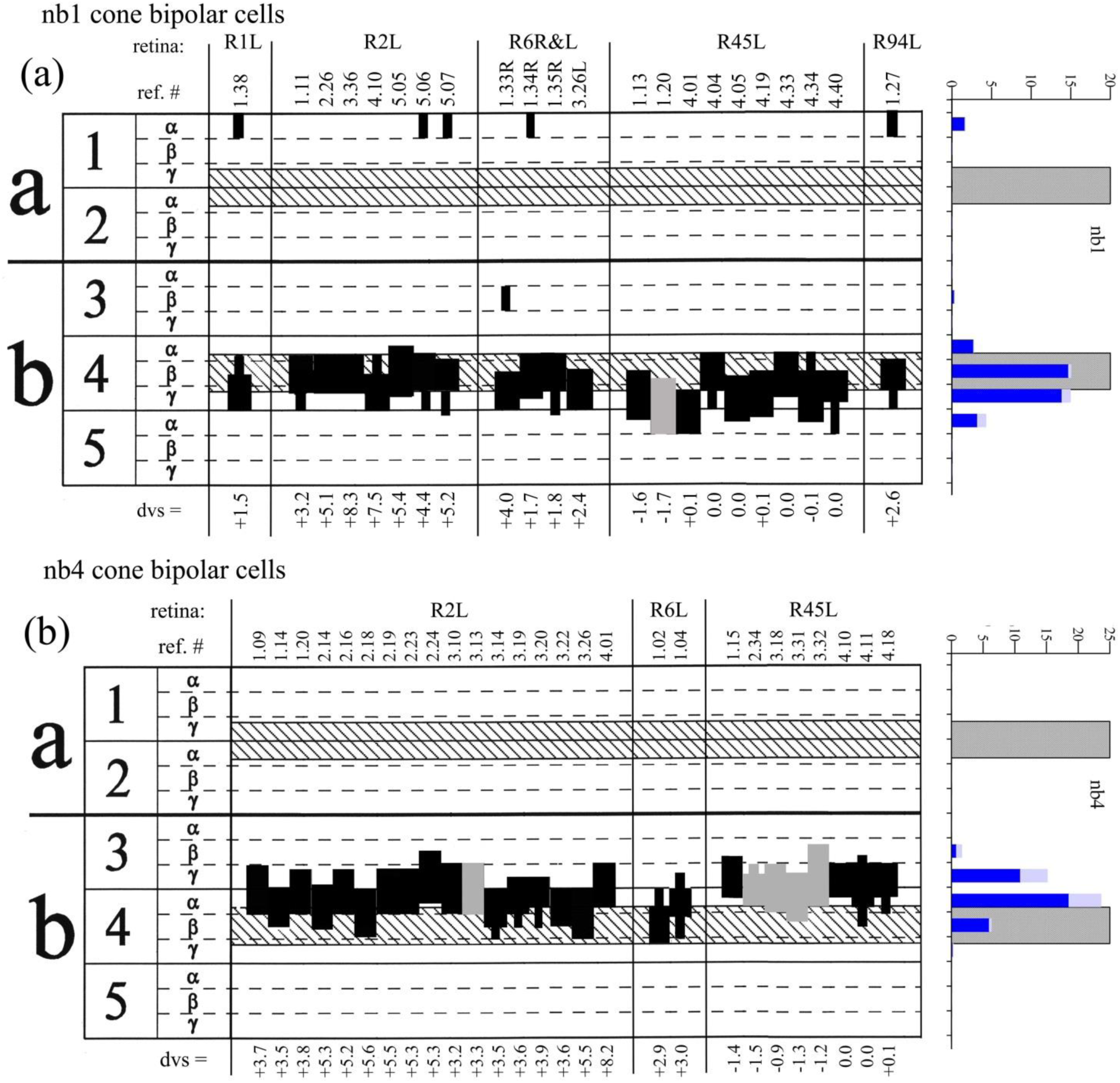
Analysis of axon terminal stratification, comparing nb1 (a) and nb4 (b) cells. Black denotes availability of overlapping SAb fiducial cells, and gray denotes estimates, using other fiducials. Sums at right show nb4 cells branching concentrated at the upper bound of the SAb band, whereas nb1 cells overlap and lie at the lower bound of the SAb band. Note the presence of S1 branches (mostly single and unbranched) of nb1 axons

#### 3.3.5 nb5 cone bipolar cells

The axon terminal fields of nb5 cone bipolar cells are about the same size as those of nb1 and nb4 cells, except in the retinal periphery, where they are somewhat smaller (Figure 3f). The nb5 dendritic trees are comparable in size to those of nb4 cells, and only a little smaller than those of nb1 cells (Figure 3e). The extended branching of their dendritic trees is marked by mostly small, scattered dendritic spines that occur along the lengths of dendritic branches and in loose aggregations at the ends of terminal dendrites (Figures 25 and A11). In some examples, the axon terminals also bear more than the usual, small number of dendritic spines (Figure A11). Otherwise, the axon terminal morphology of nb5 cells does not differ markedly from that of nb1 cells, sharing both scalloped and lobular processes, except that it occurs principally in S5 (Figure 13).

**FIGURE 25.**
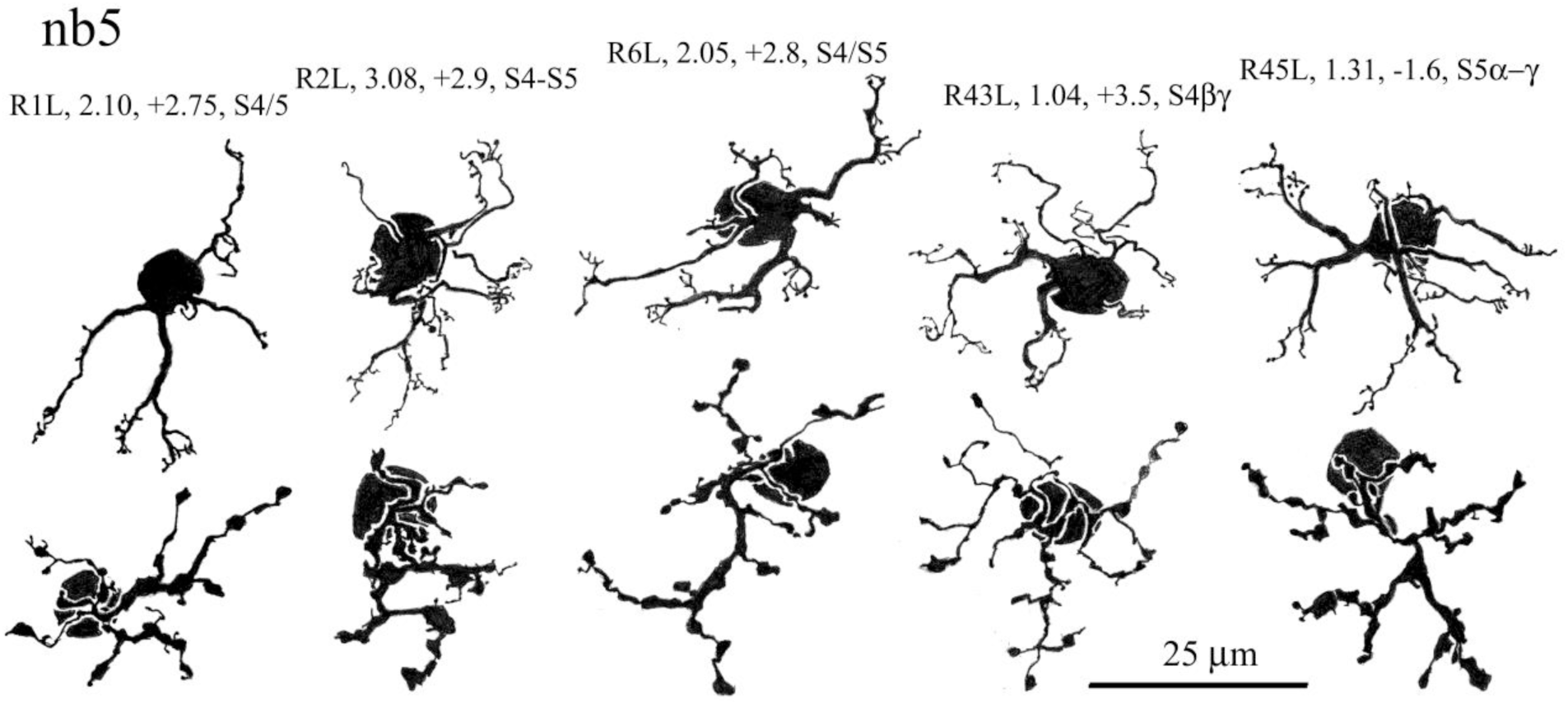
Camera lucida drawings of nb5 cone bipolar cells. Most conspicuous are the slender spines adorning both dendrites (above) and axon terminals (below), although in the case of dendrites, most of the irregular aggregations of spines occur at terminals of extensive and usually robust dendrites. Axon terminals are robust and reminiscent of those belonging to nb1 cells.

A typical example of an nb5 cone bipolar cell is illustrated in the drawings and photomicrographs of Figure 12b, and 12e-g, its axon terminal viewed in serial optical sections with reference to the overlapping fiducial BS1 ganglion cell branches in sublamina b (Figure 12a). At nine crossing points of the axon terminal processes overriding the fiducial BS1 ganglion cell dendrites, the distance between them ranges from 1.5 to 5.0 µm. Thus at no point do they come into contact, and at a distance of 1.5 µm, the axon terminal breaches the S4/S5 boundary by no more than 1.0 µm.

Close examination of axon terminal stratification could be made in 16 Golgi-impregnated nb5 cone bipolar cells from five different retinas with the aid of fiducial cells (Figure 26a). At 49 crossings, the mean distance between them and the middle of S4 is 2.3 ± 1.1 µm, which on average coincides with the middle of S5α. At 6 of 49 crossings the distance measured was 0.0 or 0.5 µm, indicating close contact and potential synaptic contact. The level of axon terminal stratification of nb5 bipolar cells (Figure 26b) positions them favorably to make synaptic contact with large bodied class II and class I (“X” and “Y”) ganglion cells (Figure 26c; cf. Figure 12 in Famiglietti, 2004a).

**FIGURE 26.**
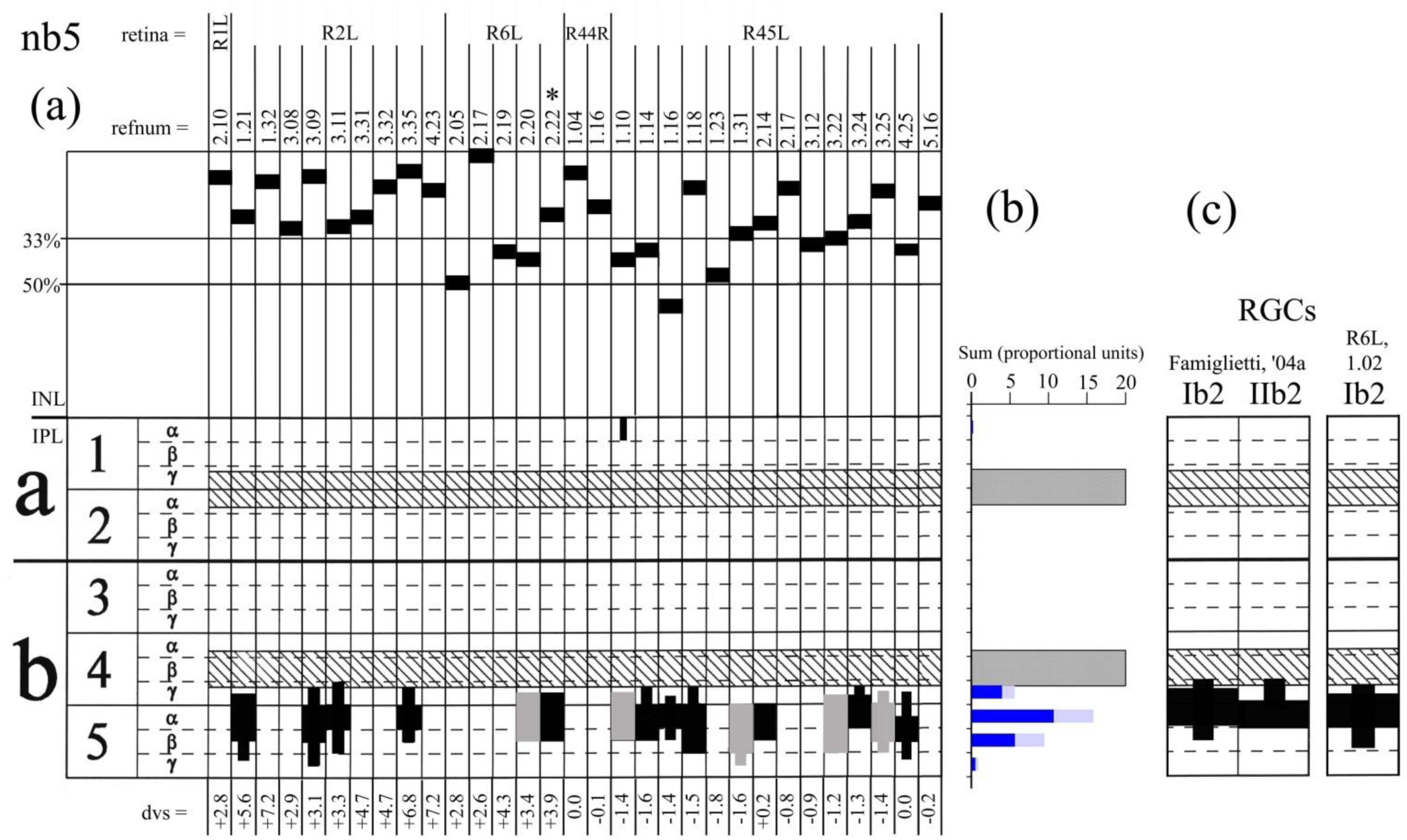
(a) Stratification diagram, illustrating features of Golgi-impregnated nb5 cone bipolar cell somatic position (above) in the INL, and axon terminal stratification (below). Mid-somatic measurements of nb5 cells place most of them in the outer third of the INL consistent with the outer location of calbindin-IR bipolar cell somata (see Figure 29). Note: of 16 cells, the axon terminal of just one reached the dendrites of SAb cells. (b) Sums of contributions to each substratum of the IPL show a concentration in S5α. (c) This peak aligns with the dendritic stratification of two types of large bodied ganglion cell, Ib2 and IIb2 (Famiglietti, 2004a). Fortuitously, one Golgi impregnated nb5 cell, R6L, 2.22 [asterisk in (a)] overlaps both ganglion cell types (Figure 27)

An example of field overlap among three cells is examined in Figure 27: an nb5 cone bipolar cell (R6L, 2.22 in Figure A11), and class Ib2 and IIb2 ganglion cells. In this case, all three are in close contact, essentially co-stratified, at a three-way crossing (arrow in Figure 27). Axon terminal processes of the nb2 bipolar cell both override and pass under the Ib2 dendrite, while the IIb2 cell terminal dendrite overrides both. In three locations (arrowheads in Figure 27), lobules of the nb5 axon terminal lie in the same plane as, and in extensive close contact with the Ib2 dendrite. These appositions are strongly suggestive of synaptic connections.

**FIGURE 27.**
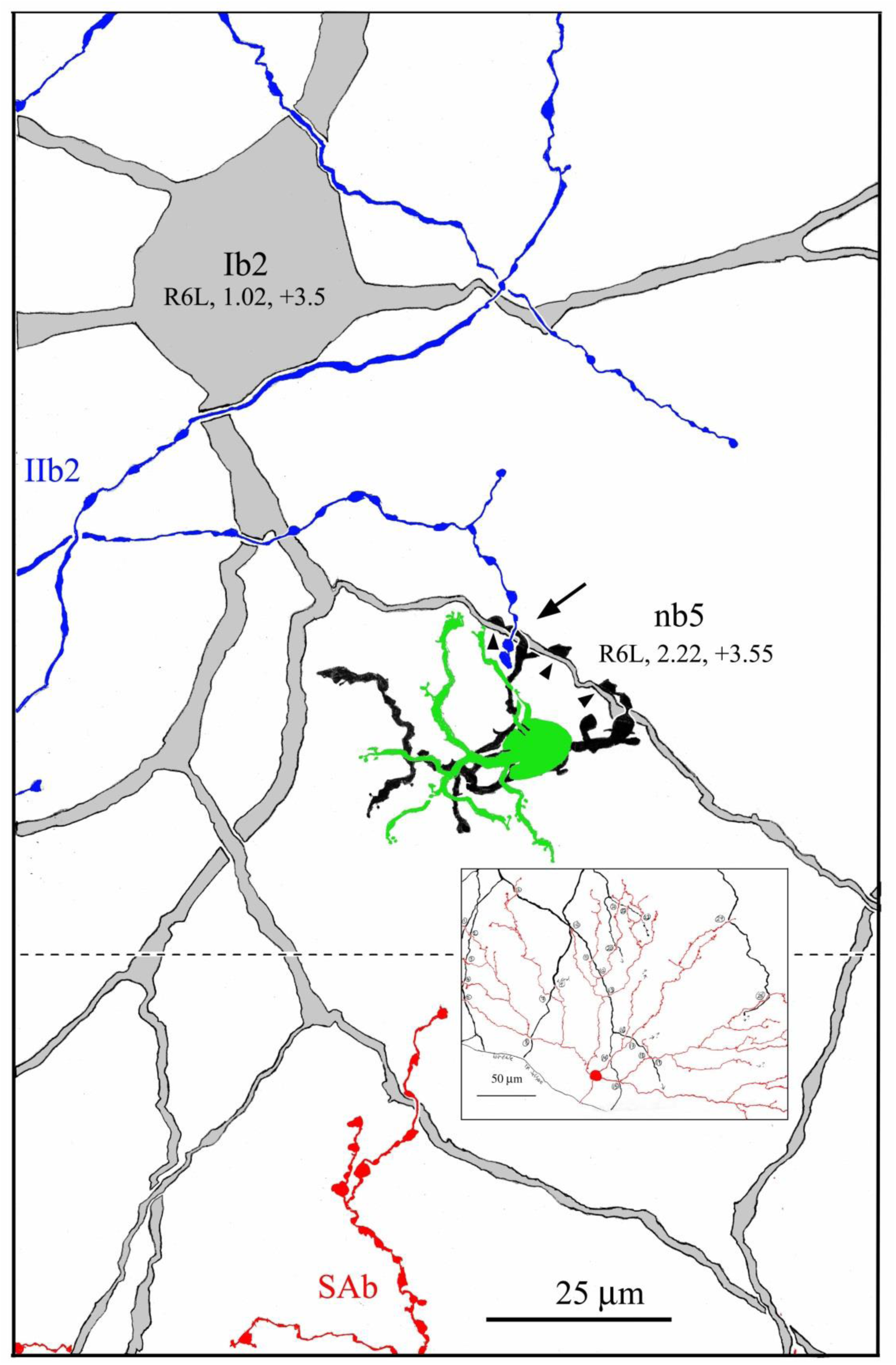
Camera lucida drawings of a class Ib2 (ON-Y) ganglion cell (gray), the distal-most dendrites of a class IIb2 ganglion cell (blue), the dendrites and cell body (green) of an nb5 cone bipolar cell, and its axon terminal (black), in contact with both ganglion cells (arrow), depicted as is viewed from the photoreceptor side, for clarity of presentation.. The type b starburst amacrine cell’s dendrites (red) cross those of the Ib2 cell at 25 locations (inset), providing a fiducial marker for ganglion and bipolar cells. In this case, lobular appendages of the nb5 bipolar cell axon terminal are in close apposition to a dendrite of the Ib2 ganglion cell (arrowheads) in a manner suggestive of synaptic connections. (IIb2 ganglion cells are contacted, instead, by wb cone bipolar cells, conveying blue cone signals; see text)

Additionally, the Ib2 ganglion cell of Figure 27 has significant overlap with a type b starburst amacrine cell (inset, Figure 27). In order to further confirm stratification of the Ib2 ganglion cell and by extension the nb5 cell, differences in stratification were measured at 25 SAb/Ib2 dendritic crossings. The mean separation of the SAb and Ib2 dendrites is 2.4 ± 0.8 µm, virtually identical to the separation of nb5 axon terminals and SAb cells (see above). In this example, Ib2 ganglion cells and nb5 bipolar cells appear to be out of range of starburst amacrine cell synaptic contacts.

In a prior immunocytochemical study, focused upon the localization of GABA_A_ receptor subunits (Sharpe et al., 1993), we noted the similarity in morphology and stratification of Golgi-impregnated “nb3” bipolar cells, as nb5 cell were then known, to bipolar cells labeled by antibodies to the α_1_ subunit of the GABA_A_ receptor, and to calbindin D 28K, as well. We also observed that recoverin appeared to mark the same bipolar cells stained with antibodies to calbindin (Famiglietti and Sharpe, unpublished). Bipolar cell labeling with anti-calbindin was later demonstrated by Massey and Mills (1996), who also showed, in double-labeling studies, that antibodies to calbindin and recoverin label the same bipolar cells. In the context of a comprehensive classification of rabbit bipolar cells, with particular emphasis upon accurate determination of axon terminal stratification, a new analysis of the immunocytochemical evidence is undertaken here, with the hope of more accurate correlation among Golgi-impregnation, imunocytochemical, and published EM studies of rabbit bipolar cells.

Anti-calbindin-, anti-recoverin, and anti-ChAT-immunocytochemistry (ICC) was performed on frozen sections of retina from New Zealand red (pigmented) or white rabbits. In the case of calbindin-ICC, horizontal cells are heavily labeled, a population of bipolar cells is moderately immunoreactive (IR), and a population of amacrine cells is lightly stained (Figure 28a and b). The finer amacrine cell processes form a narrow band of label in the middle of the IPL (arrowhead in Figure 28a), while the more lobular axon terminals of bipolar cells form a band labeling in the innermost portion of the IPL (arrows in Figures 28a and b). Anti-recoverin ICC labels photoreceptors in the outer nuclear layer (ONL in Figures 28c and d), and two populations of bipolar cells, one branching in sublamina a, and the other branching in sublamina b (Figure 28c).

**FIGURE 28.**
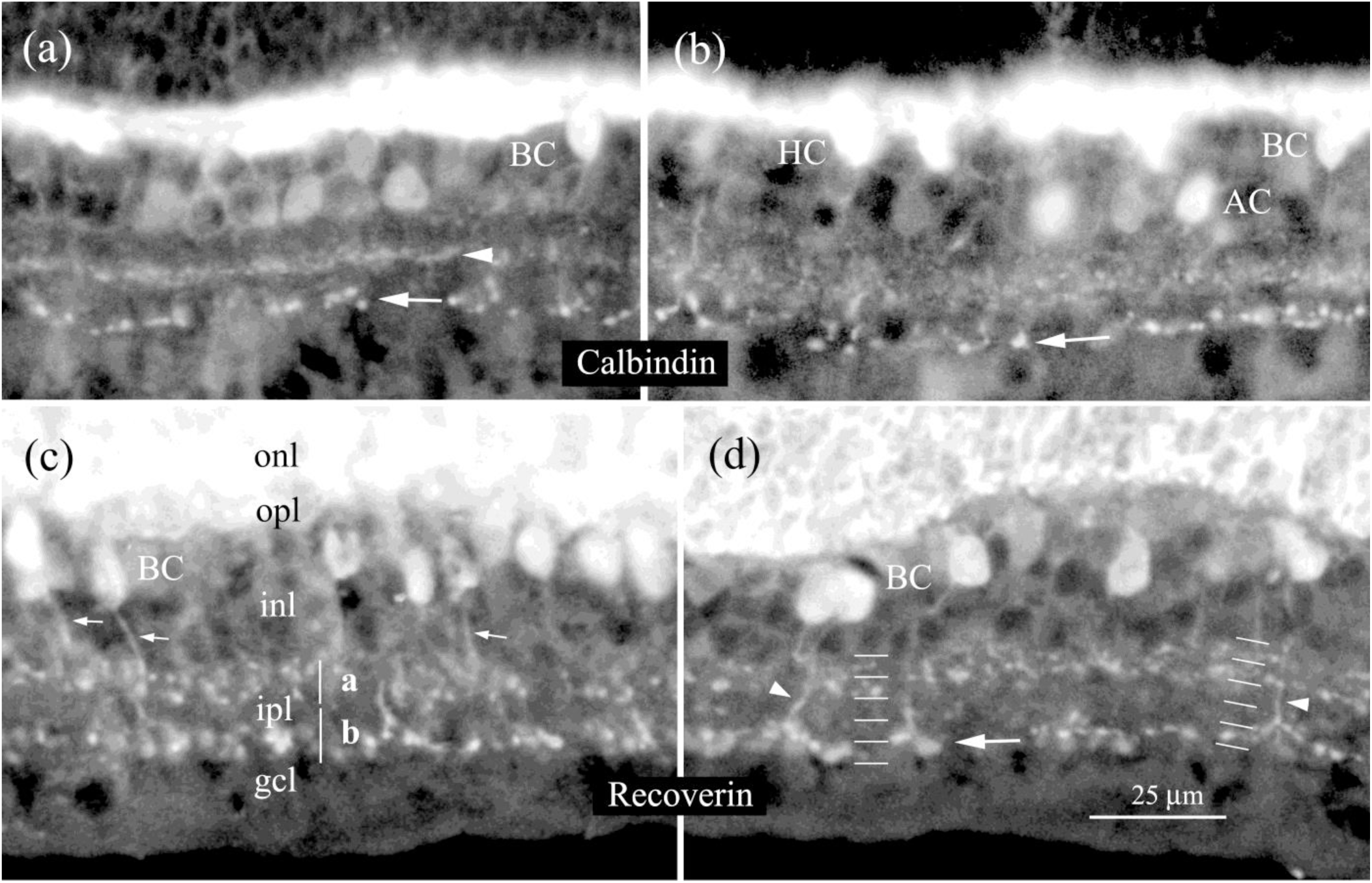
Vertical sections of rabbit retina processed by immunocytochemistry (ICC), using primary antibodies to calbindin D-28K (a), (b) and to Recoverin (c), (d). Calbindin-ICC stains horizontal cells (HC), a single population of amacrine cells (AC), branching in the middle of the IPL (arrowhead) and bipolar cells (BC), the somata of which lie at the level of horizontal cells at the outer margin of the inner nuclear layer (INL), and the axon terminals of which branched deep in the IPL (arrows). Recoverin-ICC stains photoreceptors in the outer nuclear layer (ONL), and two populations of cone bipolar cell in the INL, one of which branches broadly in sublamina a of the IPL [possibly bistratified: (d) right side], and another more narrowly, deep in sublamina b, with more robust terminals [(d). arrow], comparable to calbindin labeling. Descending immunoreactive axons are indicated by small arrows and arrowheads in (c) and (d)

In a more precisely vertical section, recoverin-positive bipolar cell axon terminal branching in sublamina a is confined to S1 and S2 with a suggestion of bistratified branching (right side, Figure 28d). In sublamina b, labeled branching occurs mainly in S5, with likely involvement of S4 as well (arrow, Figure 28d). In regard to the recoverin labeling, these results are consistent with the evidence provided by Massey and Mills (1996) that calbindin and recoverin label the same bipolar cells in sublamina b. In regard to the recoverin labeling in sublamina a, it appears that more lightly labeled bipolar cell bodies are located toward the middle of the INL (Figure 28c and d), and these may give rise to the axons (arrows in Figure 28c), and axon terminals in sublamina a. Based upon their axon terminal stratification and their cell body positions, averaging about 50% of the depth of the INL, these may correspond to na2 bipolar cells of Golgi preparations.

In order to determine the stratification of calbindin/ type b recoverin-labeled bipolar cells with greater precision, double-labeling was performed using antibodies to recoverin and to ChAT (choline acetyltransferase), in the latter case for a fiducial marker labeling type b starburst amacrine cells (Famiglietti & Tumosa, 1987). Double labeled sections were photographed sequentially on sensitive black and white film, using discriminating filter sets, and the two images pseudo-colored and combined (Figure 29a). Selected areas were magnified, rectified for vertical scanning, and scanned at several locations perpendicular to the IPL strata (vertical arrowheads, Figure 29b). The peak intensity of the recoverin-immunoreactive (IR) process closest to the inner ChAT band is 3μm from the peak intensity of the latter (magenta trace, Figure 29c). The width of the ChAT band is estimated to be 1.75-2.0 μm from the high contrast material processed with the peroxidase reaction product as the chromophore (Famiglietti & Tumosa, 1987; black trace in Figure 29c, scans from Figure 2a, above). Consequently, although most of the recoverin-IR processes lie in S5, a few are in the inner half of S4, and fluorescence micrographs suggest that these might occasionally touch the ChAT-IR processes (Figure 29b).

**FIGURE 29.**
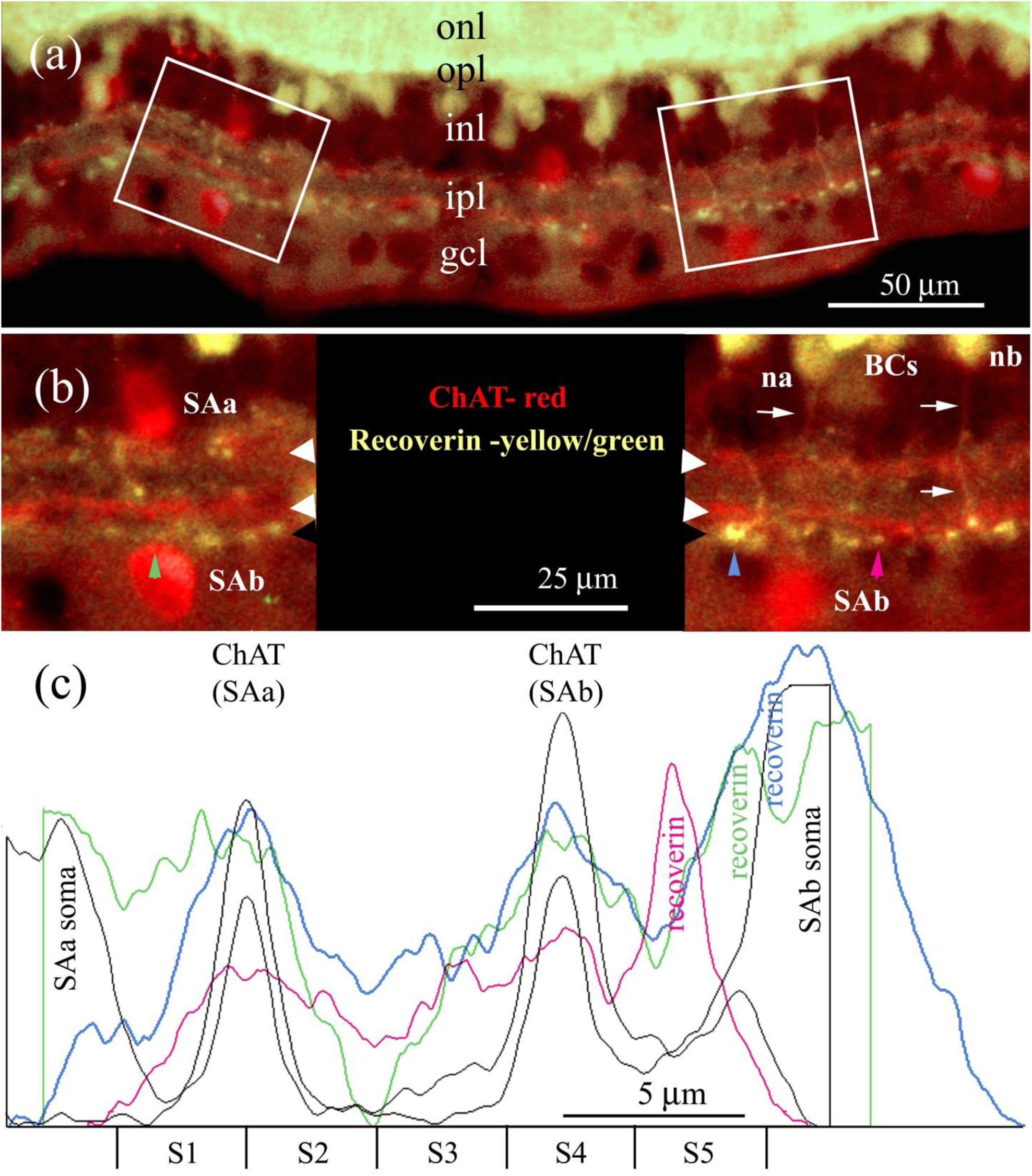
Recoverin-and ChAT-ICC applied to sections of rabbit retina in a double staining protocol. The resulting immunofluorescence is pseudo-colored to maximize visibility: ChAT-ICC in red and recoverin ICC in yellow. (a). Two regions of interest (boxes) are magnified in (b). (b) ChAT-ICC labels the fiducial type a and type b starburst amacrine cells (SAa and SAb) and their dendritic processes, seen in sublaminae a and b (white arrowheads). In sublamina a, the ChAT-immunoreactive (IR) staining and, recoverin-IR staining of the lobular bipolar cell axon terminals are overlapping, whereas below, in sublamina b, they are clearly separate. Vertical densitometric scans are taken in three locations, indicated by vertical arrowheads, color coded for correlation with waveforms displayed in (c). (c) Three scans from (b) and two scans from the higher contrast immunoperoxidase ChAT-ICC of Figure 2A, are aligned at the SAb band. Recoverin-ICC, presumably labeling calbindin-IR cone bipolar cell axon terminals, occurs between 2.5 and 6 µm from the peak of the ChAT-IR band that lies in mid-S4

In view of the many kinds of type b cone bipolar cells, the axon terminals of which branch in stratum 3 and the outer half of S4, and some diversity in dendritic morphology among nb5 bipolar cells, it was necessary to consider the possibility that “nb5”, seemingly the only narrow field type b cone bipolar cell branching in S5, constituted more than a single type of bipolar cell, only one of which is calbindin-IR. Another reason to weigh this possibility was the casual observation that not all nb5 cells appeared to have “high” cell bodies, positioned, as described for calbindin-IR cells by Massey and Mills (1996), “in the highest tier of the inner nuclear layer”. Therefore attempts were made to measure the positions of nb5 bipolar cell somata in the INL. This is challenging to do in whole-mounted Golgi preparations of retina, especially because of the optical screening effect of an entirely opaque spheroidal cell body. Rather than attempt to estimate the upper and lower bounds of the cell body, the depth of the maximum soma area in the retinal plane was determined relative to fiducial markers of the boundaries of the INL. Of 30 examples, 29 had maximum cell body diameters in the outer half of the INL, and 22 of 30 were in the outer third of the INL (Figure 26a). The mean value was 25.4 ± 15% of the thickness of the INL. Close examination of two groups, separated by the 33% boundary (Figure 26a), did not reveal any other features of morphology or axon terminal stratification distinguishing between them (Figure A11).

#### 3.3.6 mb cone bipolar cells

With the possible exception of wide-field cone bipolar cells, and the above-noted similarities of na1 and nb1 bipolar cells, the only cone bipolar cells in rabbit retina that occur as an obvious paramorphic pair (type a and type b morphological counterparts) are the medium wide field bipolar cells ma1 and mb. The former, described in detail above, was discovered in 1981 (Famiglietti, 1981), but the latter is a recent find. At the same retinal location, ma1 and mb cone bipolar cells are almost indistinguishable (Figure 30a), except for axon terminal stratification (Figures 5 and 13), and dimensions. Whereas ma1 cells have larger axon terminal arbors than mb cells throughout the retina (Figures 3e and f), there is no difference in size (Figures 3c and d), or the appearance of ma1 and mb cell dendritic trees (Figures A3 and A12). In the visual streak, where sizes are more uniform among narrow field and medium wide field bipolar cells, mb cells might in some cases be confused with nb1 cells, given the nearness of their axon terminal stratification, and to some degree the similarity of their axon terminal and dendritic morphology (Figure 30b).

**FIGURE 30.**
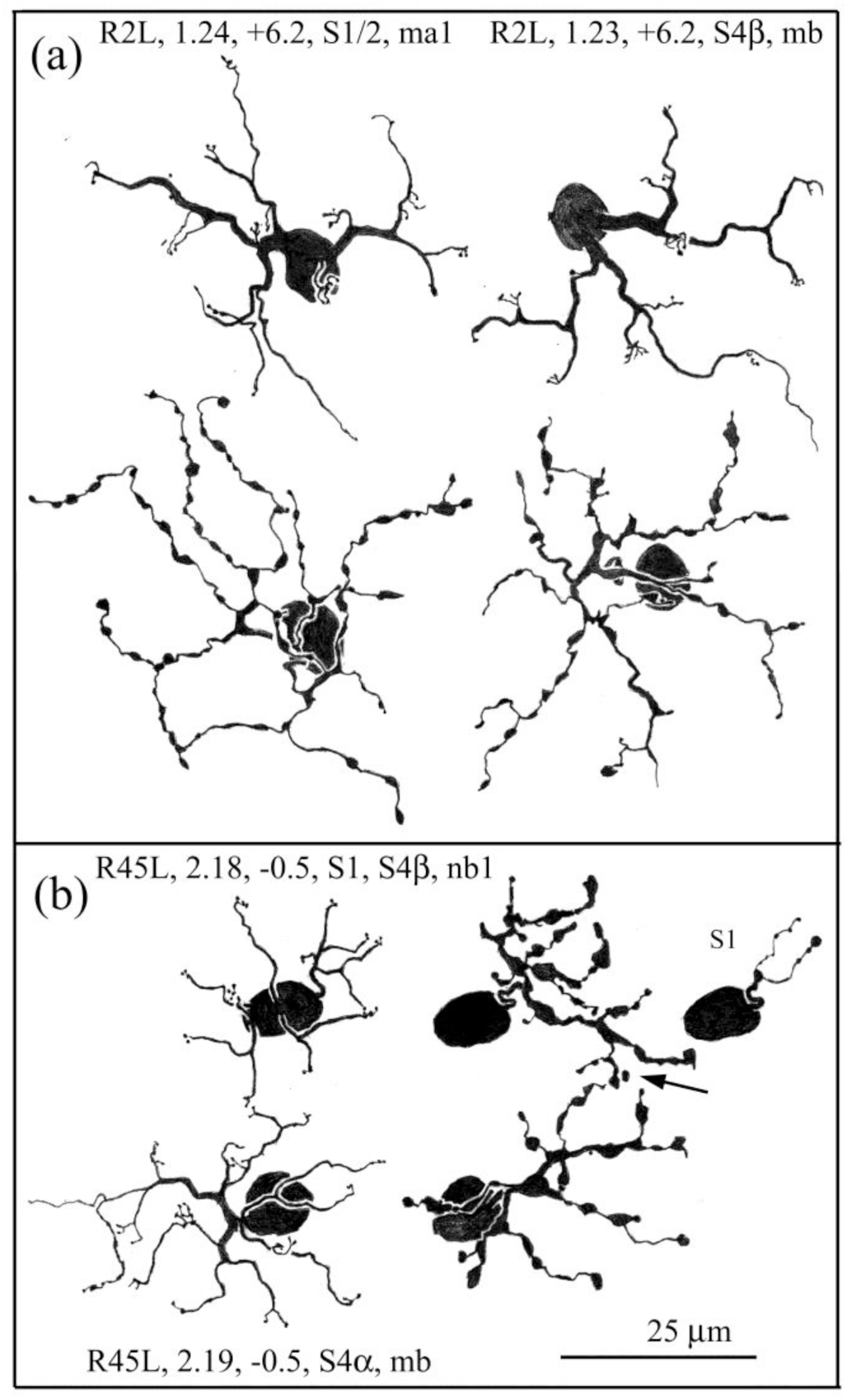
Camera lucida drawings of mb cone bipolar cells, showing that: (a) In flat view, mb cells are morphologically indistinguishable from paramorphic ma1 cone bipolar cells, and that: (b) mb bipolar cells resemble nb1 cells in or near the visual streak, where they are only a little larger (Figure 3d, f).

Of particular interest for understanding the role of mb cone bipolar cells in retinal circuitry, specifically in regard to contacts with retinal ganglion cells, is their axon terminal stratification. This is especially the case, considering their paramorphic pairing with ma1 cells. Paramorphic pairing of “midget” cone bipolar cells and midget ganglion cells is a notable feature of primate retinal organization, but I am unaware of such pairing of very similar type a and type b bipolar cells with paramorphic ganglion cells in reports on other mammalian orders.

The specific postsynaptic ganglion cell partners of ma1 bipolar cells are presently unknown. With regard to their narrow axon terminal stratification, however (Figure 4), two well-known ganglion cell types are possible candidates: BS1 (type 1 bistratified, ON-OFF directionally selective) and class Ia2 (OFF-Y) ganglion cells (Famiglietti, 1992, 2004). The paramorphic relatives of Ia2 ganglion cells are class Ib2 cells, but as the latter branch in S5, they are out of range of mb bipolar cells. Possibly within range, however, is the sublamina b branching of BS1 cells, co-stratified with the fiducial SAb cells (Famiglietti, 1992b). Such a pairing of ma1 and mb bipolar cells with BS1 ganglion cells would afford an agreeable symmetry in parallel connectivity within the IPL

In the available material, Golgi-impregnated mb cone bipolar cells are relatively rare, and they number only 22 in the present sample. Of these, 10, associated with fiducial cells, were candidates for close study of axon terminal stratification. Consequently, it was troubling to observe initially that some mb cells seem to branch predominantly in S3, while others branched mainly in S4. Precise measurements of stratification with respect to SAb cells could be made in three mb cells from retina R2L, and one from retina R6R, at 7 to 14 axo-dendritic crossings per cell (Figure 31). The mean values of the three cells from retina R2L are in a narrow range of 0.5 to 0.8 µm above (sclerad to) the starburst dendrites. In 10 instances, mb and SAb processes were co-stratified, and in one instance an mb axon terminal process was 1µm below the SAb dendrite. In contrast the example from retina R6R branches on average 1.9 µm above SAb dendrites (Figure 31), at a level lying in S3γ, with no crossing processes closer than 1 µm, thus out of the range of starburst amacrine cells. Other examples of mb cells appear to represent a similar range of stratification levels (Figure 32c).

**FIGURE 31.**
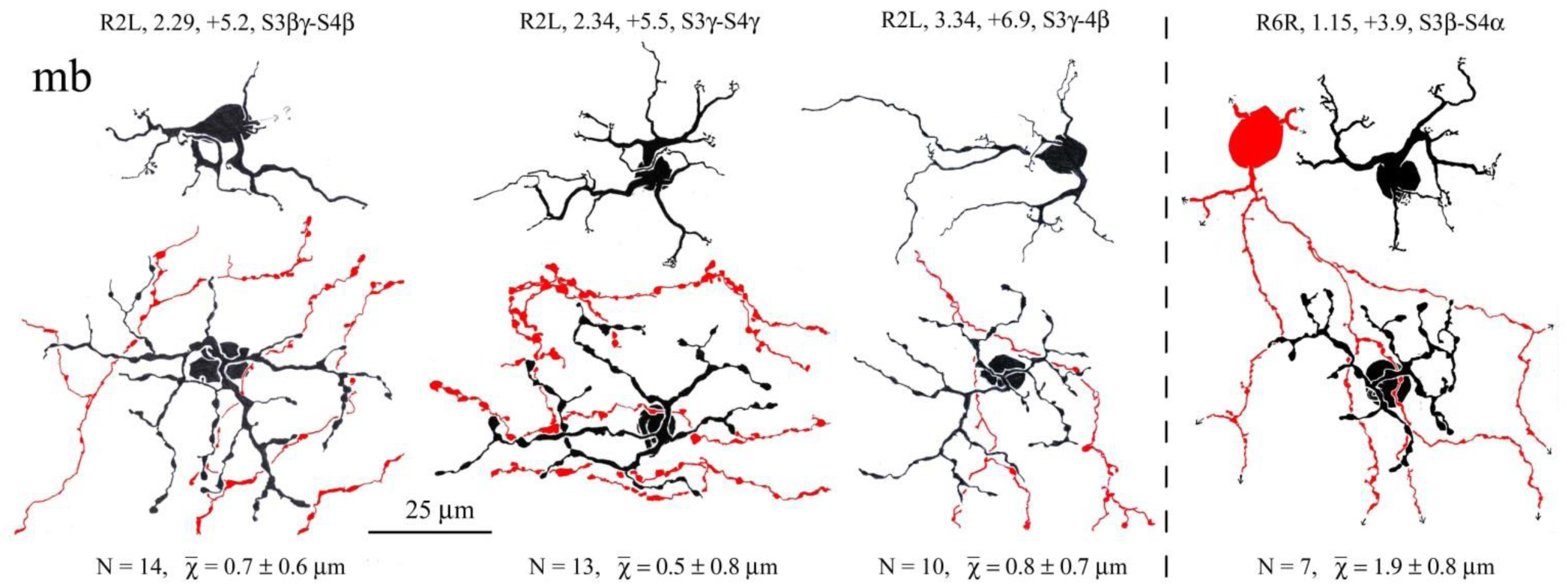
Camera lucida drawings of mb cone bipolar cells from two retinas, together with fiducial SAb cells (red), overlapping their axon terminals. The number of crossings (N) is given in each case, along with the mean distance between them (and the standard deviation), which place the mb cell dendrites above (toward the outer retina) in all cases. Distances are very small in the examples from retina R2L, but greater in R6R

**FIGURE 32.**
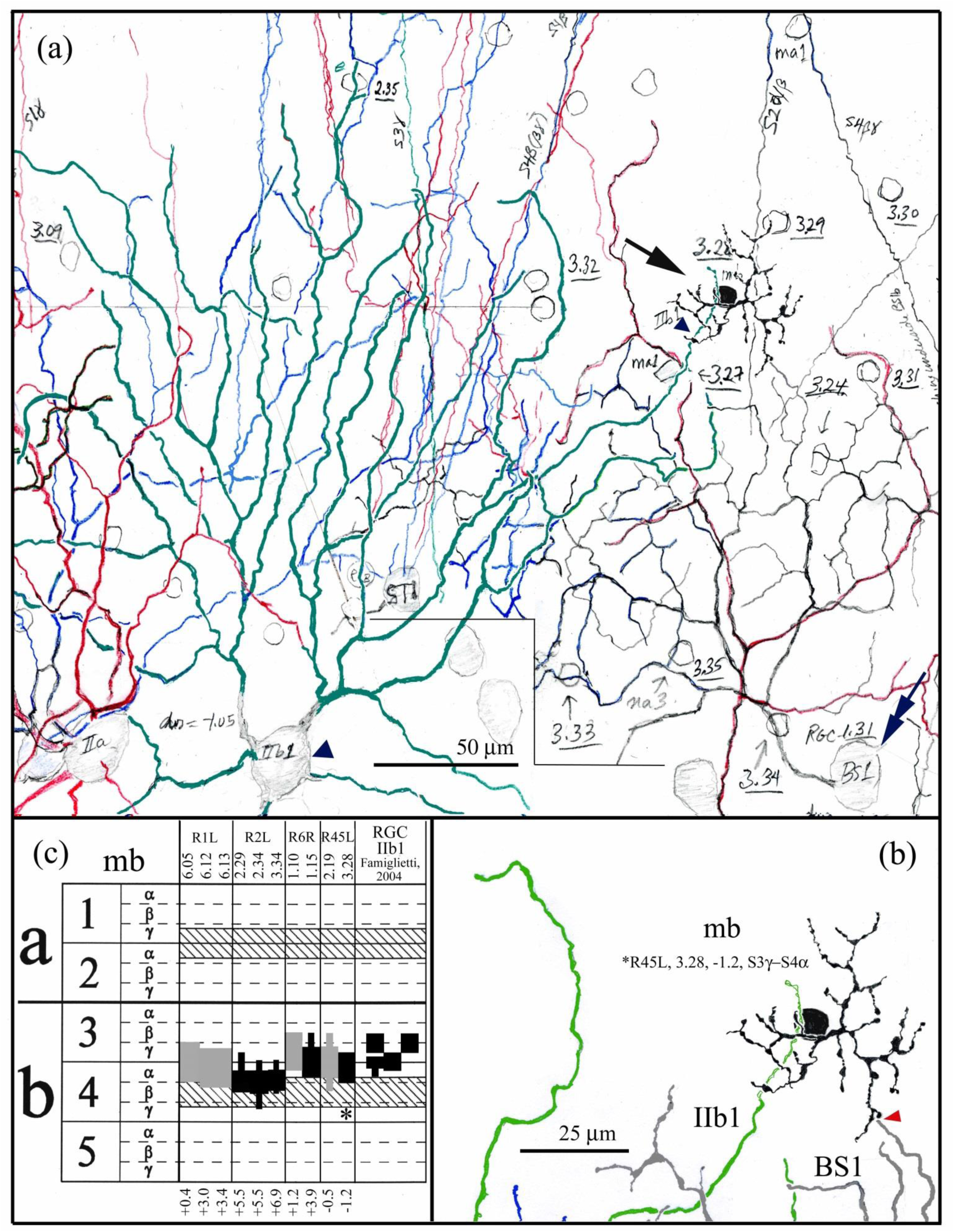
(a) Working drawing, assembled en montage from camera lucida drawings in near dorsal retina, R45L. An mb cone bipolar cell (large arrow) overlaps and contacts a distal dendrite (small arrowhead) of a IIb1 ganglion cell (large arrowhead), previously identified as an “orphan”, class II ganglion cell without a paramorphic mate (Famiglietti, 2004), and subsequently shown to be the transient-ON-directionally selective ganglion cell (see Table 1). (b) The mb bipolar cell axon terminal closely approaches a distal dendrite (red arrowhead) of a fiducial BS1 (ON-OFF DS) ganglion cell [double arrow in (a)]. (c) Stratification of 10 mb bipolar cell axon terminals, only five of which had primary fiducial cells available to confirm stratification (see text for discussion)

**Table 1.** Type a Cone Bipolar Cells of Rabbit Retina: Correlations with Other Studies.

| Cone Bipolar Cell Type | Famiglietti, 1981 | Mills & Massey, 1992 | Merighi et al., 1996 | Mills, 1999 | McGillem & Dacheux, 2001 | Li et al., 2004 | MacNeil et al., 2004 | Liu & Chiao, 2007 | Famiglietti, 2008 | Mills et al., 2014 | Bordt et al., 2019 |
| --- | --- | --- | --- | --- | --- | --- | --- | --- | --- | --- | --- |
| **na1** | na1 | Ba1? |  |  |  | Ba1? |  |  |  |  | na1 |
| **na2** | na2 | Ba1? | Ba1? | Ba1? |  | Ba1? | a2n,<br>a1-2n? |  |  |  | na2 |
| **na3** | na3 | Ba2 | Ba2? |  |  | Ba2? | a1-2? |  |  |  | na3 |
| **na4** | nab |  |  |  |  |  | a2? |  |  |  |  |
| **ma1** | ma | Ba3 | Ba3 | Ba3 | wa1? | Ba3 | a1w | WFa |  |  | ma1 |
| **ma2** |  |  |  |  |  |  |  |  |  |  | ma2 |
| **wa** | wa |  |  |  | wa1? |  |  | BB | wa | S-OFF | wa |

The variability and the location of axon terminal stratification in mb bipolar cells is reminiscent of variability at similar levels of the IPL exhibited by class IIb1 cells (Figure 32c; see Figure 12 in Famiglietti, 2004b). The latter were determined to be “orphan” ganglion cells, in the sense that they do not have a paramorphic, type a counterpart, as do class IIb2 (ON-X) ganglion cells in their pairing with class IIa (OFF-X) ganglion cells (Famiglietti, 2004a). Class IIb1 cells may branch narrowly at the S3β/γ border or at the S3/S4 border, or at both levels. Near the visual streak, where the IPL is thicker, they have broader stratification with some branches in S4γ (Figure 32c). Unexpectedly, class IIb1 cells do not have visual receptive field properties like the morphologically similar IIa and IIb2 ganglion cells. Instead they correspond to the transient ON directionally selective ganglion cells of rabbit retina (Ackert et al., 2009; Ackert et al., 2006; Hoshi et al., 2011; Kanjhan & Sivyer, 2010).

Fortunately, the present sample of mb bipolar cells includes one that overlaps a class IIb1 ganglion cell. It is also immediately adjacent to a BS1 ganglion cells that in turn overlaps the IIb1 cell (Figure 32a and b). At four points of axo-dendritic crossing, the mb and IIb1 cells are co-stratified at one point, touching at a second with the mb cell deeper, at a third separated by 0.5µm and possibly touching, and at a fourth 0.5 to 1.0 µm distant, again with the mb cell deeper. At one point, the tip of BS1 cell dendrite comes very close to an axon terminal lobule of the mb cell (arrowhead in Figure 32b). Here, BS1 dendrite is 0.5µm deeper. (It should be kept in mind that for a neural process 1 µm in diameter, at the mid--focal plane, the process projects 0.5 µm above and below the plane of focus.) As a consequence of these measurements, the narrowly stratified mb cone bipolar cell axon terminal is determined to extend from the middle of S3γ through the depth of S4α, in a planar domain less than 2 µm thick. In addition, measurements made using transitive wide-field, narrowly stratified amacrine cells and other fiducial cells support the observations of variability in stratification of mb cone bipolar cells (Figure 32c).

The dendrites of BS1 and IIb1 ganglion cells in sublamina b, depicted in Figure 32a and b, cross at a number of points. At nine of these the mean difference is 1.9 ± 0.7 µm, a value consistent with previous measurements of the distance between the dendrites of IIb1 cells and SAb cells (Famiglietti, 2016). In summary, mb cone bipolar cells appear to be associated with IIb1 ganglion cells, rather than BS1 ganglion cells, although the tracking of mb cells with IIb1 ganglion cells is imperfect.

#### 3.3.7 wb cone bipolar cells

At the beginning of the previous section, it was mentioned that wide-field (wa and wb) cone bipolar cells may represent a paramorphic pair. Well characterized in previous work (Famiglietti, 1981, 2008), they were recognized as likely to be type a and type b bipolar cells connected exclusively to blue/short wavelength cones, a proposition for which there is substantial support (Liu & Chiao, 2007; MacNeil & Gaul, 2008). Nevertheless, the two types of bipolar cell have significant morphological differences.

In regard to dendritic trees, wa cells have one to four terminal branches that typically end either separately or convergently in clusters of digitiform appendages, signifying cone contacts at the perimeter of the dendritic tree (Figure A4). Similar cone contacts of wb cells, marked by clusters of appendages, usually occur more proximally in the dendritic tree (Figure A13). One or more slender processes extend well beyond these contacts to terminate freely, without appendages. These slender processes resemble slender distal dendritic branches of ma1 and mb dendritic trees (cf. Figures A3 and A12), and in the case of ma1 cells, some of these terminate at rod photoreceptor spherules (Li et al., 2004). It appears that at least some of the distal processes of wb cells do not terminate at cone pedicles (MacNeil & Gaul, 2008). Thus, if such wb dendritic processes terminate in rod spherules, it would be another instance of rod and cone convergence, possibly of value in escape from predation, under crepuscular light, as wb bipolar cells innervate class IIb2/ON-X ganglion cells (Famiglietti, 2008; Mills et al., 2014).

The wa and wb bipolar cells differ in their axon terminals, as well. Although of similar field size (Figure 3e and f), wb axon terminals are robust and often coarsely lobulated (Figure A13), whereas those of wa cells are more delicate tend to have a beaded appearance (Figure A4). The two also differ in stratification. The axon terminals of wa cells, though narrowly stratified in the main, usually have vertical extensions that may cross the a/b sublaminar border (Famiglietti, 2008; and Figure 4, above). The axons of wb cells terminate in narrowly stratified arbors branching in S5α and S5β (Figure 13). In addition, two examples of wb cells exhibit branches arising from the descending axons in S1α (Figure 33). They share this feature with many nb1 cells and one nb5 cell in the present sample. The significance of this feature is taken up in the Discussion.

**FIGURE 33.**
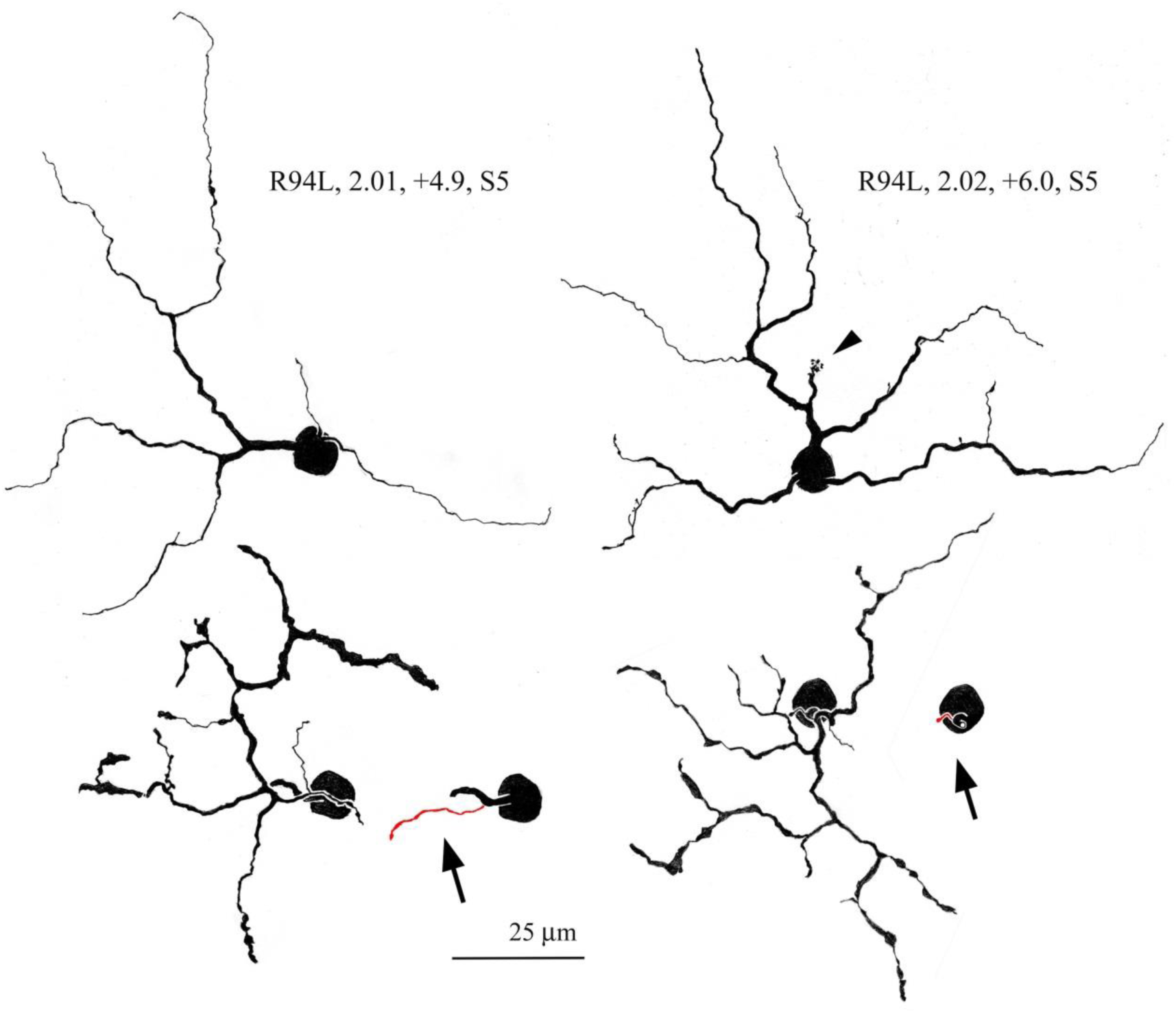
Camera lucida drawings of two wb cone bipolar cells, illustrating their sparsely branching, widespread dendritic trees with very few conspicuous aggregations of terminal digitiform appendages (arrowhead). In these two examples, short branches emanate from axons in S1 (in red, arrows), their axons giving rise to a mixture of robust and slender axon terminal branches deep in sublamina b of the IPL.

#### 3.3.8 rod bipolar cells

Rod bipolar cells are quite uniform in appearance, and in axon terminal stratification that occurs with expansions in S4 and with large lobular appendages in S5 (Figure 34). No evidence has been found for more than a single type of rod bipolar cell in the present study.

**FIGURE 34.**
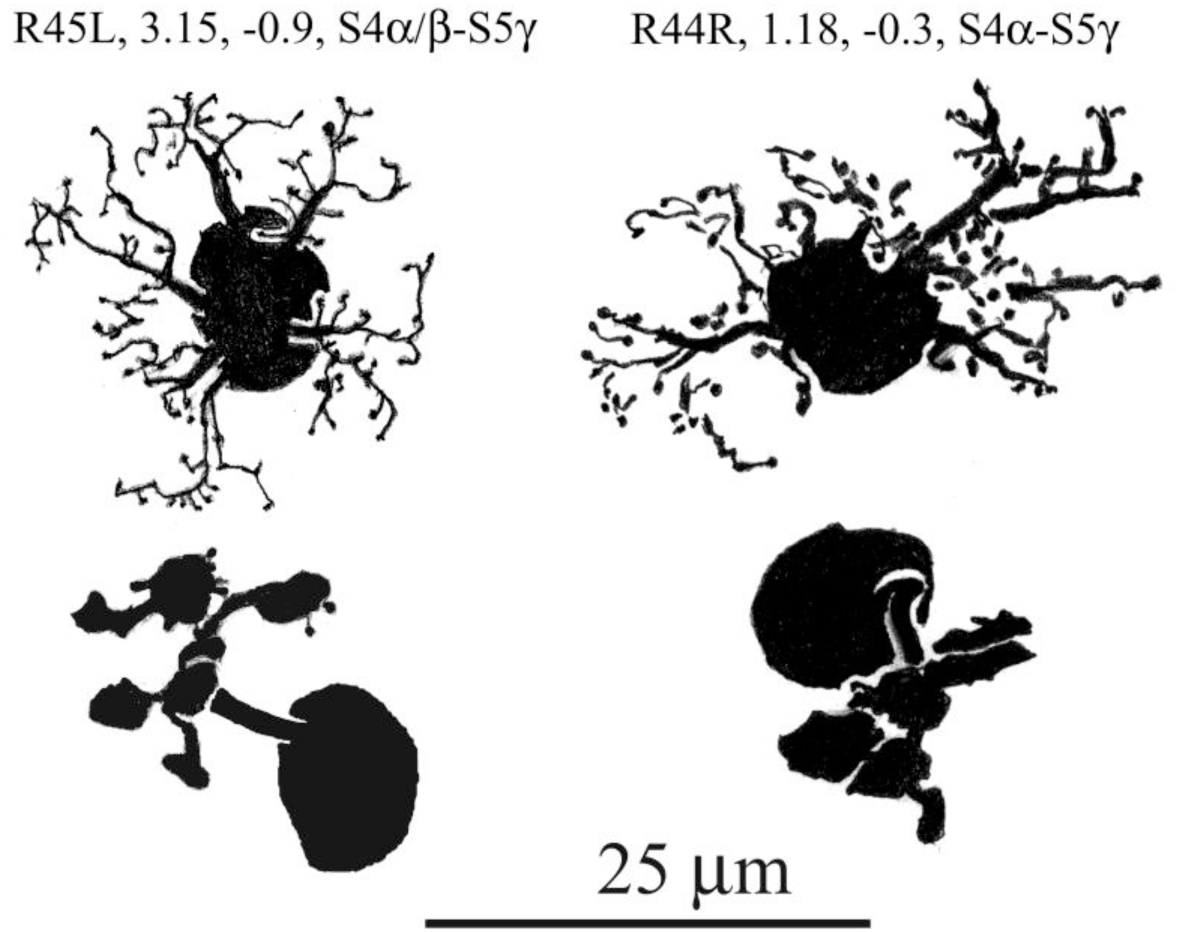
Camera lucida drawings of two rod bipolar cells from different retinas. In typical fashion, short dendritic branches are festooned with slender, sometimes long sinuous terminals. Axon terminals, lying in S4 and S5, form a cluster of bulbous lobular appendages bearing a few short, pedunculated spines.

#### 3.3.9 unclassified cone bipolar cells

A dozen examples, the majority in a single retina, were separated from the rest of the sample, based upon characteristics of the dendritic trees (Figure A14). Their dendritic branching is delicate, bearing some resemblance in this regard to na4 bipolar cells. Unlike the latter, however, the dendrites bear a scattering of slender appendages, sometimes in festoons along the course of preterminal dendrites, and in a modest number of slender terminal spines of varying length, in irregular and sometimes splayed configurations. Their axon terminals, like the dendrites, are in moderately extended conformation in the plane of the retina, and typically have a beaded appearance. Where they are most abundant, in retina R45L, their axon terminal stratification is inconsistent, occurring at several different levels, and may be broad or narrow. In other retinas, examples branching in S1 have also been found (Figure A14). The small number of examples and their inconsistent axon terminal stratification are factors preventing their recognition as a distinct morphological type of cone bipolar cell in rabbit retina.

Other examples of cone bipolar cells, judged to be completely impregnated, but nevertheless morphologically ambiguous, form a second group of unclassified bipolar cells in this study (Figure A15). These constitute 5 to 6% of the sample, despite the nonrandom, selective sampling methods employed (see Methods).

### 4.0 DISCUSSION

The present sample of nearly 800 bipolar cells was taken from ten Golgi-impregnated rabbit retinas, and these were analyzed morphometrically and morphologically, with limited use of immunocytochemical markers. They were determined to consist of 16 kinds of cone bipolar cell, seven of type a and nine of type b, as well as a single type of rod bipolar cell (Figure 35). In addition, the possibility was raised of a seventeenth kind of cone bipolar cell, bearing delicate dendrites festooned with fine terminal appendages (Figure A14). Extensive analysis has been required to distinguish between nb3.1 and nb3.2 cells, differing mainly in level of axon terminal stratification, and the difference between nb2.1 and nb2.2 cells depend upon sometimes subtle differences in dendritic branching and in the terminal concentration of clusters of digitiform appendages. The conclusion is reached, therefore that, at a minimum, there exist 14 kinds of cone bipolar cell in rabbit retina. Should there be a type of bipolar cell in rabbit retina that loses its dendrites and assumes a monopolar form in maturation, as identified in mouse (Della Santina et al., 2016), a type not identifiable in Golgi preparations for technical reasons, then rabbit would have a minimum of 15 types of cone bipolar cell.

**Figure 35.**
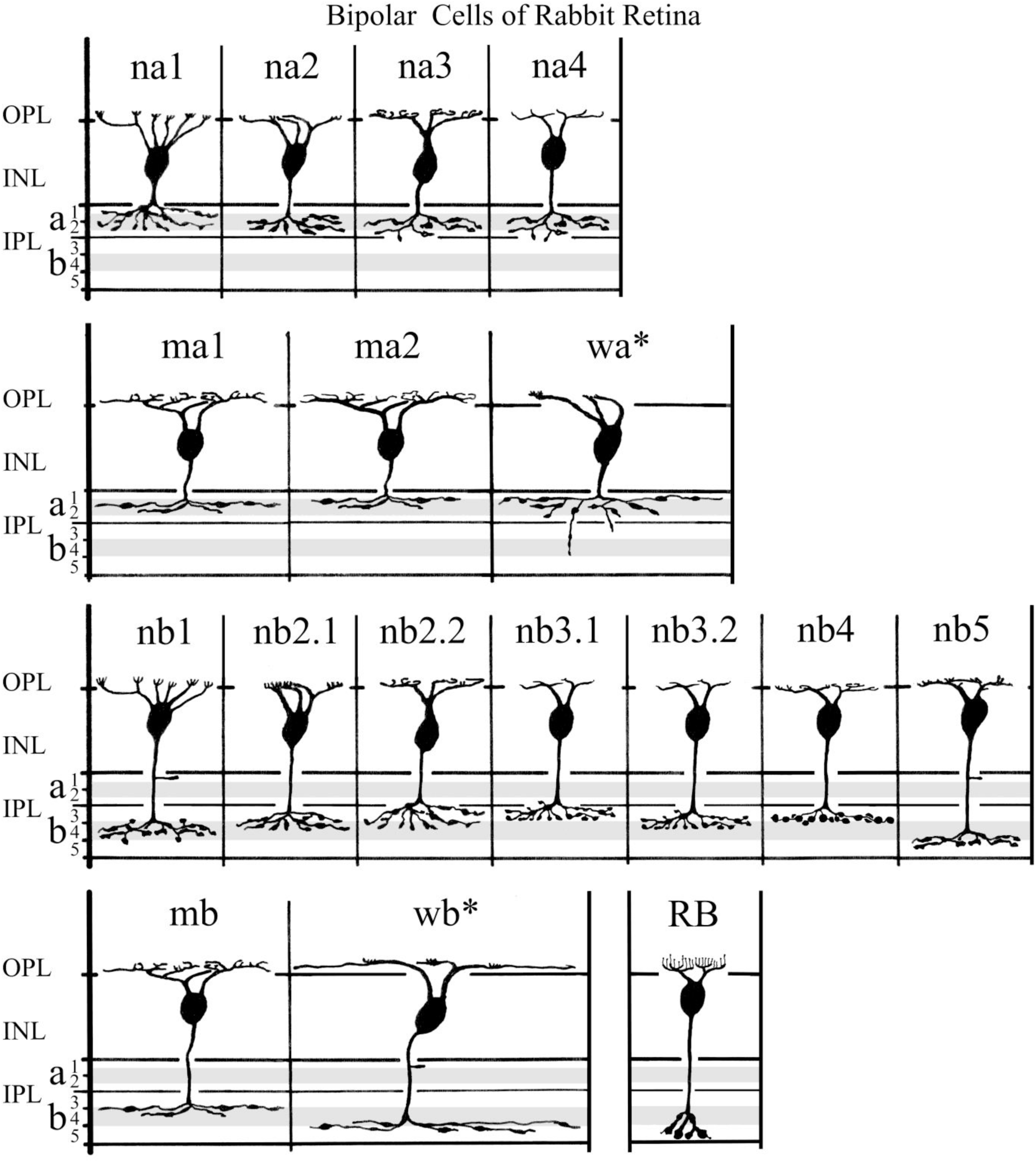
Classification of the bipolar cells in rabbit retina. Sixteen types of cone bipolar cell, and one type of rod bipolar cell were identified. Asterisks indicate bipolar cells connected to blue cones. Gray bands represent the stratification of type a and type b starburst amacrine cells

### 4.1 type a cone bipolar cells

In previous work, five kinds of type a cone bipolar cells were identified: narrow-field “na1”, “na2”, na3”, medium-wide field “ma”, and wide-field “wa” cells (Famiglietti, 1981). These were characterized by details of their dendritic branching, dimensions of their axon terminal and dendritic trees in the plane of the retina, and stratification of their axon terminals in sublamina a. Recently, a second type of medium-wide field, type a cone bipolar cell, ma2, was identified (Bordt et al., 2019), resulting in the renaming of the ma cone bipolar cell”ma1”. The ma1 cells were previously observed to be approximately co-stratified with type a starburst amacrine cells (Famiglietti, 1991) and with BS1 (ON-OFF directionally selective) ganglion cells at the S1/S2 border in sublamina a of the IPL, but proof of synaptic connection has not yet been established.

Three kinds of type a cone bipolar cell, named “Ba1”, “Ba2”, and “Ba3”, were reported to take up diamidino-phenylindole (DAPI) when rabbit retinas are incubated in its presence (Mills & Massey, 1992). There is little doubt that Ba3 cells correspond to the medium-wide field ma1 cells, because of their size and branching; Ba2 likely corresponds to na3, and Ba1 may correspond to na2 (Table 1).. Interestingly, with the combination of ICC and intracellular staining, Ba3/ma1 cells were shown to contact rods; this is also the case for Ba2 cells and to a lesser extent for Ba1 cells (Li et al., 2004), but no type b cone bipolar cells were found to contact rods (Whitaker et al., 2021). In the course of identifying type a bipolar cells connected to blue cones (their BB cells), Liu and colleagues (2007) have also apparently stained ma1 cells (their WFa cells). Immunocytochemistry with antibodies to the hyperpolarization-activated cyclic nucleotide-gated cation channel 1 (HCN1) may also label ma1 cells, for their narrow axon terminal stratification (Figure 36a and b) is unique among the type a cells of the present study, and the same as the latter (Kim et al., 2003). Nearest neighbor analysis of HCN1-IR bipolar cell bodies indicates, however, that these are narrow-field cells, rather than medium-wide-field cells; thus their Golgi-impregnated counterparts are indeterminate.

**FIGURE 36.**
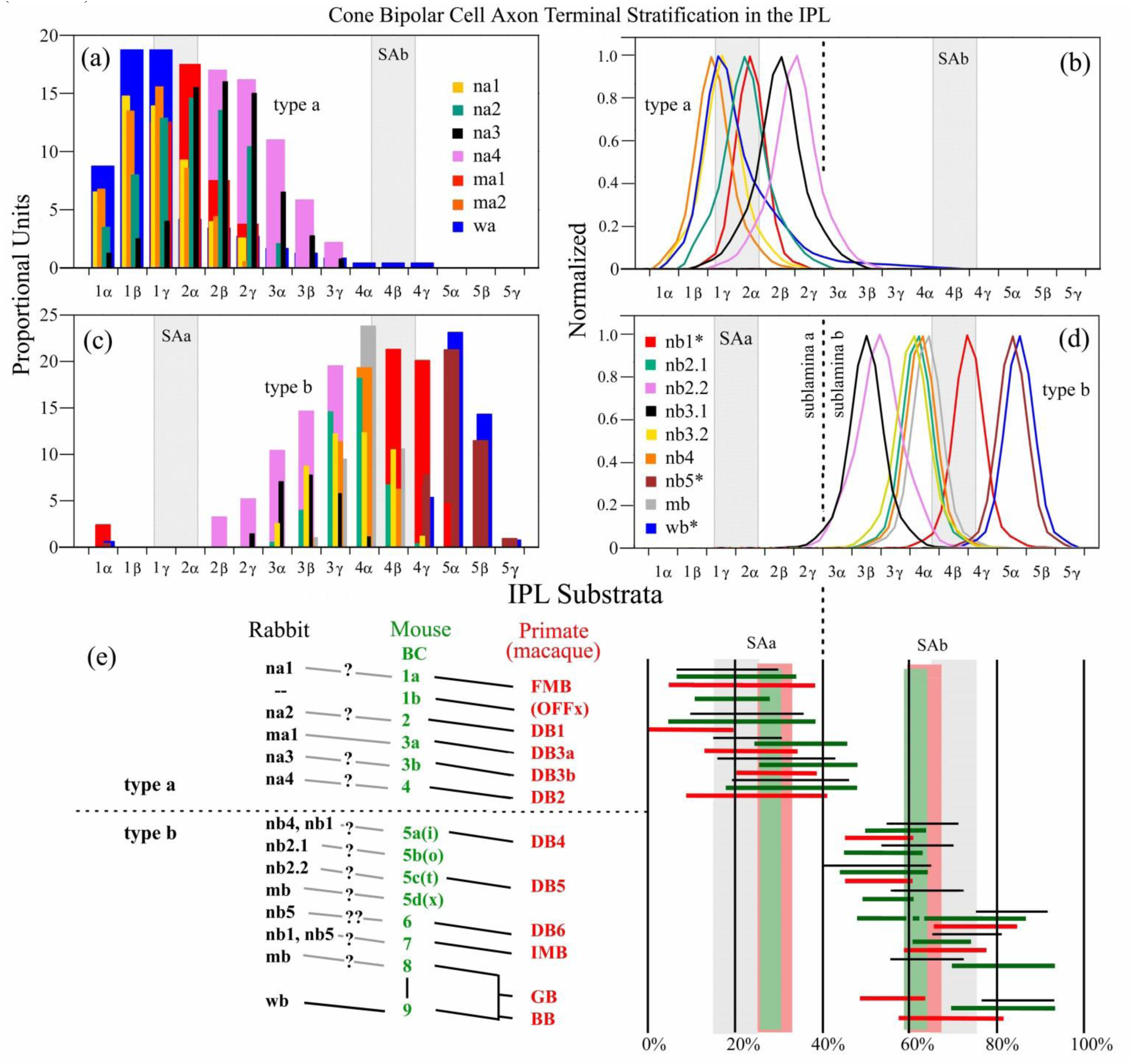
Cone bipolar cell axon terminal stratification in the IPL of rabbit retina, with a comparison to mouse and monkey. (a) and (c) collect the summarized data from multiple cells of the same type, displayed in Figures 5 (type a) and 13 (type b). In order to avoid excess variability, resulting from pooling of data from many retinas and retinal locations, and to give a clearer picture of the overlap in stratification of individual cell types, means are determined for the midrange values of each bipolar cell type and also the average range perpendicular to the strata, of each type. For each of the 16 types, curves were constructed: (b).and (d), using the distance-weighted least squares (DWLS) curve fitting, the peak value representing the midrange mean, and the width at about 12.5% of the height, representing the mean value of the range in the sample of a given type. (b). Despite this compensatory procedure, varieties of type a cell still exhibit considerable overlap. ma1 cells are observed to have the narrowest stratification (red), and na3 (black) and na4 (violet) cells the widest, except that some wa cells (blue) can have one or two processes extending as deep as S4. (d) Type b cone bipolar cells are more spread out. Nevertheless, considerable overlap occurs in S4α, where nb2.1, nb3.2, nb4, and mb bipolar cells have virtually identical stratification. Only nb1 cells (red) occupy the region, centered on the S4β/S4γ border that appears relatively free of other bipolar cell processes. nb1 cells are positioned to provide the maximum input to SAb cells, to the sublamina b branching of BS1/ON-OFF DS, and the dendrites of US1/sustained ON-DS ganglion cells, although bipolar cells with axon terminals in S4α, and nb5 bipolar cells cannot be entirely excluded. (e) Correlations between rabbit (black, gray), mouse (green) and monkey (red, pink) bipolar cells. Mouse data on starburst AC stratification (Bae et al., 2018). Monkey data on starburst AC stratification (Kim et al., 2022). Mouse data on bipolar cell stratification (Greene et al., 2016; Kim et al., 2014). Monkey data on bipolar cell stratification (Tsukamoto & Omi, 2016). Molecular genetic correlations between mouse and monkey are indicated by black lines (Hahn et al., 2023; Peng et al., 2019). Cross-comparisons between rabbit and mouse are indicated by gray or black lines, based upon morphology, axon terminal stratification, and cone connections (wb/BC9). In comparing axon terminal stratification, species differences in the positioning of reference starburst amacrine cell bands should be taken into account.

Formerly, a narrow-field cone bipolar cell, ambiguous in its sublaminar stratification, was named.”nab” (Famiglietti, 1981). In the present study, closer examination shows that while their dendritic branching is very different, they do not differ significantly from na3 cells in their axon terminal stratification. The variable crossing of the a/b sublaminar border, noted in the prior study, is apparently due in part to individual variation among animal subjects. Consequently, nab cone bipolar cells have been renamed “na4” cells.

The only rabbit cone bipolar cells branching principally in sublamina a, for which a specific function is known, are wa cells, proposed to connect selectively to blue/short wavelength cones (Famiglietti, 1981). The study of the spacing of their putative cone connections in Golgi preparations also yielded a prediction that the density of blue cones unexpectedly increases with increasing ventral distance from the visual streak. The first proposal was proved, using a combination of ICC and intracellular staining (Liu & Chiao, 2007) (Table 1), and the second using ICC with antibodies selective for the two principal types of cone in rabbit retina (Famiglietti & Sharpe, 1994, 1995; Juliusson et al., 1994).

**Tables 1 and 2**. Abreviations:: n, narrow-field; m, medium-wide field; w, wide field; wf/WF, wide-field; a, type a, branching in sublamina a of the inner plexiform layer (IPL); b, type b, branching in sublamina b of the inner plexiform layer (IPL); CaBP, immunoreactive for calbindin-28K calcium-binding protein; recov., immunoreactive for recoverin, a calcium-activated sensor of guanylate cyclase; CD15, immunoreactive for CD15, a carbohydrate epitope expressed by a variety of immune and neural cells; NK1, immunoreactive for the neurokinin 1 receptor. Citations, See References.

**Table 2.**
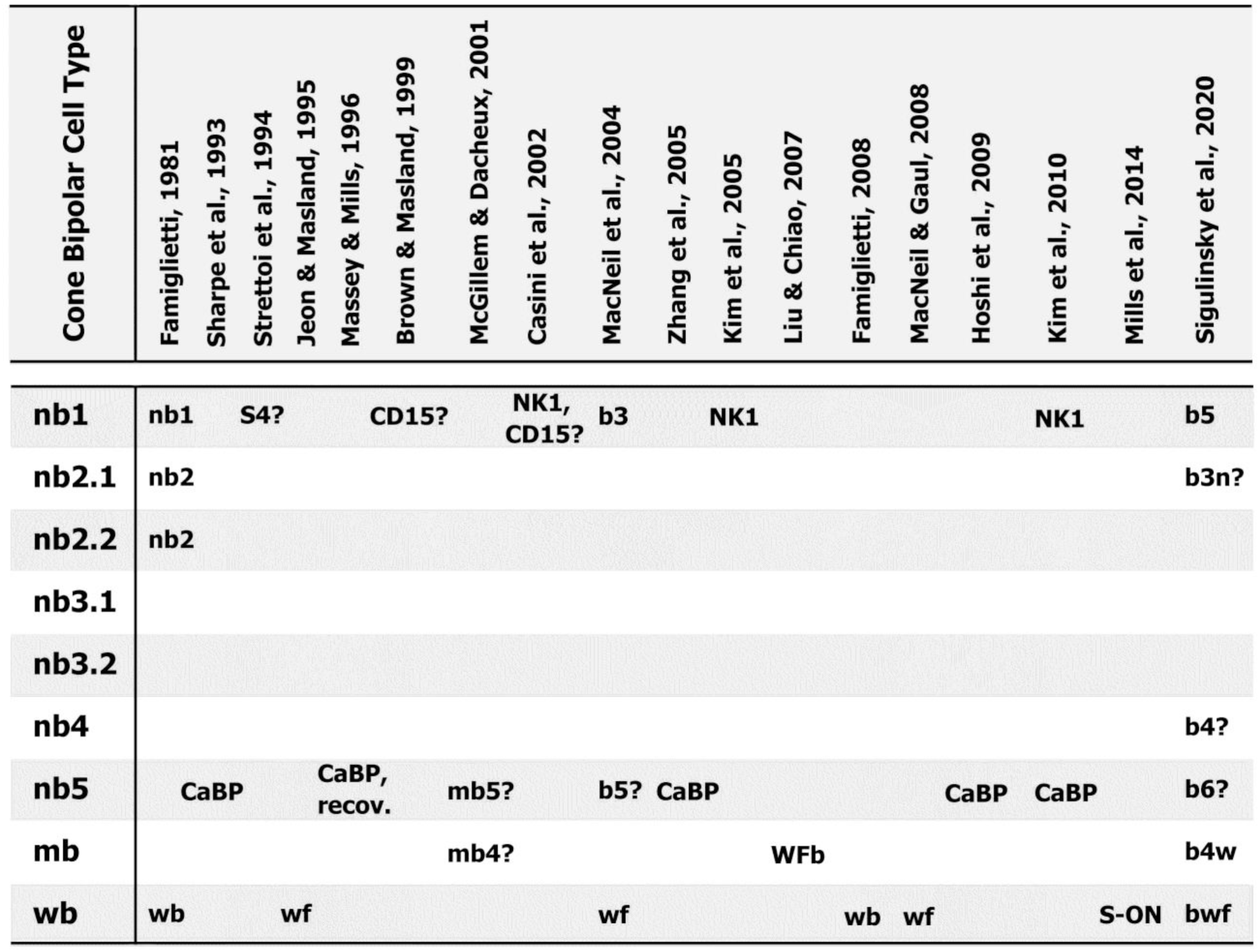
Type b Cone Bipolar Cells of Rabbit Retina: Correlations with Other Studies.

The peculiar ventral distribution of short wavelength sensitive cones that rabbits share with mice (Famiglietti & Sharpe, 1994, 1995; Juliusson et al., 1994; Szel et al., 1992) is a possible explanation for the extreme variability in dendritic field size and appearance of wa cone bipolar cells (Figures 3c, 10 and A4). The lower frequency of blue cones in and near the visual streak, about 1500/ mm^2^ or 9% of cones (Famiglietti & Sharpe, 1995) presents a sampling problem for blue cone bipolar cells, sometimes apparently resulting in contact with only a single cone. If such a bipolar cell directly underlies such a cone, the dendritic field can be very small, indeed, as observed in some examples lying in or near the visual streak (Figure A4).

The axon terminal stratification of wa cone bipolar cells is different from all others, in regard to its depth. Although the majority of wa axonal processes branch horizontally in S1, a small number descend vertically, entering S2. A few may cross the a/b sublaminar border into S3 and even S4, although not to the level of wb (blue-ON) cone bipolar cells. The principal destination of wa axon terminals is the type 3 bistratified ganglion cell (BS3 in Famiglietti, 2009), the only ganglion cell with dendrites in sublamina a known to give OFF responses to blue light (Mills et al., 2014). The BS3 cell’s branching is more developed in sublamina a than in sublamina b and coincides with the main level of axon terminal branching of wa cells (Figure 36b and Famiglietti, 2009). Still unknown, however, is the purpose of the BS3 cell’s sublamina b branching in S5α or the descending branches of the wa bipolar cell’s axon terminal, which do not reach that level.

Previous surveys of rabbit bipolar cells, based on intracellular staining with the fluorescent dye sulfarhodamine B (McGillem & Dacheux, 2001), or Golgi-impregnation (MacNeil et al., 2004) are difficult to reconcile with the results of the present study, but attempts at correlation have been made in Table 1. In the former study of sectioned material, the principal difficulties are: inability to visualize the dendritic trees in sufficient detail, doubtful completeness of cells in sectioned material and hence limited ability to evaluate size, and finally the comparatively indistinct representation fine features in conventional fluorescence microscopy. In the latter study, few examples and limited information on retinal location hamper evaluation. Moreover, dendritic features, notably absent in a summary drawing, are accorded little importance (MacNeil et al., 2004), and lack of adequate fiducial markers for axon terminal stratification make correlation difficult. Nevertheless, a few comparisons can be made. “CBa1_w_” cells are very likely ma1 cells, as MacNeil and colleagues surmise (MacNeil et al., 2004). On the other hand, based largely upon dendritic features illustrated, “CBa2n” cells are likely na2 cells, and CBa2 may be na4 cells (Table 1).

### 4.2 type b cone bipolar cells

In the previous study (Famiglietti, 1981), only three kinds of type b cone bipolar cell were identified in Golgi preparations: two narrow field cells, nb1 and nb2, and a wide field bipolar cell, wb. As noted above, a third type of narrow field cone bipolar cell branching in S5, “nb3”, was later recognized (Famiglietti, 2002; Sharpe et al., 1993), now termed “nb5”. In the present study, “nb2” cells have been subdivided, in recognition of the fact that two distinctive dendritic features, formally attributed to nb2 cells, are differentially expressed in two different types: “nb2.1” and “nb2.2”. In addition to these five, four others have been newly discovered: “nb3.1”, “nb3.2”, “nb4”, and “mb”, based upon dendritic features, axon terminal stratification, and size, for a total of nine type b cone bipolar cell varieties.

#### 4.2.1 wb cone bipolar cells

The wb cone bipolar cells, putative “blue-ON” bipolar cells (Famiglietti, 1981), are now known to connect to blue cones, based upon ICC with antibodies to cone opsins, combined with biocytin uptake into wb cells (MacNeil & Gaul, 2008). In a previous biocytin uptake study, wb cone bipolar cells were shown to form networks of dendritic and axonal processes (Jeon & Masland, 1995), suggesting mutual recognition and potential cellular coupling. Unlike wa cells, the dendrites of wb cells typically have long, thin processes extending beyond clusters of appendages contacting cones, terminating without appendages. In the work of MacNeil and Gaul (2008), it appears that at least some of these terminals do not contact cones. These processes are reminiscent of terminal dendritic branches of Ba3/ma1 bipolar cells shown to contact rods (Li et al., 2004), and perhaps such processes of wb cells also contact rods. Rod contacts of “cone” bipolar cells may in fact be a relatively common feature in mammalian retina (e.g. Hack et al., 1999), as formerly shown in “mixed” bipolar cells of goldfish retina (Ishida et al., 1980). In the present study, two wb cone bipolar cells are also shown to produce small axonal branches in S1, as first shown for nb1 bipolar cells (Famiglietti, 2002).

#### 4.2.2 nb1 and nb4 cone bipolar cells

The nb1 cone bipolar cells were among the first of the narrow field bipolar cells identified in rabbit retina (Famiglietti, 1981), as a consequence of their distinctive, extended dendritic trees with independent terminal dendrites bearing compact clusters of digitiform appendages, and axon terminal stratification, closely associated with type b starburst amacrine cells. Of all rabbit cone bipolar cells, nb1 cells are unique in regard to their dendritic terminal clusters, constituted of slender digitiform processes ascending vertically, and potentially forming the majority of the invaginating synaptic contacts with mid-wavelength cone pedicles. In the course of the present comprehensive study, it gradually became clear, however, with the accumulation of examples, that some narrow field bipolar cells of about the same size as nb1 bipolar cells, with close but different axon terminal stratification, and similar extended dendritic branching pattern, but without terminal clusters of appendages, constituted a new type of bipolar cell, “nb4”, rather than incompletely impregnated nb1 cells.

Close examination of the axon terminal stratification of nb1 cells shows that they are either co-stratified with SAb cells (Famiglietti, 2002), or branch slightly below them, with some lobular appendages extending upward to co-stratify with the SAb cells. In contrast, nb4 cells have narrower axon terminal stratification just above SAb dendrites, sometimes with lobules descending to co-stratify with the latter. Another important distinction between nb1 and nb4 cells is that 25% of the nb1 cells in the present sample gave rise to an axonal branch in S1, shown in one instance to contact the dendrites of a PA4 polyaxonal (presumed dopaminergic) amacrine cell (Famiglietti, 2002), whereas none of the 86 nb4 cells exhibited such processes in S1. Such small branches were also observed in a serial-section, EM, 3D reconstruction study of “CBb5” bipolar cells (Lauritzen et al., 2013), likely correlates of nb1 cells (see below).

Brown and Masland (1999) labeled a population of narrow field cone bipolar cells by immunocytochemical means using antibodies to CD15, demonstrating a degree of stratification with ChAT-IR SAbs, and thus concluding that these bipolar cells play a role in the mechanism of direction selectivity,. They stated that the CD15-IR bipolar cells branch just above the SAb band, with lobules descending into the matrix of ChAT-IR dendrites. In the present study, the axon terminals of some nb1 cells in one retina had prominent descending lobular appendages like those described by Brown and Masland (1999), but the precise stratification of these cells could not be verified with fiducial cells. More commonly, nb1 cells were co-stratified with fiducial SAb cells, or else branching slightly deep to SAbs with both ascending and descending lobular axon terminal processes. Unfortunately, published photomicrographs of the CD15-IR dendrites afforded insufficient resolution of either of dendritic terminal branches or axon terminal branching in S1 to identify these cells definitively as nb1 or nb4 cells.

Two other studies relevant to the distinction between nb1 and nb4 cone bipolar cells select a population of type b bipolar cells by means of neurokinin-1-receptor (NK1R) immunoreactivity (Casini et al., 2002; Kim et al., 2005; Kim et al., 2010). In double-labeling studies, these bipolar cells are not labeled with antibodies to protein kinase C (PKC), CaPB (Casini et al., 2002; Kim et al., 2010), or recoverin (Casini et al., 2002). They are reported as distinct from CD15-IR cells, “with terminal axons arborizing at about the same level of the IPL”, but rarely, double-labeled cell bodies, dendrites, and axon terminal processes are observed (Casini et al., 2002). In triple labeling studies, Kim and colleagues (2010) show quite clearly that NK1R-IR bipolar cell axon terminals lie between the dendrites of SAb cells and the axon terminals of calbindin-IR bipolar cells, immediately adjacent to the SAb band. In an earlier study, they display a well-oriented 50 µm thick section, stained for NK1R-IR, in which it can be estimated that the stained bipolar cell axon terminals lie between 67 and 76% of the IPL’s thickness (Kim et al., 2005). This corresponds closely to the axon terminal stratification determined for nb1 bipolar cells in the present study, and the peak densities of stratification of the two nearest kinds of cone bipolar cell, nb4 and nb5, lie more than 10% distant from that of nb1 cells (Figure 36d).

More detailed examination of NK1R-IR dendrites may also be of value, for an array of immunoreactive “puncta” are seen in horizontal sections of the OPL, connected one to another in a network by relatively straight dendritic branches (Casini et al., 2002). These puncta, about 2 µm in diameter, appear lobulated and are suggestive of a convergence of two or three clusters of digitiform appendages of nb1 bipolar cells near the center of single cone pedicles (cf. Figures 11 and 12, above). The similarities noted here between nb1, NK1R-IR, and CD15-IR cone bipolar cells raises two possibilities: either 1) another kind of cone bipolar cell similar to nb1 cells in axon terminal stratification was missed in the present study, or 2) under some conditions NK1R-IR and CD15-IR label the same bipolar cells. In regard to the former possibility, an exhaustive EM reconstruction study of the RC1 data set (Sigulinsky et al., 2020) does not support the presence of another,”missed” cone bipolar cell with axon terminal branching at about the same level as that of nb1 cells. It can be said with a relative degree of confidence that NK1R-IR labels nb1 cone bipolar cells (Table 2).

Strettoi and colleagues (1994) performed EM-3D-reconstruction of three type b cone bipolar cell axon terminals that they had previously studied in relationship to AII/rod amacrine cells. They reported them to be “narrowly confined to stratum 4 (S4) of the IPL”, and an illustrations of the axon terminals shows narrow stratification and lobular expansions, notably scalloped in contour. Moreover, a drawing of a Golgi-impregnated bipolar cell, believed to be of the same type, is suggestive of an nb1 cell. Quantitation of the output of the “S4” bipolar cells was 30.0% to ganglion cells, 51.7% to amacrine cells, and 18.3% to unidentified processes. Interestingly, these values are quite similar to those found for NK1R-IR bipolar cells: 28.2% to ganglion cells,, 60.1% to amacrine cells and 11.7% to unidentified processes (Kim et al., 2005). They also found that these bipolar cells make a large number of gap junctions with AII amacrine cell. These are ultrastructural features also examined by Sigulinsky and colleagues (2020) in a serial-section EM study.

That study reconstructs neurons in the inner retinal layers of a disc of retinal tissue (RC1) 250 µm in diameter, and the axon terminal connectivity of 178 type b cone bipolar cells within RC1, and in the process classifies 80% of them into seven types, based upon connectivity, as well as axon terminal stratification and field size. Useful comparison of results from that work and the present study can be made (Table 2), despite the absence of data concerning dendrites, and uncertainty about the (peripheral) retinal location of RC1.

In the study of Brown and Masland (1999), CD15-IR bipolar cells were seen to form a continuous network in peripheral retina, and similar continuity of axon terminal processes was found in the present study of Golgi-impregnated nb1 bipolar cells, suggestive of electrotonic coupling. Notably, in RC1, CBb5 bipolar cells were highly interconnected by gap junctions (Sigulinsky et al., 2020). A more specific correlation between nb1 and CBb5 cells is axon terminal stratification. That of CBb5 cone bipolar cells is reported as ranging from 59% to 86% of IPL depth (Sigulinsky et al., 2020), corresponding very closely to the stratification range of nb1 cells: S3/4 (60%) through S5α (86.7%) (Figures 13 and 36c). Moreover, CBb5 cells were found to provide more than half the bipolar cell input to SAb cells (Marc et al., 2018).

Among other narrow-field cells, the stratification of CBb3n (n = narrow field size), CBb3, and CBb4 cells is centered at about the 60% level (Sigulinsky et al., 2020), as is that of nb2.1, nb3.2 and nb4 cells (Figures 13 and 36). There is some resemblance in axon terminal morphology and branching pattern between nb4 and CBb4 cells. Furthermore, comparing axon terminal field sizes of the three types in each study, CBb4 and nb4 are both the largest. On the other hand, CBb4 cells make virtually no synapses with SAb cells, nor do CBb3 cells, and CBb3n cells make only a very few (Marc et al., 2018), whereas nb4 cells are often in contact with SAb cells, with the expectation, based upon proximity, of some synaptic connection.

#### 4.2.3 nb2 cone bipolar cells

The nb2 cone bipolar cells, originally identified based upon dendritic morphology and levels of axon terminal branching (Famiglietti, 1981) are subdivided here in two groups, based principally upon the presence or absence of multitudes of digitiform terminal appendages. They also have different patterns of axon terminal stratification (Figure 11). Based upon the fact that nb2.1 cells have the smallest axon terminal field diameters (Figure 3f), and (Zhang et al., 2005)the correspondence in stratification between nb2.1 and CBb3n bipolar cells, nb2.1 cells may correspond to the CBb3n bipolar cells of RC1 (Sigulinsky et al., 2020) (Table 2).

#### 4.2.4 nb3 cone bipolar cells

The nb3 cone bipolar cells have long been observed with caution in the study described here, due to their compact, and sparsely branched dendritic trees mostly lacking in appendages, out of concern for the possibility that they represent incompletely impregnated bipolar cells. They have been seen in sufficient numbers, however, in all retinas examined (N = 108, drawn and analyzed), with similar axon terminal morphology, to justify regarding them as complete and classifying them in a separate group. The nb3 bipolar cells have been subdivided in two, based upon differences in axon terminal stratification. Given the partial overlap in the distribution of axon terminal stratification levels of nb3.1 and nb3.2 cells, the validity of this bipartite subdivision is open to interpretation, but should be weighed in the context of evidence in other species for “pairing” (see below), in ground squirrel (Light et al., 2012), and mouse (Shekhar et al., 2016).

#### 4.2.5 nb5 cone bipolar cells

In mouse retina, the axon terminals of three types of cone bipolar cell seem to branch in the inner 20% of the IPL (Helmstaedter et al., 2013; Sabbah et al., 2022; Tsukamoto & Omi, 2017; Wassle et al., 2009), although in some studies the broadly stratified type 6/CB3,4 bipolar cell appears not enter S5 (Pignatelli & Strettoi, 2004). Therefore, the possibility had to be entertained that in rabbit retina the axon terminals of more than one type of cone bipolar cell branch in S5 together with wb cone bipolar cells. The homogeneity of nb5 cone bipolar cells was considered with regard to several morphological parameters. In the final analysis, nb5 cone bipolar cells were determined to be the only narrow-field cone bipolar cell branching in S5 of the IPL. This conclusion is similar to that reached in the comprehensive EM reconstructions of type b cone bipolar cells, performed on the RC1 data set: none but CBb6 and wide-field cone bipolar cells branch primarily in S5 (Sigulinsky et al., 2020).

The discovery of a CaBP-IR, narrow-field cone bipolar cell, with axon terminals deep in the IPL, suggested the possibility of a selective marker for nb5 cells in the present study (Massey & Mills, 1996; Sharpe et al., 1993). Rod bipolar cells, labeled by anti-protein kinase C (PKC) ICC, have been used routinely as S5 fiducials, in order to gauge the stratification of other fluorescently labeled processes in the IPL, such as the axon terminals of CaBP-IR bipolar cells (e.g. Hoshi et al., 2009; Massey & Mills, 1996). The view that the “axonal endings of rod bipolars are…restricted to S5” (Strettoi et al., 1990, 1994) may have led to confusion, however, for Massey and Mills (1996) subsequently concluded that CaBP-IR bipolar cells were the same as those reconstructed by Strettoi and colleagues (1994), although the latter described their stratification as “narrowly confined to S4”. In fact, many robust presynaptic rod bipolar terminals are present at the level of the SAb band in the middle of S4 (e.g. Figure 12 in Famiglietti, 1991), while the majority of large lobulated processes of rod bipolar axon terminals do lie in S5.

Elegant studies by Massey and Mills (1996), combining ICC and intracellular staining and showing extensive dye coupling between AII amacrine cells and CaBP-IR bipolar cells provided supporting evidence for their correlation with the work of Strettoi and colleagues, who had shown that the S4 bipolar cell axon terminals make the “highest number” of gap junctions (4-5 per cell) with AII amacrine cells (Strettoi et al., 1994). Also seeming to support that conclusion is the evidence from RC1 that CBb6 bipolar cells have the most gap junctions (µ = 22.8 per cell) with AII amacrine cells. Nevertheless, CBb5 cells also make many such gap junctions (µ = 13.2 per cell) (Sigulinsky et al., 2020). Of relevance here is the observation in double labeling studies, with antibodies to glycine, in combination with antibodies to NK1R or CaBP, that 85% of NK1R-IR and 96.2% of CaBP-IR bipolar cells were immunoreactive for glycine, indicating the relative prevalence of gap junctions with AII amacrine cells (Kim et al., 2005), again suggesting correspondence between nb5, CBb6, and CaBP-IR cells, on the one hand, and nb1, CBb5, and NK1R-IR cells on the other.

I show here that no type of cone bipolar cell has axon terminals entirely confined to S4, but the only type that is mostly confined to S4 is the nb1 cone bipolar cell, not the nb5 cell (Figure 36c and d). Secondly, it is shown here that nb5 cone bipolar cells co-stratify with and likely make synaptic contact with class Ib2/ ON-Y ganglion cells, in agreement with the results of Zhang and colleagues (2005) using multi-labeling ICC and intracellular staining. Thirdly, nb5 cone bipolar cells exhibit the same stratification as CaBP-IR bipolar cells, with which they are surely identical. The nb5 axon terminals only occasionally extend up to the level of type b starburst amacrine cells, and ON-OFF (and sustained ON) DS ganglion cells, involving perhaps 10% of axon terminal processes. If nb5 cone bipolar cells are the correlates of CBb6 cone bipolar cells of RC1, it is surprising to learn that nearly half of the cone bipolar cell input to type b starburst amacrine cells derives from CBb6 bipolar cells, whereas none comes from CBb3 or CBb4 cells and very little from CBb3n cells (Marc et al., 2018).

It is even more troubling that Marc and colleagues (2018) find significant input from CBb6 cone bipolar cells to what they believe to be a transient ON DS/class IIb1 ganglion cell in RC1. Class IIb1 cells are found to branch on average 3.2 – 3.5 µm above the middle of the type b starburst amacrine cell dendritic band (Famiglietti, 2016), beyond the reach of nb5 bipolar cells. Those measurements were made in the visual streak, however, where the IPL is thickest. In the retinal periphery, where the IPL becomes progressively thinner (Famiglietti & Vaughn, 1981), and from which RC1 is likely derived, the IPL substrata are in closer proximity, and if CBb6 cells are promiscuous, rather than selective, in their synaptic connectivity, increased laminar proximity could be an explanation for CBb6 input to t-ON DS ganglion cells in the retinal periphery.

In a study of the synaptic connections of starburst amacrine cells in rabbit retina, accompanied by electrotonic modeling (Famiglietti, 1991), it was recognized that at least two different types of cone bipolar cell innervate starburst amacrine cells, one favoring spines near the cell body, above (for type a) or below (for type b) the respective starburst substrata. This duality of bipolar cell inputs has stimulated further modeling both in rabbit and mouse of directional selectivity in the starburst amacrine cell dendritic tree (Ding et al., 2016; Greene et al., 2016; Kim et al., 2014). If nb1 and nb5 cells correspond to CBb5 and CBb6 cells, respectively, then based upon the analysis of stratification in the present work, nb5/CBb6 synapses would be expected to occur more frequently on the proximal dendrites of type b starburst amacrine cells as they pass from the cell body, through S5, to stratify in S4. No information was provided, however, concerning differential input of CBb5 and CBb6, nor of any wb/CBbwf input to SAb cells (Marc et al., 2018). Again, increased laminar proximity in the retinal periphery could provide CBb6 bipolar cells access to the distal as well as the proximal dendrites of type b starburst amacrine cells.

It was noted above, that nb1 bipolar cells commonly give rise to small axon terminal branches in S1 and that these may form synaptic connections there with PA4 polyaxonal amacrine cells, presumed dopaminergic amacrine cells (Famiglietti, 1992c, 2002). In the present study, an nb5 cell was also shown to produce such a process, as were two wb cone bipolar cells. Hoshi and colleagues (2009), using multi-label ICC, showed that rare branches of CaBP-IR bipolar cells are formed in S1, and that both these branches and the axons shafts themselves form synaptic contact with dopaminergic amacrine cells, with the long spines of “bistratified diving” ganglion cells (aka type 3, tristratified, IVts3, ganglion cells in Famiglietti, 2020), and with M1 (OD1 ganglion cells in Famiglietti, 2020), intrinsically photosensitive ganglion cells, all known to give ON responses to light. Lauritzen and colleagues (2013) confirmed and extended this result in an EM study of the RC1 data set, finding that one third of type b cone bipolar cells of multiple types, give rise to axonal synapses throughout sublamina a, and also illustrate a rare S1 axonal branch of a CBb6 cell (Sigulinsky, personal communication). The axonal synapses of type b cells in S2 may be directed at dendrites of “bistratified, diving”/ IVts3 ganglion cells, significant portions of which lie in S2 as they travel (dive) from S1 back into sublamina b (Famiglietti, 2020).

#### 4.2.6 mb cone bipolar cells

The mb cone bipolar cells, newly identified here, but apparently encountered in a study of blue-OFF bipolar cells (Liu & Chiao, 2007), are paramorphic partners of the ma1 cell, previously identified (Famiglietti, 1981). In the present study, one example was found in close contact with the dendrites of a class IIb1 ganglion cell (Famiglietti, 2004a; Famiglietti, 2020), the morphological equivalent of the transient ON DS ganglion cell of rabbit retina (Ackert et al., 2009; Ackert et al., 2006; Famiglietti, 2016; Hoshi et al., 2011; Kanjhan & Sivyer, 2010). In a study of the RC1 data set, CBb4w (w = wide-field) bipolar cells were found to connect profusely with a large-bodied ganglion cell, partly included in RC1 and judged to be a transient ON directionally selective cell (Marc et al., 2018). The axon terminals of CB4w cells are reported to lie between 45 and 72% of the IPL’s thickness (Sigulinsky et al., 2020). The axon terminals of the mb bipolar cells lie between 46.7 and 80% (substrata 3β - 4γ) of the IPL (Figures 13 and 36c and d). The latter are also intermediate in size between the several narrow field types and the wide-field, “blue-ON” cone bipolar cells. For reasons of relative size and axon terminal stratification, mb cells and CBb4w cells constitute the most secure correlation of bipolar cells in the two studies (Table 1).

### 4.3 comparative analysis of bipolar cells: rabbit, mammalian, and other vertebrate retinas

It is of particular interest to correlate the results of the present classification of bipolar cells in rabbit retina with those of mouse and of monkey, for reasons related to their current frequent use as favorable model systems, enabling genetic manipulation on one hand, and close comparison with human retina on the other. A direct molecular genetic comparison of mouse and macaque retinal bipolar cells has in fact been performed (Peng et al., 2019; Shekhar et al., 2016). Additionally, comparison with ground squirrel retina offers insights into mammalian dichromatic vision shared with mouse and rabbit.

#### 4.3.1 mammalian cone bipolar cells: comparison of type a cells

In the case of type a bipolar cells, six kinds of cone bipolar cell were shown to be related, comparing mouse and monkey, based upon molecular data (Table 3). It is reassuring that a prior morphological comparison of mouse and macaque, with particular emphasis upon axon terminal field size and stratification (Tsukamoto & Omi, 2014) agrees with the single cell RNA sequencing studies (Peng et al., 2019), except for a crossing of DB1/BC2 and FMB/BC1a (Table 3).

**Table 3.** Mammalian Bipolar Cells: Comparison in Four Species.

| 1) Rabbit | 2) Mouse | 3) Macaque | 4) Ground Squirrel |  |
| --- | --- | --- | --- | --- |
| na1? | BC1A | FMB | C6 | 1b? |
| ND | BC1B | OFFx | C13 | X? |
| na2? | BC2 | DB1 | C8 | 1a? |
| ma1 | BC3A | DB3a | C5 | 2? |
| na3? | BC3B | DB3b | C1 | 3a? |
| na4? | BC4 | DB2 | C2 | 3b? |
| na4, nb1? | BC5A | DB4 | C10 | 5a? |
| nb2.1? | BC5B |  | (C3) |  |
| nb2.2? | BC5C | DB5 | C4 | (C3) |
| mb? | BC5D |  | C7 | 5b? |
| nb5?? | BC6, (BC8) | DB6 | C14, C15 | 6b? |
| nb1, nb5? | BC7 | IMB | C11 | 7b? |
| mb? | BC8 | GB/BB | C12 | 7a? |
| wb | BC9 | BB | C12 | bb |
| rbc | RBC | RB | C9 | rbc |

The comparatively large size of the related mouse BC3a and macaque DB3a bipolar cells accords with the dimensions and axon terminal stratification of the medium-wide field, rabbit ma1 cone bipolar cell. Moreover, hierarchical clustering shows BC3a cells are more distantly related to the other five type a cone bipolar cells (Hahn et al., 2023). Among narrow field bipolar cells, dimensions and axon terminal stratification of DB3b/BC3b and DB4/BC2 correlate well with those of rabbit na3 and na4 bipolar cells, respectively (Figure 36e). Less certain are the correlates of rabbit na1, na2, and ma2 cells with the remaining DB1/BC2 and FMB/BC1a bipolar cells. The rabbit ma1 bipolar cell is narrowly stratified at the S1/S2 border, suggesting limited and possibly selective connectivity with postsynaptic ganglion cells. On the other hand, its probable correlate in marmoset retina, DB3a, a homologue of DB3a in macaque, is connected to six types of ganglion cell (Masri et al., 2016), a fact signaling caution, when considering a relationship between narrow axon terminal stratification and selectivity of synaptic connections.

Rabbit retina contains wide-field, wa bipolar cells connected to blue cones (Famiglietti, 1981; Liu & Chiao, 2007). Similar bipolar cells have not yet been identified, however, in other mammals, and surprisingly scRNAseq has not revealed a corresponding cluster in a study encompassing 13 mammalian species (Hahn et al., 2023), although wa correlates may represent sparse populations, also sensitive to cell sorting. Macaque retina has FMB bipolar cells that contact blue cones only (Wool et al., 2019), but the FMBs of marmosets do not (Lee et al., 2005), nor does the cone-dominant ground squirrel retina have OFF bipolar cells selective for blue cones (Sher & DeVries, 2012). Missing from the scRNAseq analysis (Shekhar et al., 2016) is a mouse correlate of the wa (type a, blue) bipolar cell, but a possible morphological correlate in mouse, based upon large axon terminal field size and predominantly S1 stratification, has been termed BC”9o” (Sabbah et al., 2022), although its connection with blue cones is not established. Current electrophysiological studies do suggest that a wa homologue connected to blue cones could be present in mouse (Z. J. Zhou, personal communication).

In rabbit, the destination of wa bipolar cells is the type 3 bistratified (BS3) ganglion cell, branching in sublaminae a and b (Famiglietti, 2009), which has blue-OFF responses (Mills et al., 2014). Like rabbit, monkey has color-coded, bistratified ganglion cells, but their responses to blue light are opposite in polarity (Dacey & Lee, 1994; Dacey & Packer, 2003).

**Table 3**. Black typeface: secure molecular and morphological correlation; gray typeface: proposed morphological correlations. Molecular and morphological correspondence of mouse and macaque types (Hahn et al., 2023; Peng et al., 2019; Shekhar et al., 2016). The scRNAseq study of ground squirrel retina yielded 15 clusters (C), correlated with vertebrate “orthotypes” (column 2), named for mouse types (Hahn et al., 2023), but molecular correlations with morphological types remain to be determined. Ground squirrel morphological correlations here based principally on comparison with mouse axon terminal stratification and morphology (see Light et al., 2012).

#### 4.3.2 mammalian cone bipolar cells: comparison of type b cells

##### comparison of wide-field type b cells and potential for color vision

In all mammalian retinas, dichromatic color vision appears to require dedicated and unique pathways for transmission of short wavelength or blue photoreceptor signals to the inner retina, at least for the ON pathway. This was proposed for rabbit and cat retinas, both for ON and for OFF pathways (Famiglietti. 1981), as wb and wa bipolar cells, respectively, and was confirmed in later studies of rabbit (Liu & Chiao, 2007; MacNeil & Gaul, 2006; Mills et al., 2014).

Type b cone bipolar cells that are selectively connected to blue cones, like the wb bipolar cells of rabbit, have been identified in macaque retina (Kouyama & Marshak, 1992), and in mouse retina (Haverkamp et al., 2005). In most mammals studied, these are wide-field cells, but in some cases they may have narrow, converging dendritic fields like wa (blue-OFF) bipolar cells of rabbit, as in ground squirrel retina (Li & DeVries, 2006). Conservation of type selectivity between macaque and mouse has been demonstrated for blue-ON bipolar cells using single cell RNAseq (Peng et al., 2019).

Regarding the destination of wb cone bipolar cell and their homologues, the only blue-ON center cells encountered in a survey of rabbit retinal ganglion cells were ON-X (linear) cells (Caldwell & Daw, 1978). These have the morphology of large-bodied, IIb2 cells (Famiglietti, 2004a; Jensen, 1991), as confirmed by Mills and colleagues (2014), and it is clear that these are the principal recipient of wb cone bipolar cell output (Famiglietti, 2008; Mills et al., 2014). Blue-OFF responses can also be produced in some class IIb2 ganglion cells, derived from blue-ON pathways (Mills et al., 2014), presumably by the interposition of an inhibitory amacrine cell, as is likely in ground squirrel retina (Chen & Li, 2012; Sher & DeVries, 2012). In primate retina, large and small bistratified ganglion cells, and a unistratified ganglion cell all are blue-ON, yellow-OFF, color coded recipients of BB bipolar cell input (Dacey & Lee, 1994; Dacey & Packer, 2003). It has been challenging to find blue-selective ganglion cells in mouse retina. Sustained ON alpha (M4) ganglion cells show blue-ON responses in the mouse retina’s narrow transitional zone (Chang et al., 2013), where blue and green cone visual pigments are both expressed in single cones (Rohlich et al., 1994), and “ON-delayed”/EW73 exhibit a form of color selectivity (Hofling et al., 2024), but it may be that in mouse, true color-opponency first appears in the LGN (Mouland et al., 2021; Schwartz, 2021). In summary, the blue cone pathways have been studied both morphologically and physiologically in a variety of mammalian retinas. Although they all share blue-ON cone bipolar cells, branching in S4/S5 of the IPL, there are considerable differences in their postsynaptic connectivity.

More difficult to understand are the relationships among the other large field bipolar cells branching in sublamina b. In rabbit, the axon terminal of the mb cone bipolar cell stratifies at about the same level as reported for the giant bipolar cell of macaque (Joo et al., 2011; Tsukamoto & Omi, 2016), as well as the BC5d (x) bipolar cell of mouse (Figure 36e). The molecular studies show, however, that there is no correlate in monkey of the medium wide field mouse BC5d(x) bipolar cell. It was mentioned above that mb/CB4w co-stratify with and synapse on IIb1/transient ON DS ganglion cells in rabbit (Figure 32, and Marc et al., 2018). The axon terminals of mouse BC5d (x) cone bipolar cells, branching at the 50%-60% level of the IPL (Figure 36e), appear to co-stratify with the middle stratum of the weakly tristratified EW 7o/transient ON DS ganglion cell in mouse (Bae et al., 2018; Gauvain & Murphy, 2015; Goetz et al., 2022), but the morphology of the latter is very different from the large, unistratified IIb1/transient ON DS ganglion cell of rabbit retina, raising doubt about their homology.

The primate giant bipolar cell has the same stratification as BC5d(x), just above the SAb band (Figures 36e and 37a), as noted above, but it is related instead to the mouse blue bipolar cell (BC9), which is also related to and co-stratified with the wide field mouse BC8 bipolar cell, deep in the IPL (Figure 36e). In this regard, it is interesting that there is some evidence suggesting that the monkey giant bipolar cells are selective for medium or long wavelength cones (Joo et al., 2011) and there is strong evidence for green-selective bipolar cells in ground squirrel retina (Li & DeVries, 2006), while there is less for such bipolar cells in rabbit retina (Mills et al., 2014). On a morphological basis, rabbit mb bipolar cells could be related to monkey giant bipolar cells, but it is more likely that they correspond to mouse BC5d(x) cells. It would be interesting to know if mouse transient ON DS ganglion cells receive their principal bipolar cell input from BC5d(x) cells, as rabbit ganglion cell homologues apparently do from CBb4w/mb bipolar cells.

**FIGURE 37.**
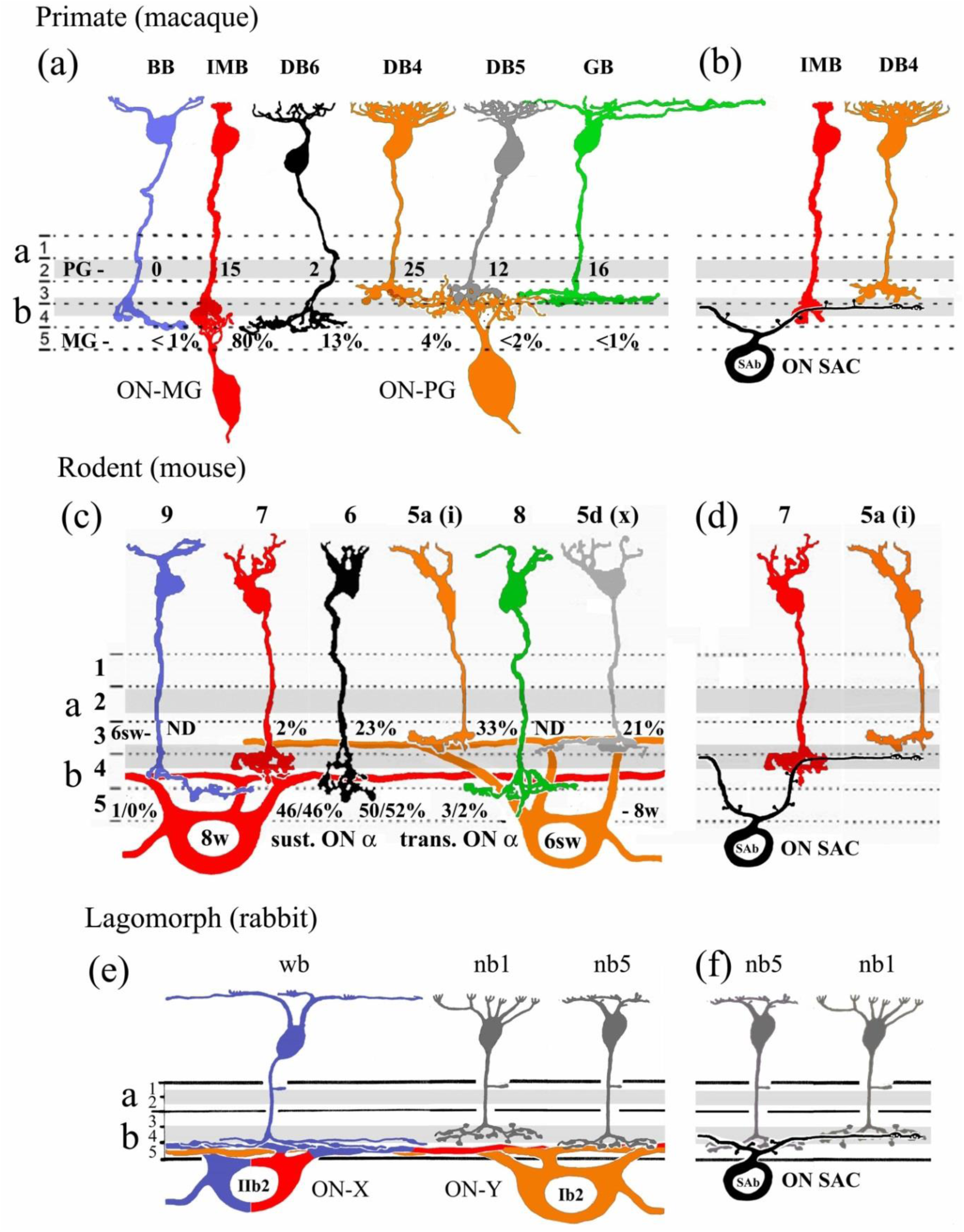
Homologous, parallel ON pathways from bipolar cells to ganglion cells (a, c, e) and to starburst amacrine cells (b, d, f) in sublamina b of the IPL in three classes of mammals. (Drawings refigured from Famiglietti (1981, 2004a) – for rabbit & and adapted for mouse; Tsukamoto and Omi (2017) – for mouse; Polyak (via Rodieck, 1988) and Tsukamoto and Omi (2016) – for macaque. “Orthology” of bipolar and ganglion cells in primate and mouse: (Hahn et al., 2023; Shekhar et al., 2016). (a) Bipolar-to-ganglion cell synaptic connectivity in macaque: Tsukamoto and Omi (2016) (parasol [PG] #s = number of ribbons); (c) connectivity in mouse (Kuo et al., 2024; Sabbah et al., 2022; Swygart et al., 2024). Proximal –to-distal topographic distribution of bipolar cell input to starburst amacrine cells: (b) macaque (Kim et al., 2022; Patterson et al., 2022), and (d) mouse (Greene et al., 2016). Identical color indicates homology, and blue indicates pathways from blue cones; homologies are inferred for ganglion cells in rabbit. (e) Rabbit wb, nb1 and nb5 cells are the only cone bipolar cells with axon terminal stratification deep enough to overlap that of IIb2 and Ib2 ganglion cells; rabbit appears to be an outlier in bipolar-to-ganglion cell connectivity for these ganglion cell homologues (e). In absolute size of ganglion cells, rabbit is more similar to mouse, but in relative size of the two ganglion cell homologues, rabbit is the reverse of mouse, and more similar to primates (and to carnivores). (f) In rabbit, type b starburst amacrine cells receive bipolar cell input from nb1 and nb5 cells and possibly from wb cells.

### comparison of narrow-field type b cells with axon terminal branches and synapses in sublamina a of the IPL

As noted above, branches of rabbit nb1 cone bipolar cells, presumably presynaptic to dopaminergic amacrine cells, have been demonstrated in S1 of the IPL (Famiglietti, 2002). Similar branches, and evidence for axonal synapses on ipRGCs and “bistratified-diving”/IVts3 ganglion cells in sublamina a has been presented for nb5/calbindin-IR bipolar cells (Hoshi et al., 2009), and EM reconstruction of type b bipolar cells in RC1 indicates that type b bipolar cells make synaptic contacts in sublamina a (Lauritzen et al., 2013). Similar results were obtained in mouse retina, where among type b cone bipolar cells, BC6 bipolar cells give rise to the majority of input to melanopsin-expressing ipRGCs in both sublaminae a and b (Dumitrescu et al., 2009; Sabbah et al., 2022). The presence of type b cone bipolar cell axonal synapses in S1 of sublamina a is not a new finding, however, for such so-called *en passant* axonal synapses in S1, as well as rudimentary branches have been demonstrated in type b bipolar cells in EM study of cat retina (Cohen & Sterling, 1990; McGuire et al., 1984). What is surprising is that such synapses occur at all levels of sublamina a, and in all varieties of type b cone bipolar cells in both rabbit (Lauritzen et al., 2013) and mouse (Tsukamoto & Omi, 2017). Bipolar cell input in S1 to intrinsically photosensitive ganglion cells and dopaminergic amacrine cells affords them ON input, but the specific role of functional input from nb1 and other type b bipolar cells to other inner retinal neurons in S2 and S3 remains to be determined. These type b bipolar cell inputs could be directed mainly along the whole extent of multistratified and diffuse ganglion cells, as may be the case for bistratified-diving/IVts3 ganglion cells, but the existence of such eccentric bipolar cell outputs foreshadows still more complexity in the ON/OFF “crossover connectome”.

#### 4.3.3 mammalian cone bipolar cells: parallel pathways directed to paramorphic ganglion cells

##### Historical perspective

Cajal studied the retinas of at least 25 species of five Linnean classes of vertebrates, beginning with birds (Cajal, 1889), the most complex, and finishing with mammals and teleost fishes, in a search for”simpler” retinas, in order to arrive at fundamental principles of retinal organization, mainly concerning the inner retina (Cajal, 1893). He regarded the mammalian retinas of ungulates, carnivores, and rodents as essentially similar and was inspired in their study to generalize about conduction pathways from bipolar cells to unistratified and to diffuse ganglion cells. Thinking about unistratified ganglion cells, he wrote “les voies de conduction, même les plus étroites et les mieux individualisées, sont toujours representées par un goupe de cellules bipolaire reliées a une, ou à quelques cellules ganglionaires seulement” (Cajal, 1893, p.220); transl.: *the conduction pathways, even the narrowest and the most specialized, are always represented by a group of bipolar cells connected to one or only a few ganglion cells*.

As much as he discerned and depicted the stratification of amacrine and ganglion cells, Cajal gave priority to bipolar cells in the stratification of the IPL, suggesting that in peripheral retina, where the number of bipolar cell body layers is reduced (i.e. decreased in density), that the number of strata in the IPL is reduced from five to three. In this conjecture he appears to have been incorrect, for immunocytochemical staining shows that generally in peripheral retina strata become more compressed rather than fewer (Famiglietti & Vaughn, 1981), and amacrine cells are primary in forming the strata of the IPL (Kay et al., 2004). Nevertheless, this narrowing does have the effect of bringing bipolar cell axon terminals of different types into closer laminar proximity (see Discussion above). In summary, these conclusions concerning bipolar-to-ganglion cell transmission prefigure the concept of parallel pathways that has guided much retinal neurobiology in the recent past.

In the modern era, a quest for the meaning of stratification in the IPL, and revelation of parallel pathways in the inner retina, resumed in discovery of the bisublaminar dendritic architecture of ganglion cells transmitting ON and OFF photic responses in sublaminae a and b of the IPL of cat and carp (Famiglietti et al., 1977; Famiglietti & Kolb, 1976; Nelson et al., 1978). Retrospectively, similar bisublaminar organization of ganglion cells could be discerned in Cajal’s illustrations from dog retina (Cajal, 1893), and in Polyak’s illustrations of midget ganglion cells in monkey retina (Polyak, 1941). In carnivores and in monkey, the parallel pathways were conducted through”paramorphic” pairs of ganglion cells of similar appearance but different bisublaminar stratification (Famiglietti & Kolb, 1976). The elucidation of these paramorphic ganglion cell types in rabbit retina was more challenging (Famiglietti, 2004a), and escaped full understanding in rodents until recently (Baden et al., 2016; Bae et al., 2018; Krieger et al., 2017)

Now, with the power of single cell RNA sequencing, it is possible to compare ganglion cell types across a number of vertebrate species (Hahn et al., 2023). Furthermore, it is possible to discern homologs of two sets of paramorphic ON and OFF midget ganglion cells and ON and OFF parasol ganglion cells, characterized physiologically first in cats and rabbits as parallel sets: ON-center and OFF-center pairs of X and Y cells (Caldwell & Daw, 1978; Cleland & Levick, 1974; Enroth-Cugell & Robson, 1966). In mouse, these occur as paramorphic ON and OFF sustained and transient “alpha” cells, respectively (Krieger et al., 2017), although the “paramorphic” ON or OFF pairs appear less similar to each other morphologically and physiologically in mice (e.g. Bae et al., 2018; Goetz et al., 2022), than those in primates, carnivores and rabbits.

##### Homologous ganglion cells

Unlike the narrow-field midget and small-to-medium size parasol ganglion cells of primate (macaque) retina, their homologues in mouse and rabbit are wide-field ganglion cells even in central retina (Figure 37). The stratification of comparable ganglion cells in sublamina a are similar in all three animals. Midget, sustained alpha, and IIa ganglion cells branch primarily in S1, above the type a starburst amacrine (SAa) cell band of dendrites, while parasol, transient alpha, and Ia2 ganglion cells branch primarily in S2 below the band (Bae et al., 2018; Famiglietti, 2004a), although in monkey, there is considerable overlap with the SAa band (Kim et al., 2022).

In sublamina b, however, the situation is different (Figure 37). Primate midget and parasol ganglion cells overlap the sublamina b band of starburst amacrine (SAb) cell dendrites, the former extending below the band and the latter extending above the band. Similarly, in mouse, sustained ON alpha/EW8w ganglion cells and transient ON alpha/EW6sw cells lie below and above the band, respectively, but exhibit less overlap with the SAb band. The peak density of EW8w cell dendrites occurs at about 78% and of EW6sw cells at 56.5%, while the middle of the SAb band lies at 62% (Figure S5C in Bae et al., 2018). In contrast to monkey and mouse, both ON-X/IIb2 and ON-Y/Ib2 cells of rabbit retina lie, essentially co-stratified, in the middle of S5α at about 83% of the IPL’s depth (Figures 27 and 37 and Famiglietti, 2004a), whereas the middle of the SAb band lies at 67-70% of the depth of the IPL in rabbit retina (Famiglietti & Tumosa, 1987).

##### Parallel bipolar cell pathways to ganglion cells

In monkey and mouse, where homologies have been established, parallel pathways from specific bipolar cell to ganglion cell types can be appreciated, particularly in sublamina b (indicated by identical colors in Figure 37a-d), and stratification of bipolar cell axon terminals with the main branching of postsynaptic ganglion cell dendrites is overlapping if not coextensive.

In sublamina a of the IPL, the stratification of the cone bipolar axon terminals of type a cells in monkey, mouse, and rabbit overlap considerably (Figure 36). Where the outputs of type a bipolar cells have been studied in mouse, particularly in relationship to ipRGCs, their synapses on ganglion cells were correlated primarily with stratification depth, and where overlapped in depth, showed limited output specificity, with BC3a and BC3b providing about 40% each of the bipolar cell input to transient OFF alpha cells, and most of the remaining bipolar cell input deriving from BC4 cells (Sabbah et al., 2022). Sustained OFF alpha cells were not studied. It was suggested above that mouse BC3a and rabbit ma1 cells are homologous, and in rabbit, these together with na2, nb3 and nb4 may all provide input to class Ia2 ganglion cells, mainly in S2, while na1 and ma2 are best positioned to provide input to class IIa ganglion cells, mainly in S1. The primate homologue of mouse BC3a cells, the DB3a cell, is more specialized in its output, providing 58%of the bipolar cell input to parasol cells, and the monkey homologue of the BC1a bipolar cell, the FMB cell, more specialized still, provides 80% of the bipolar cell input to type a midget ganglion cells (Tsukamoto & Omi, 2015).

In sublamina b of the IPL in mouse retina, the axon terminals of five types of cone bipolar cell: 5a(i), 5b(o), 5c(t), 5d(x), and the outer part of 6, branch at about the same level (Figure 36e), coinciding with the dendritic branching of transient ON alpha cells, whereas the inner part of 6 and 7, and the whole of 8, and 9 (Figure 36e) overlap the dendritic branching of sustained ON alpha ganglion cells (Figure 37c; see Figure S5C in Bae et al., 2018). In sublamina b of the IPL in rabbit retina, there are only three types of cone bipolar cell with axon terminal processes capable of reaching the dendrites in S5α of the two presumed homologs, IIb2 and Ib2 ganglion cells: the co-stratified wb and nb5 bipolar cells, and a few descending processes of nb1 bipolar cell axon terminals (Figure 37e).

##### Bipolar-to-ganglion cell connectivity in sublamina b: similarities

In primate retina, the midget system of paired bipolar and ganglion cells has been viewed as an extreme specialization, affording the possibility of exclusive connections between a single cone and a single ganglion cell, at least in the fovea centralis, mediating both high visual acuity and color coding. Such connections have been examined in serial-section, 3D-EM-reconstruction of macaque perifovea (Tsukamoto & Omi, 2016). Type b midget ganglion cells branching primarily in S4, at a mean depth of 68% (Kim et al., 2022), received a total of 93% of their input from two types of bipolar cell, 80% from IMB cells and 13% from DB6 cells, both of which branch mainly in S4, less than 1% from co-stratified BB cells, and about 6% from DB4, DB5, and GB, which branch mainly in S3 (Figure 37a). Primate type b parasol ganglion cells, branching in S3 and S4, at a mean depth of 61% (Kim et al., 2022), receive input from five types of bipolar cell, including IMBs, a slight preponderance coming from DB4 bipolar cells (25 of 70 synapses counted), and with significant input from giant bipolar cells, but none from BB cells (Tsukamoto & Omi, 2016). These data favor the view that the primate midget system is more highly specialized than the parasol system in terms of connectivity as well as function, and that neither participates in the blue-ON pathway.

In mouse retina, the homolog of the type b midget ganglion cell, the sustained ON alpha cell, receives 96-98% of its bipolar cell input from the two bipolar cell homologs of DB6 and IMB: BC6 and BC7, respectively (Figures 37a and c), and no more than 1% from BC9 (Sabbah et al., 2022; Swygart et al., 2024). The synaptic connections of the mouse homolog of the type b parasol ganglion cell, the recently described transient ON alpha cell (Krieger et al., 2017), involve BC6 and all four kinds of type 5 bipolar cells, with BC5a(i) cells, the homologs of monkey DB4 cells, predominating (Kuo et al., 2024); similarly the connections of the transient OFF alpha cell has been shown to receive input from the three type a bipolar cells: BC3a, BC3b, and BC4, that share its stratification (Sabbah et al., 2022). These data support the idea of a traceable, derivative, mammalian origin of the midget system (Hahn et al., 2023), and a related, but less specialized and more transient parallel system of generally larger paramorphic ganglion cells among mammals.

##### Bipolar-to-ganglion cell connectivityin sublamina b: differences

Comparing mouse and monkey, some lack of specialization in mouse is suggested by the finding that the ratio of the two principal homologous bipolar cell inputs are 80:13 (IMB:DB6) for midget and 46:50 (BC7:BC6) for sustained ON alpha ganglion cells (cf. Figure 37a and c). Furthermore, functional similarities of the two homologous ganglion cells, in particular their sustainedness, may be derived differently in regard to their bipolar cell inputs. Midget bipolar cells are very sustained in their responses (Dacey et al., 2000), but the sustained responses of sustained ON alpha cells in mouse apparently derive not from the IMB homolog, BC7, but from BC6 bipolar cells. This sustainedness is reportedly due not to the BC6 cells’ moderately sustained responses, measured at the cell body, but to enhanced availability of releasable neurotransmitter at their larger synaptic ribbons (Kuo et al., 2024). On the other hand, there is the seemingly opposite finding of comparatively more sustained release at ribbon-free active zones, directly visualized in goldfish bipolar cell axon terminals (Midorikawa et al., 2007). Quantitative scrutiny of active zones could be moot, however, if small numbers of bipolar cell inputs can have outsized effects, as proposed for directionally tuned mouse BC7 bipolar cell axon terminals, that are few but perhaps powerful inputs to ON-OFF DS ganglion cells (see below) (Matsumoto et al., 2021).

Comparing rabbit with mouse and monkey, there are sound physiological and morphological data for rabbit, but EM evidence for synaptic connectivity between bipolar and ganglion cells is lacking, as is comprehensive molecular genetic evidence. In rabbit, only two types of bipolar cell, wb/blue and nb5, stratify with the co-stratified, putative homologs of midget and parasol ganglion cells: IIb2 and Ib2, respectively. Electrophysiological recordings and intracellular staining make clear that IIb2 cells are dominated by blue cone input in the receptive field center (Caldwell & Daw, 1978; Mills et al., 2014). That evidence does not rule out some input, however, from nb5 bipolar cells to IIb2 ganglion cells in rabbit. There is little doubt that wb bipolar cells are homologous with monkey BB and mouse BC9 bipolar cells, rather than midget or BC7 bipolar cells. The preferred target of nb5 bipolar cells appears to be Ib2/ON-alpha ganglion cells (Figure 27 above, and Zhang et al., 2005). Considering homologies, one may ask: is nb5 related to mouse BC6, which has significant input to transient ON alpha cells (Figure 27c), even though its homolog in monkey, DB6, has negligible input to parasol ganglion cells (Figure 37a)? That homology seems unlikely. On the other hand, is nb5 related to monkey IMB which has significant input to parasol ganglion cells, even though its homolog in mouse, BC7, has negligible input to transient ON alpha ganglion cells (Figure 37c)? The answer is probably not. Homologous DB4 and BC5a(i) bipolar cell provide the predominant input to parasol and transient ON alpha cells, respectively; is nb5 homologous with these? Molecular genetic analysis would be more decisive than EM-level study of bipolar cell outputs in resolving this issue.

The differences in connectivity of primate midget and parasol pathways, compared to their homologs in mouse, seem to be largely quantitative, despite large differences in their roles in visual perception and behavior in mice and monkeys. Beyond the scope of this comparative perspective on paramorphic ganglion cells and parallel pathways conducted through bipolar cells, differences in color vision are manifest at the level of ganglion cells in these two species, as noted above. When rabbit is brought into the comparison, the fusion of blue-ON pathway with the IIb2/ON-X pathway shows that evolutionary pressure can induce rerouting that associates or dissociates bipolar to ganglion cell connections, probably in order to refashion photoreceptor inputs to ganglion cells, serving higher visual centers and differing animal behaviors. If the finding of increased sensitivity of blue cones to light decrements (Baden et al., 2013) holds in rabbit retina, one can appreciate the advantage of linking blue pathways to ganglion cells with large cell bodies and large axons with rapid conduction velocities in order to facilitate escape from airborne predators in daylight.

#### 4.3.4 mammalian cone bipolar cells: pathways to starburst amacrine cells for directional selectivity

##### Comparison of narrow-field type b cells, in relationship to starburst amacrine cells and directional selectivity

A comparison of narrow field type b cone bipolar cells in rabbit, macaque, and mouse is interesting on account of the apparently selective roles of some bipolar cells in visual feature extraction, originally demonstrated in the responses of rabbit retinal ganglion cells (Barlow et al., 1964; Caldwell & Daw, 1978; Levick, 1967). The anatomical basis of selectivity for direction of motion in several types of retinal ganglion cell is instructive for understanding the complexity of bipolar cell function.

The co-stratification of starburst amacrine cells and putative ON-OFF DS ganglion cells (Famiglietti, 1983) pointed to the importance of these radially organized neurons in the asymmetric responses of DS ganglion cells. DS responses have been demonstrated in the distal dendritic zone of starburst amacrine cell boutons (Euler et al., 2002; Lee et al., 2010; Lee & Zhou, 2006), but the likely combination of synaptic connectivity, morphological features, and intrinsic properties underlying such responses has been challenging to disentangle, and appears to include reciprocal GABAergic inhibition among SA cells (Lee & Zhou, 2006), and positional input to individual SA cells from sustained and transient bipolar cells, so-called “space-time” wiring, based upon EM reconstruction of bipolar cell contacts with starburst amacrine cells in mouse retina (Kim et al., 2014).

The first evidence for differential bipolar cell input to SA cells was provided in EM reconstructions of rabbit retina, where it was also noted in electrotonic modeling, based upon measured morphological parameters, that sustained bipolar cell inputs in the proximal dendritic zone of SA cells could have a significant effect on the output of distal synaptic boutons (Famiglietti, 1991). EM studies in mouse retina, stained to reveal synapses, have generally confirmed the differential input of BC7 input to the proximal dendritic zone and BC5 cells to more distal regions of the SA cell dendritic tree, as proposed by Greene and colleagues (2016), showing minor differences between their mouse data and the rabbit data, related to eye size (Ding et al., 2016). Problematic for the “space-time” wiring model, however, is that BC7 bipolar cells are not really sustained in their responses (Ichinose et al., 2014), but have both transient and sustained components (Z. J. Zhou, personal communication). Nevertheless, recent studies measuring glutamate release onto SA cells, assuming that its effect are localized at synapses, indicates that proximal bipolar inputs are more sustained than distal ones (Srivastava et al., 2022).

Further complicating matters are conflicting studies testing whether bipolar cell axon terminals of either BC5 or BC7 cells may be directionally selective themselves (Chen et al., 2014; Matsumoto et al., 2021). In the later study, a subset of boutons of a single BC7each exhibit an individual directional preference. Moreover, evidence is found for complex interactions: BC7 bipolar cells driven by extrasynaptic ACh from starburst boutons and inhibited by directionally selective wide-field GABAergic amacrine cell inputs that are modulated by inhibitory input from SA cells. As key elements in DS neural circuitry, BC7 bipolar cells provide surprisingly little input to ON-OFF DS ganglion cells (Sabbah et al., 2022). Nevertheless, it is argued that only a few of such inputs, aligned with the preferred direction of the recipient DS ganglion cell are sufficient, depending upon the array of postsynaptic glutamate receptors and potentiation by ACh (Matsumoto et al., 2021). The concentration of BC7 cells in the proximal dendritic zone of SA cells is consistent with their level of axon terminal stratification at the level of SAb primary dendrites, but if their individual boutons are already selectively tuned to a cardinal direction (Matsumoto et al., 2021) and control activity in a SAb dendritic sector (Euler et al., 2002; Famiglietti, 1991), it is not entirely clear why distal bipolar cell inputs are required except as potentiators.

Recent work in primate retina indicates that SAb cells receive input from three types of bipolar cell: IMB (invaginating midget bipolar) cells that exhibit sustained responses (Dacey, 2000), primarily in the proximal dendritic zone, and more transient DB4 and DB5 bipolar cells principally in more distal regions (Kim et al., 2022), with DB4 cells predominating (Patterson et al., 2022),in accord with their levels of axon terminal stratification (Figure 36e). It appears that a key difference between the two is the presence of voltage-gated sodium channels in the distal portion of the axon, absent in IMB cells (Puthussery et al., 2013). Interestingly, it appears that the centrifugal preference underlying the motion sensitivity of SAb cells in monkey is dependent upon the surrounds of bipolar cells generated in the OPL (Kim et al., 2022).

##### Gap junctions afford widespread influence of mouse BC6 bipolar cells, but probably not on DS mechanisms

Another complicating factor, influencing the responses of bipolar cells differentially, is the presence of slower components transmitted through gap junctions between bipolar cells. While the amount of coupling among cells of the same type varies from one bipolar cell type to another, and appears to vary for homologues from one species to another, specific cross-coupling between different types may be more significant, when considering mouse (Tsukamoto & Omi, 2017) and rabbit (Sigulinsky et al., 2020).

Mouse BC6 bipolar cells are in an intermediary position to control a number of processes in the IPL, with the most frequent branches in S1, like nb1 cells and to a lesser extent nb5 cells in rabbit, making a majority of the bipolar cell synapses with ipRGCs throughout the IPL (Sabbah et al., 2022), and making the largest number of gap junctions with AII amacrine cells (Tsukamoto & Omi, 2017). In rabbit, bipolar cells with the deepest axon terminals (CBb5, 6, and wf) make the most gap junctions with AII amacrine cells, whereas type b bipolar cells branching near the middle strata (especially CBb4 and 4w) have more heterocellular bipolar cell gap junctions, and more gap junctions overall (Sigulinsky et al., 2020).

The axon terminals of BC6 cells are broadly stratified in S4 and S5 (Figures 36e and 37c), where both nb1 and nb5 cells branch, but BC6 cells make little contact with SAb cells (Greene et al., 2016), despite having proportionally the greatest synaptic output to amacrine cells (Tsukamoto & Omi, 2017). If nb1 and nb5 cells are the CBb5 and CBb6 cells, respectively, of the RC1 rabbit data set, making the majority of bipolar cell synapses onto SAb cells (Marc et al., 2018), then neither is a match for BC6 cells, or for homologous monkey DB6 cells. Thus it seems that BC6 bipolar cells play no role in directional selectivity, nor do they have a clear homologue among rabbit bipolar cells.

##### Multiplexed pathways of bipolar cells influencing DS complicate functional comparison of homologues

Mouse narrow-field BC5 bipolar cells of three different types: BC5a(i), BC5b(o), and BC5c(t), functionally among the most varied type b bipolar cells (Chen et al., 2014; Ichinose et al., 2014), make gap junctions with each other (Tsukamoto & Omi, 2017). All three BC5 types branch at about the same level in the IPL (Figure 36e), and all three make synaptic contacts with SAb cells (Ding et al., 2016; Greene et al., 2016). BC5a(i), the homolog of monkey DB4, is more concentrated in the periphery of the SAb dendritic field, but has less input (30%) than BC5c(t) cells (46%) (Ding et al., 2016), which also provide the most bipolar cell input to ON-OFF DS ganglion cells in sublamina b (Sabbah et al., 2022). In addition, BC5a(i) also performs a different task for DS ganglion cells, providing vertical orientation selectivity, involving selective connections with oriented dendrites of wide-field amacrine cells to DS ganglion cells of all preferred directions (Chander et al., 2024). Notably, BC5a(i) cells have divided responsibilities, not only in service of orientation and directional selectivity, but also as the principal driver of transient ON alpha cells (see above and Figure 37c). Comparing mouse and monkey, taking into account molecular similarity, axon terminal stratification, distribution in the starburst dendritic field, and connectivity, the correlations are secure between BC7 and IMB cells, central in the SAb dendritic field, and between BC5a(i) and DB4 cells, peripheral in the SAb dendritic field.

In a comparison of rabbit with mouse and monkey bipolar cells in sublamina b, the question arises as to which are the more important criteria for classification: field size, axon terminal stratification, identity of postsynaptic neurons, gap junctional coupling, or types of synapses with cones. Regarding synapses with cones, IMB and blue bipolar cells in monkey are the only monkey bipolar cells in which virtually all dendritic terminal clusters of digitiform appendages are fully invaginating, ribbon-related terminals (Tsukamoto & Omi, 2016). In similar fashion, as demonstrated in the present study, nb1 cells are conspicuous among all rabbit bipolar cells in having narrow clusters of vertically rising digitiform terminal appendages, resembling those of IMB cells (Figure 11, above). On that account, nb1 bipolar cells might be homologues of IMB and BC7 bipolar cells in monkey and mouse.

Regarding input to starburst amacrine cells, evidence in the present study of rabbit favors the view that nb1 cells are the primary source of bipolar cell input to SAb cells, as well as ON-OFF DS ganglion cells, based upon co-stratification with the main band of SAb dendrites (Figures 12 and 13, above). Doubts were raised in the present study about the proximity of nb5 bipolar cells to the main band of SAb dendrites that lie in the middle of sublamina b, and the likelihood of a large amount of input to starburst amacrine cells (Figures 26-29, above). The nb5 cells are positioned, however, to contact the more proximal portions of type b starburst amacrine cells. Their morphology, stratification, and to a degree their connectivity (Marc et al., 2018), favor a relationship, when comparing the three species, among and DB4, BC5a(i) [or BCc(t)], and nb1/CBb5, on the one hand, and IMB, CBb7, and nb5/CBb6 on the other (Figure 37b, d, and f). It is difficult to discern relationships between the rabbit narrow-field types: nb2.1, nb2.2, nb3.2, and nb4, and mouse BC5 subtypes that branch at about the same level of the IPL (cf. Figure 36d and e).

The greater similarity of bipolar cell types in relationship to type b starburst amacrine cells between monkey and mouse, as compared to rabbit, is surprising, considering the shorter phylogenetic distance between the latter two. Further EM studies of connectivity at both axonal and dendritic ends of rabbit bipolar cells would be helpful in resolving some discrepancies noted here, but molecular genetic studies will be required to determine homologies among these mammalian species.

#### 4.3.5 mammalian cone bipolar cells: general conclusions

It has been a compelling idea that parallel pathways in the visual system originate in the responses of bipolar cells, conveying ON and OFF responses (Famiglietti et al., 1977; Famiglietti & Kolb, 1976; Nelson et al., 1978), photoreceptor selective chromatic responses, and sustained or transient responses (Awatramani & Slaughter, 2000; Ichinose et al., 2014; Kuo et al., 2024), as well as linear and nonlinear responses (Demb et al., 2001) to a variety of retinal ganglion cells. Rod versus cone pathways, the latter comprising the nearly universal mammalian blue cone selective pathway (Jacobs, 2013; Miyagishima et al., 2014) and green cone selective pathways, documented in mouse (Breuninger et al., 2011) and ground squirrel (Li & DeVries, 2006), highlight a key role of bipolar cells. The presence of 22 bipolar cell types in a galliform retina, which has eight types of photoreceptor (Yamagata et al., 2021), indicates that a degree of bipolar cell diversity is required to channel photoreceptor-selective responses to ganglion cells.

Photoreceptor types in rabbit and mouse are few, however, and the presence of about 15 bipolar cell types in each indicates that chromatic responses in ganglion cells do not account for such diversity. Macaque retina, with an additional visual pigment, in fact has fewer bipolar cell types than mouse (Peng et al., 2019) or rabbit. The roles and interactions of bipolar cells are nearly as diverse as those once attributed to amacrine cells. Perhaps a better correlation, at least among the species addressed in the above correlations, though a rough measure, is the number of ganglion cell types in each, increasing, like the number of bipolar cell types, from macaque to mouse to rabbit retina (Famiglietti, 2020, 2024; Shekhar et al., 2016).

## ACKNOWLEDGEMENTS

I thank E. C. Siegfried, D. McGlone, and B. R. Ferguson for assistance preparing Golgi-impregnated retinas, and S. J. Sharpe for assistance with the immunocytochemistry and fluorescence photomicrography. C. L. Sigulinsky clarified several points regarding published data from RC1, in helpful discussions, accompanied by illustrative material from RC1. Z. J. Zhou and G. W. Schwartz provided insight into aspects of the electrophysiological studies and identification of bipolar and ganglion cells, pertaining to the comparative analysis of rabbit and mouse bipolar cells. I thank J. R. Sanes and K. Shekhar for comments and clarifications regarding the derivation of “orthologs” from scRNAseq data, supporting the comparison of parallel pathways in several species. Data analysis, graphics, and camera lucida drawings were performed, designed, and made by E. V. Famiglietti. This research was supported in part by P.H.S. Award R01EY03547, the Alfred P. Sloan Foundation, the Alberta Heritage Foundation for Medical Research, the Medical Research Council of Canada, and the Natural Sciences and Engineering Research Council of Canada.

## Appendix Figures A1-A15

**FIGURE A1.**
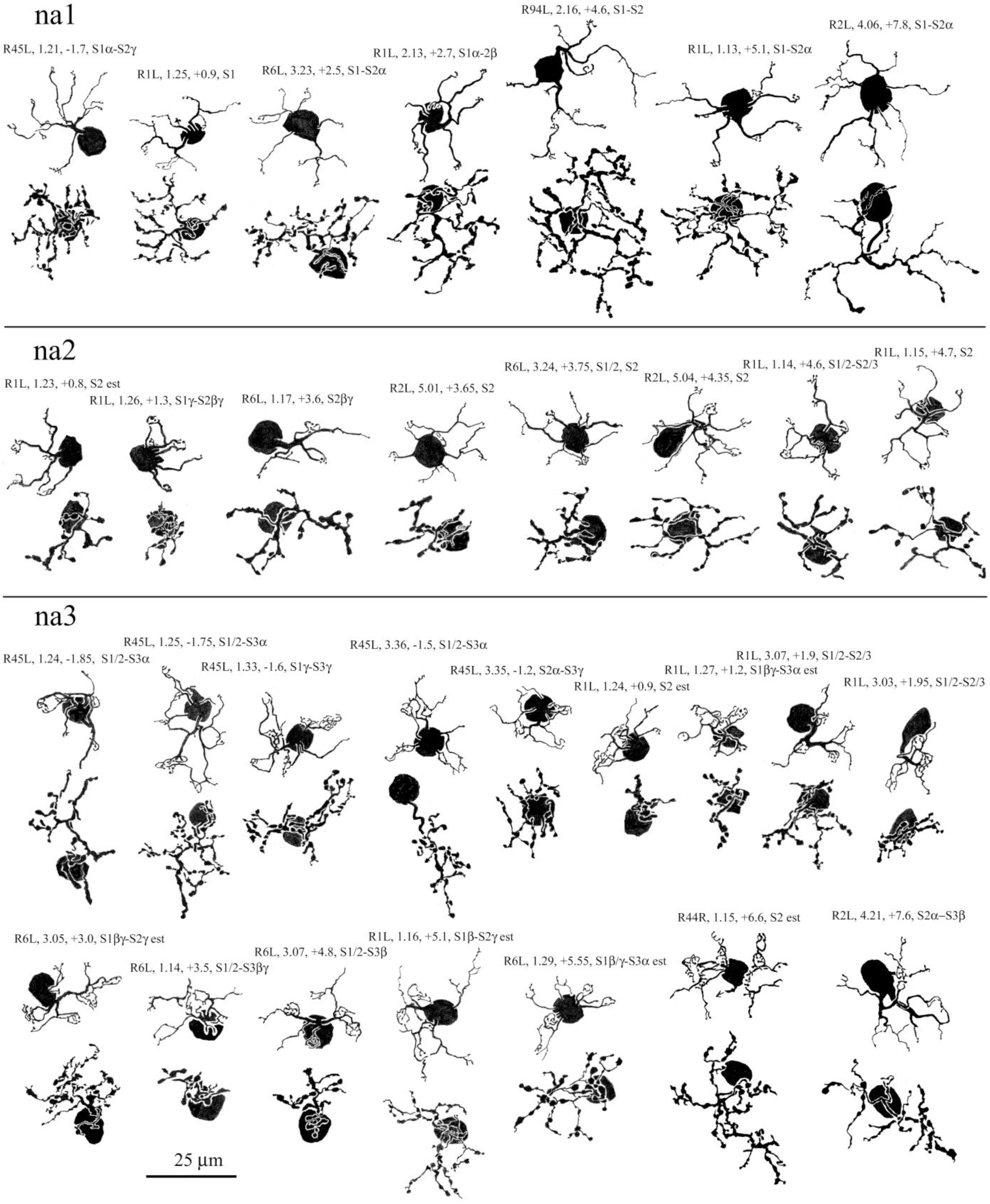
Camera lucida drawings of narrow field, type a cone bipolar cells: na1, na1, and na3, from six retinas. Dendritic trees above, axon terminal arborizations below in this and subsequent figures. Calibration bar applies to all cells in this figure. Similarly, calibration bars and subsequent appendix figures apply to cells in each figure

**FIGURE A2.**
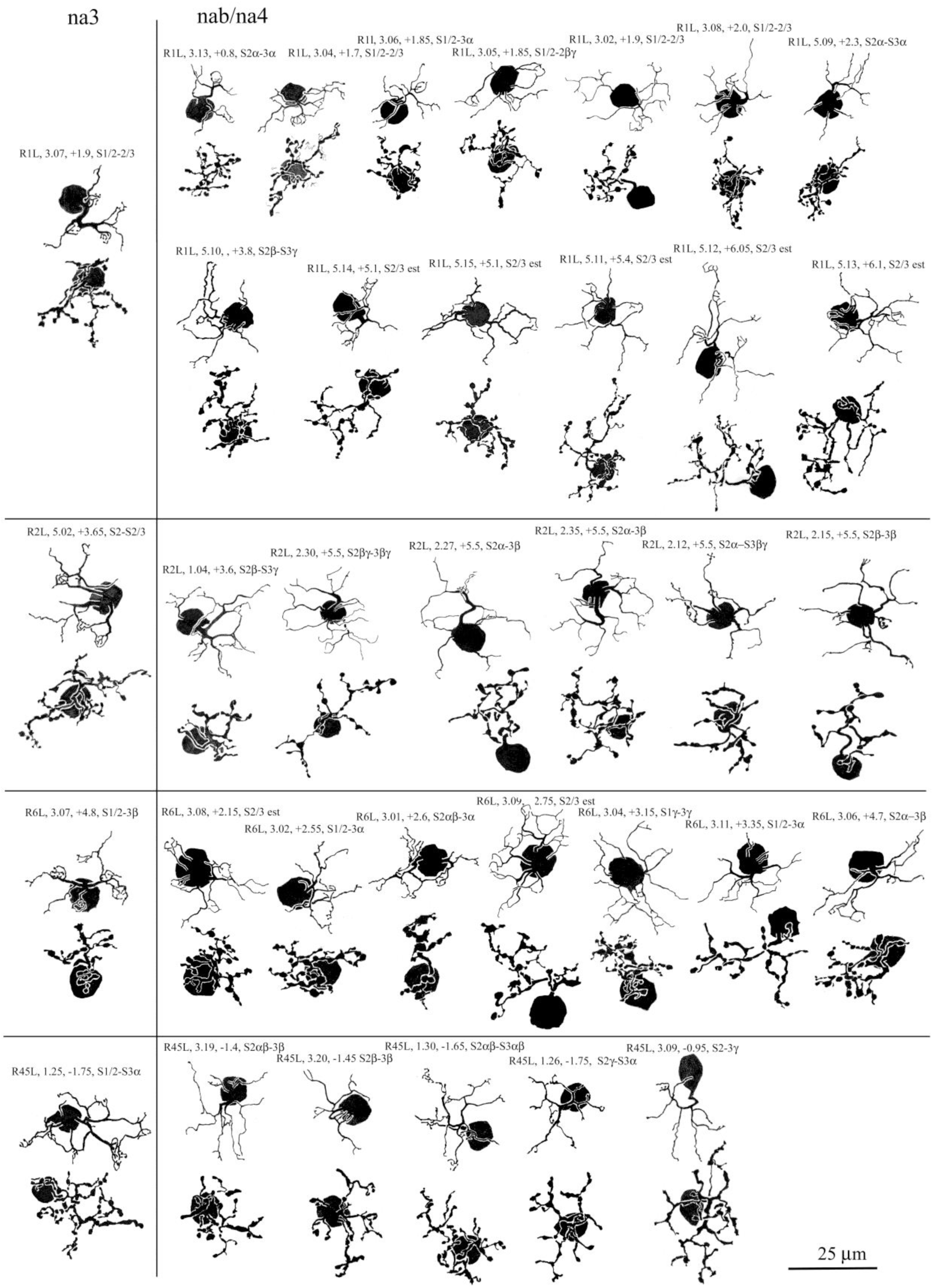
Camera lucida drawings of narrow field, type a cone bipolar cells: na4 (formerly nab), right, compared with na3 cells, left

**FIGURE A3.**
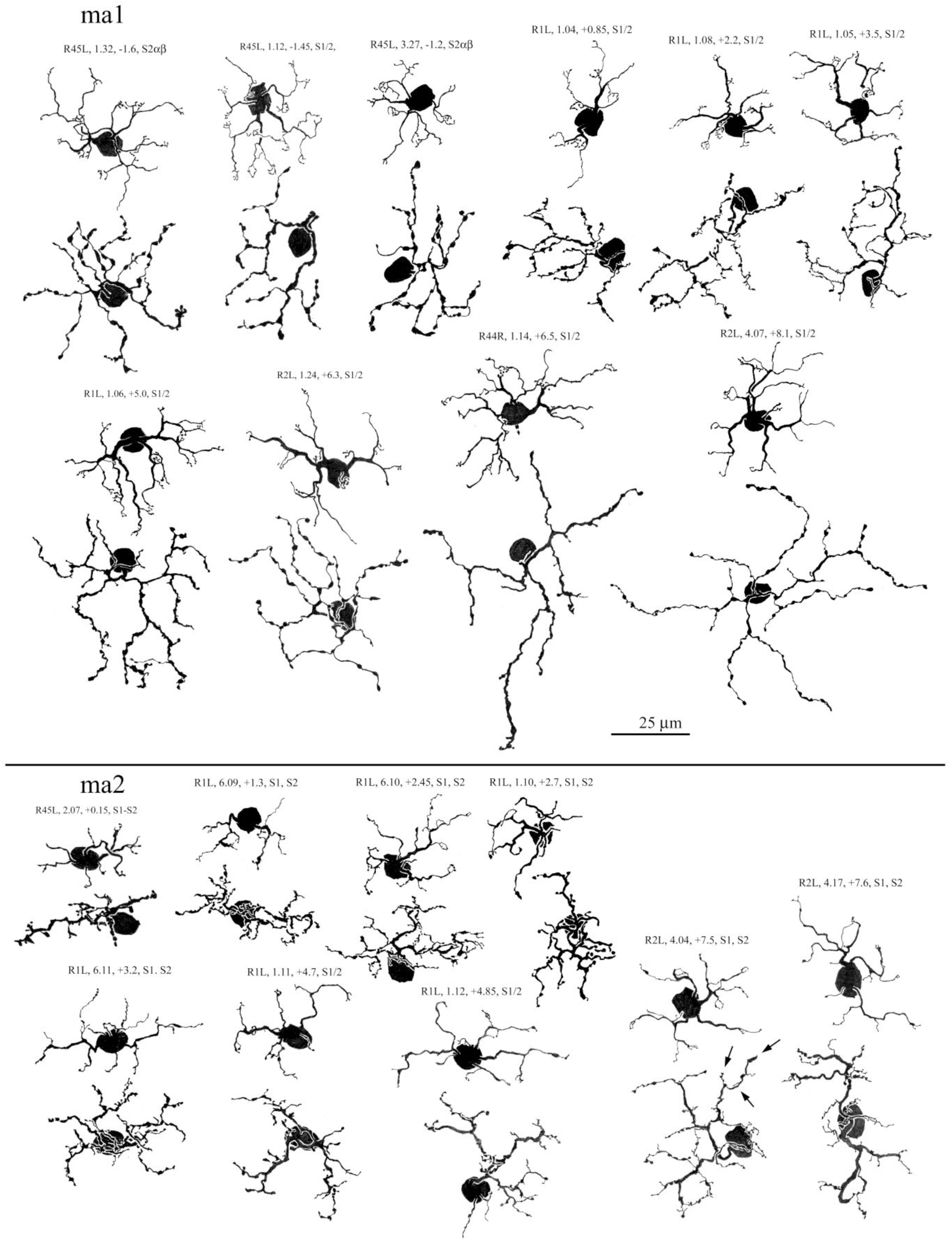
Camera lucida drawings of medium wide field, type a cone bipolar cells: ma1 and ma2 cells

**FIGURE A4.**
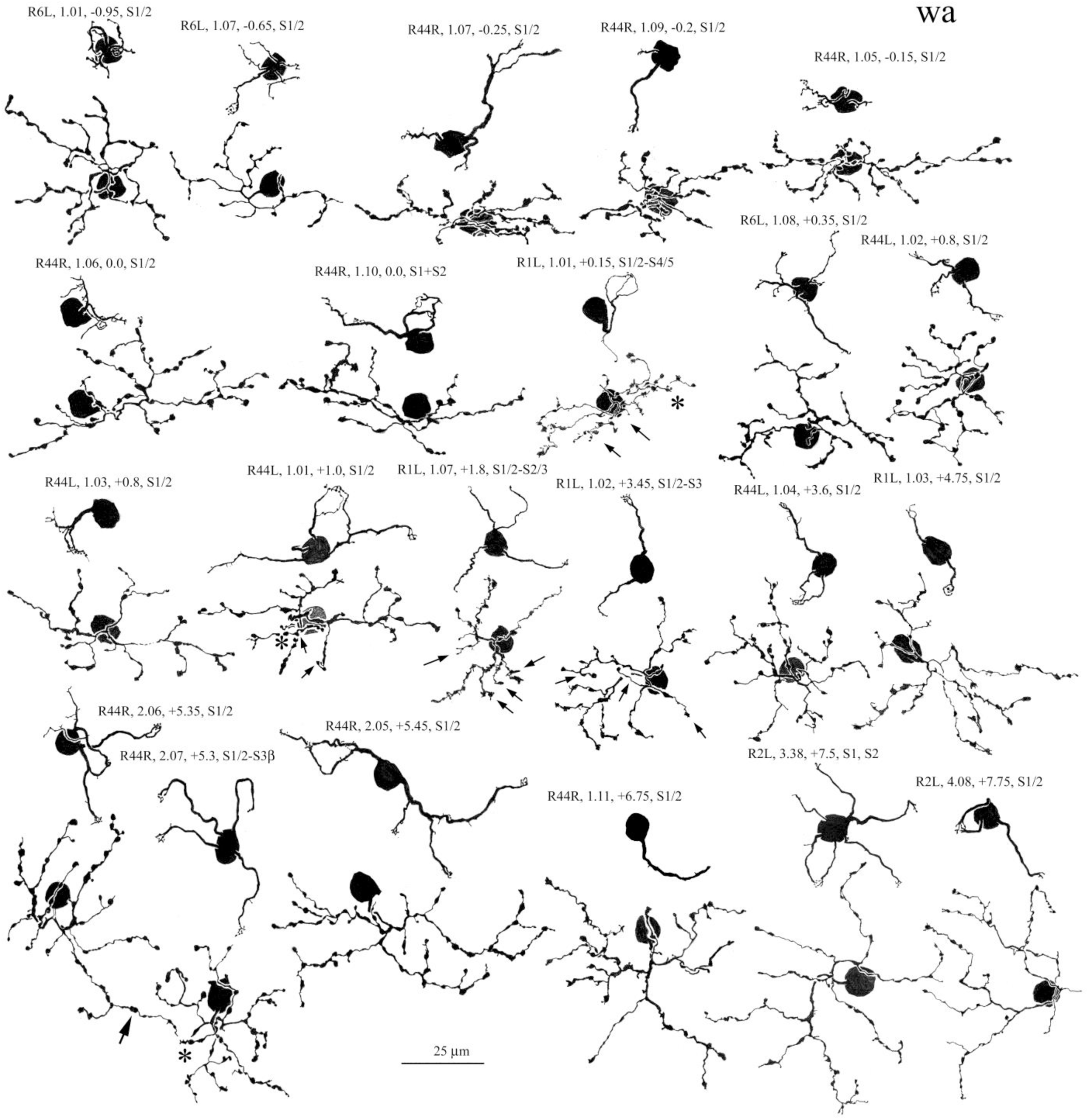
Camera lucida drawings of wide-field, type a cone bipolar cells: wa. Wide-field designation applies principally to axonal arborization. Most axon terminals are in strata (S)1 and 2. Small arrows indicate axonal branches descending to S3, and asterisks indicate processes in S4. Large arrow indicates close contact between axon terminals of adjacent cells

**FIGURE A5.**
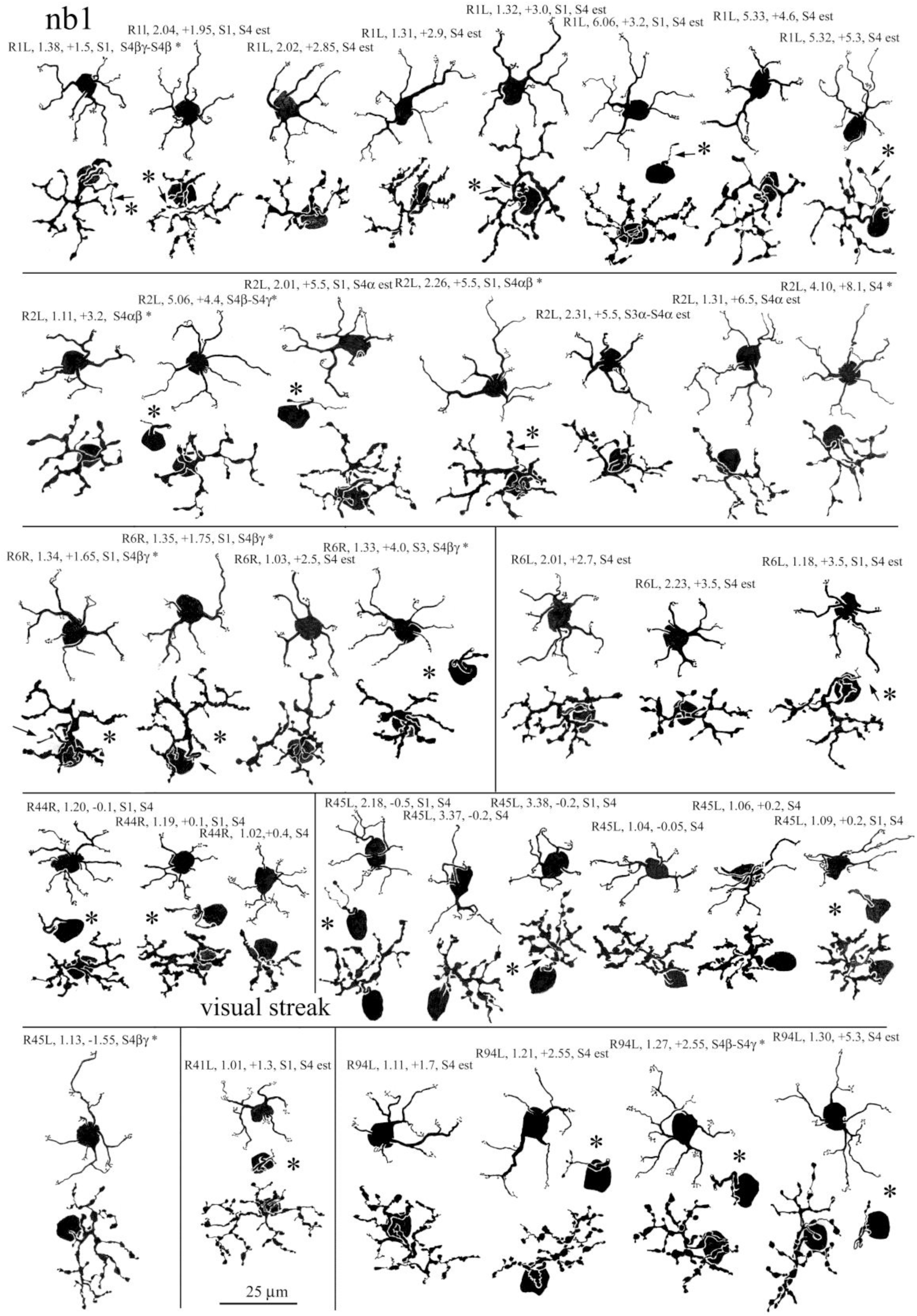
Camera lucida drawings of narrow field, type b cone bipolar cells: nb1 cells. Small arrows and asterisks indicate short axonal branches in S1, in some cases drawn separately between dendritic and axonal trees. In this and subsequent appendix figures, samples from different retinas are separated by horizontal and vertical lines. In this and subsequent figures, asterisks at the end of the cell-identifying text indicate association with a fiducial type b starburst amacrine cell

**FIGURE A6.**
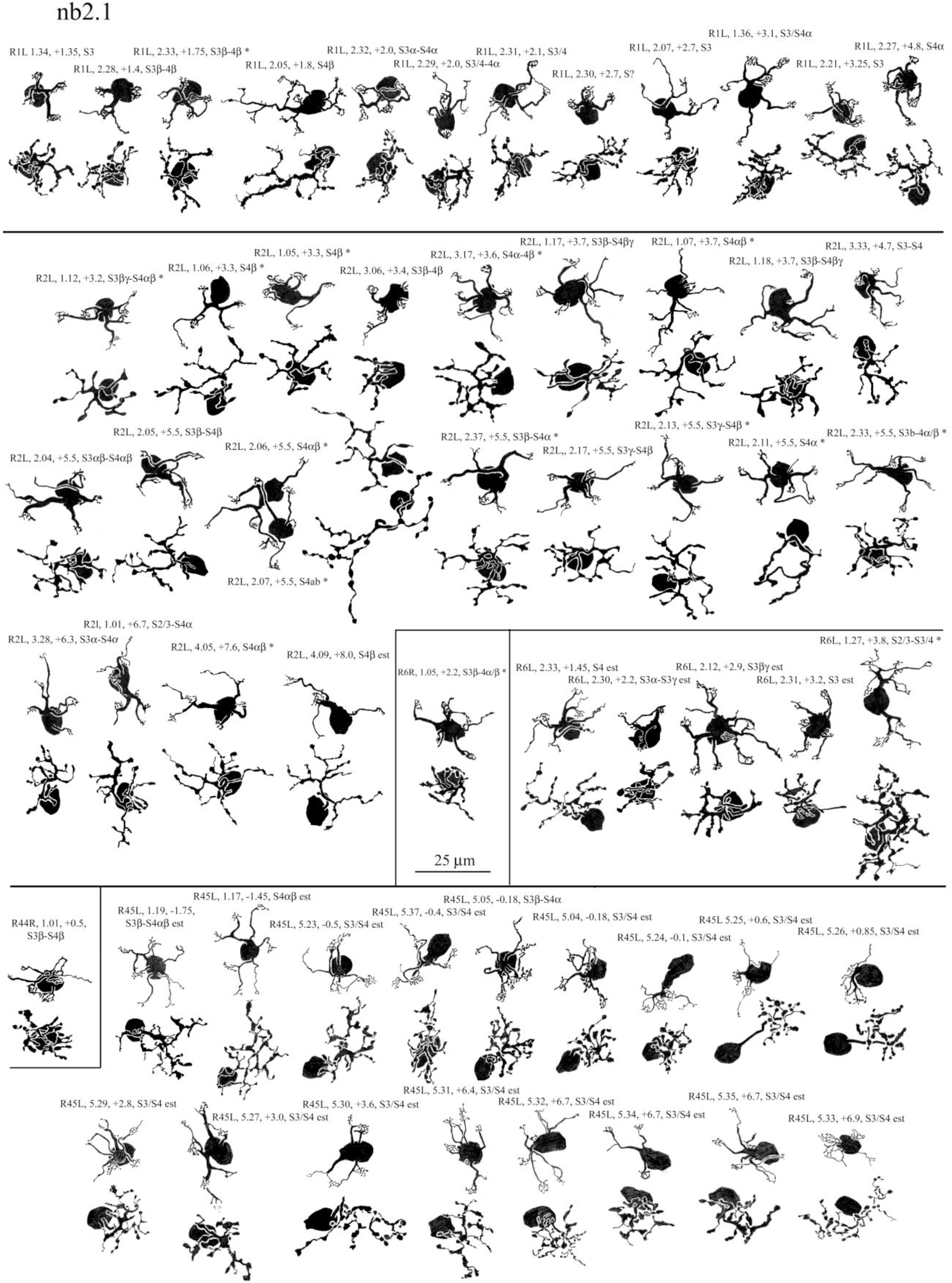
Camera lucida drawings of narrow field, type b cone bipolar cells: nb2.1 cells

**FIGURE A7.**
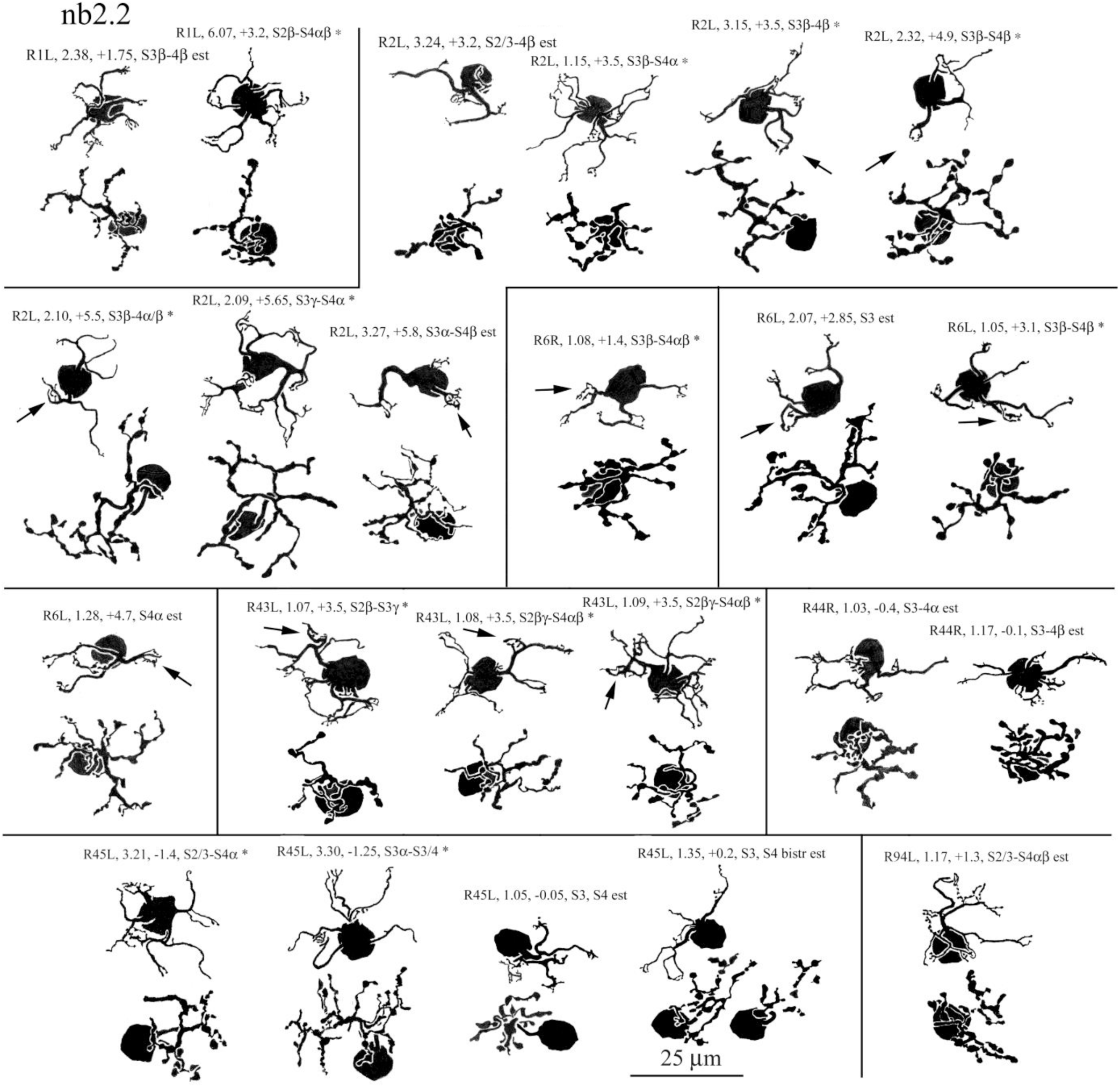
Camera lucida drawings of narrow field, type b cone bipolar cells: nb2.2 cells. Arrows indicate robust, claw-like terminal dendritic appendages, characteristic of this type

**FIGURE A8.**
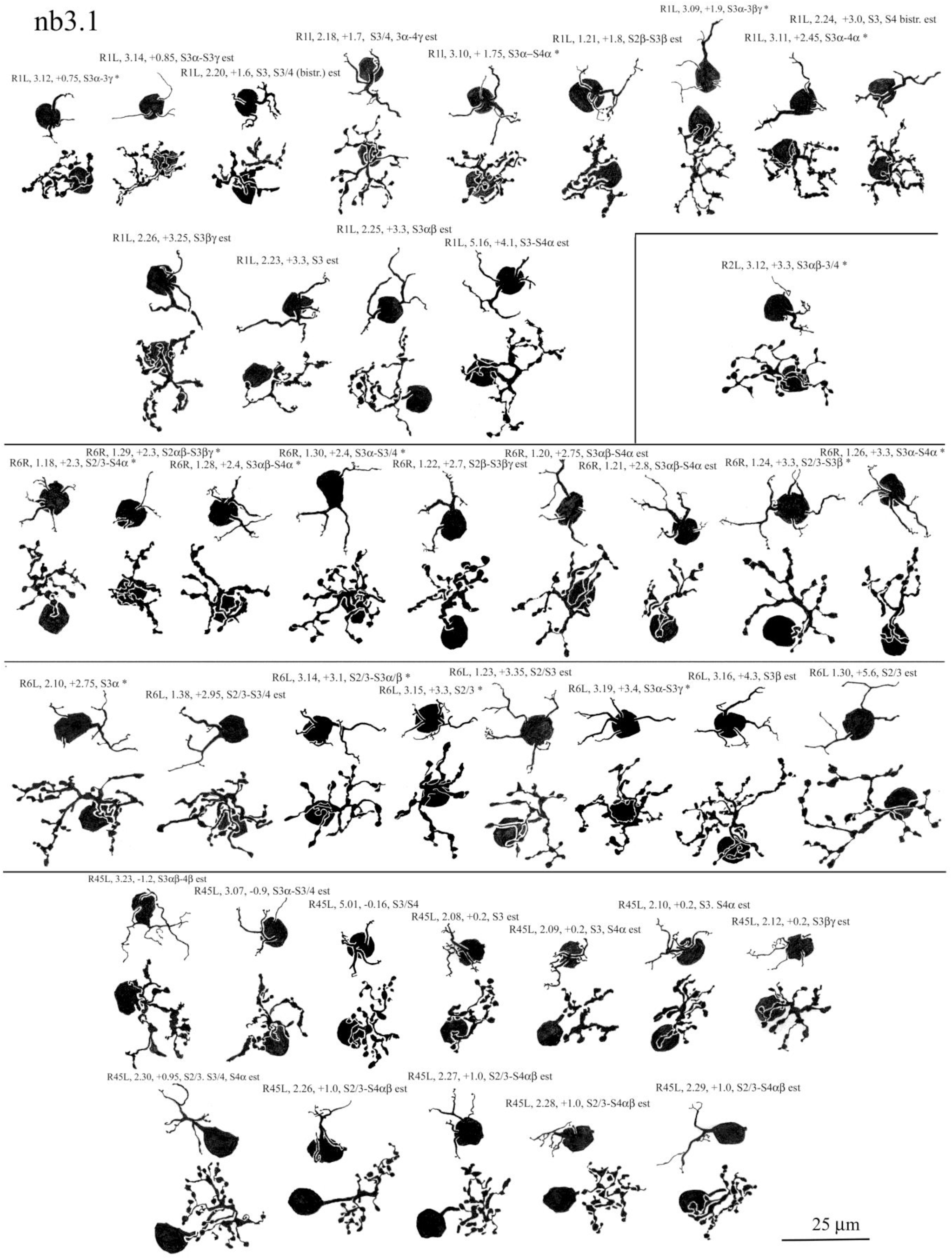
Camera lucida drawings of narrow field, type b cone bipolar cells: nb3.1 cells

**FIGURE A9.**
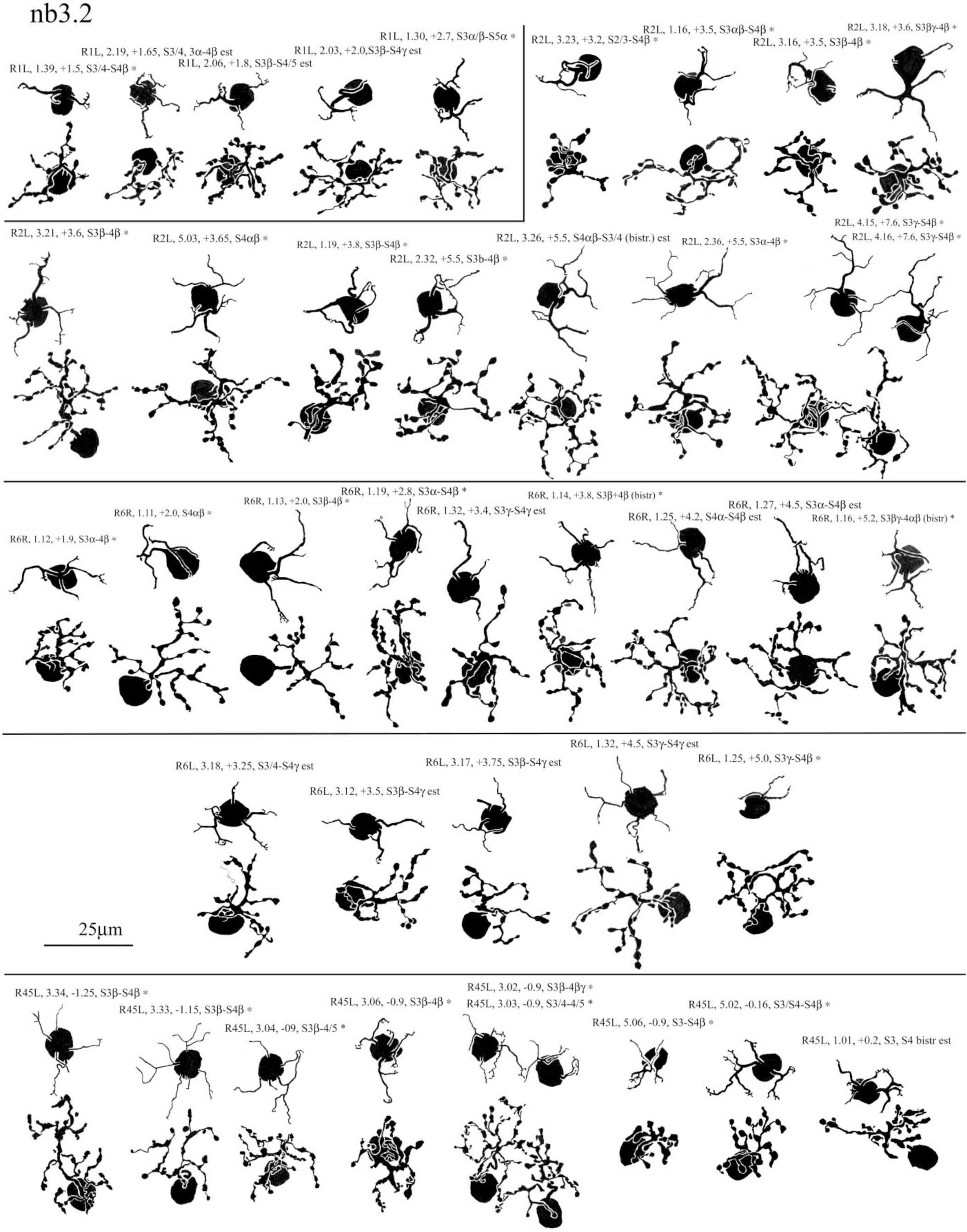
Camera lucida drawings of narrow field, type b cone bipolar cells: nb3.2 cells

**FIGURE A10.**
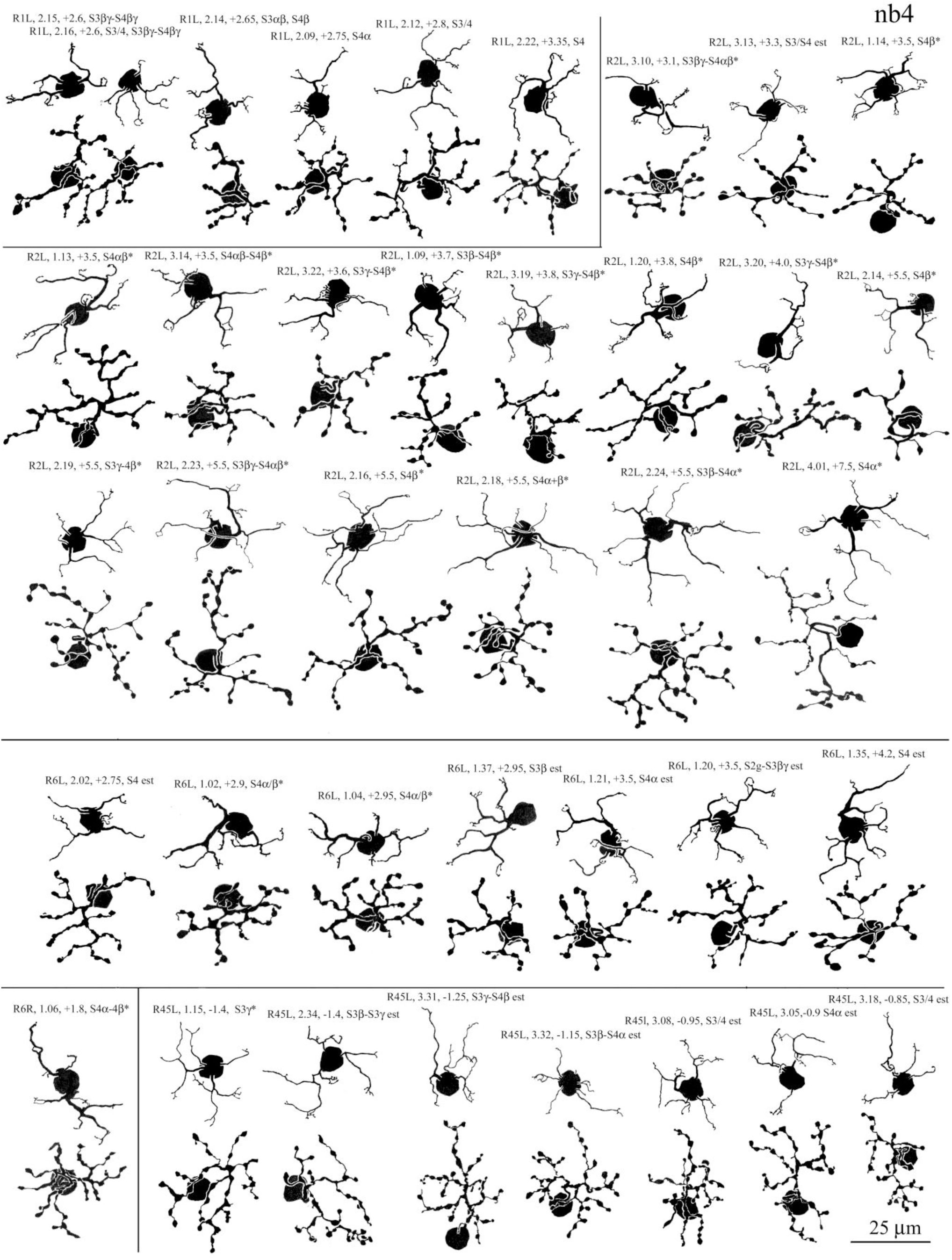
Camera lucida drawings of narrow field, type b cone bipolar cells: nb4 cells

**FIGURE A11.**
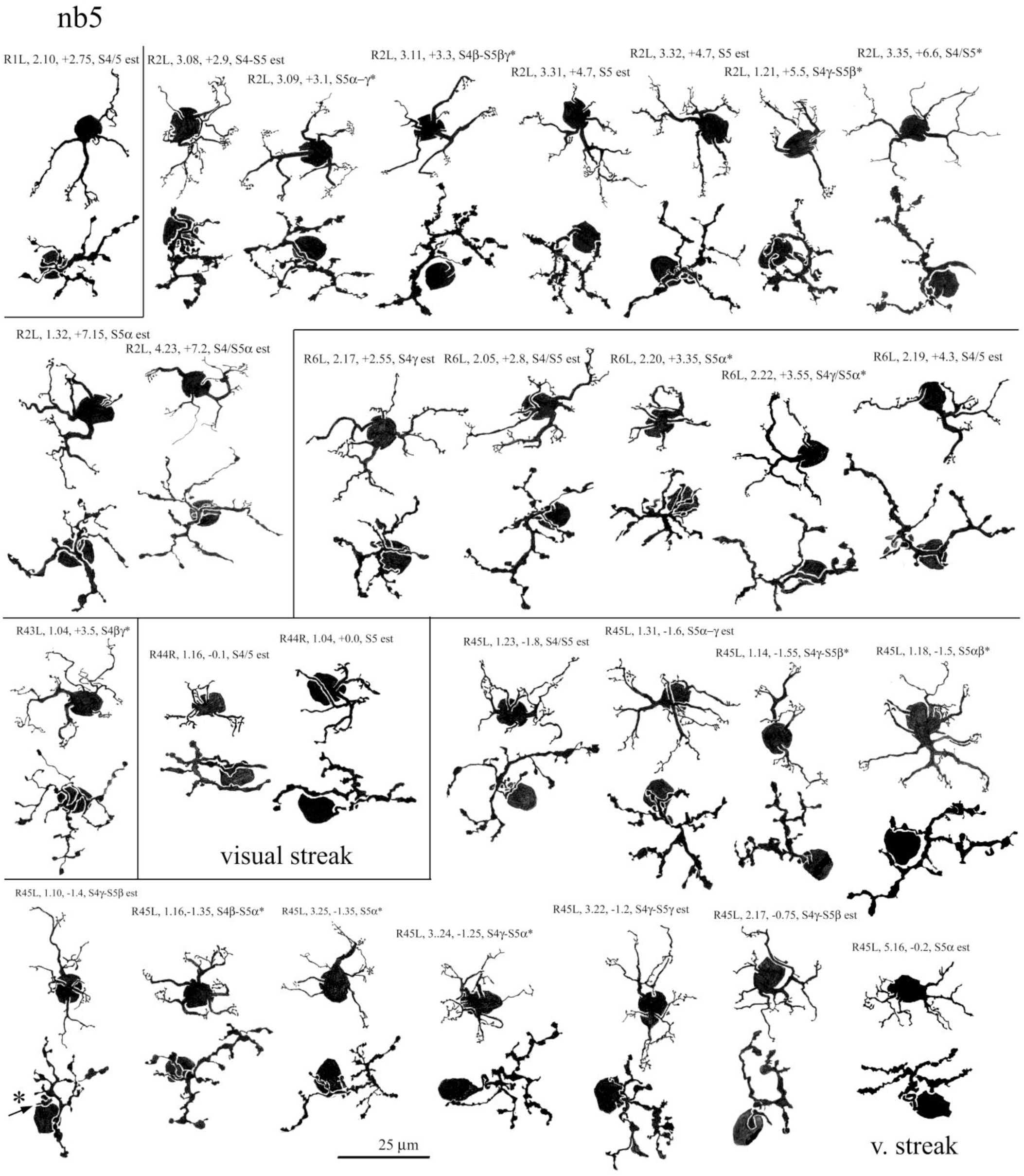
Camera lucida drawings of narrow field, type b cone bipolar cells:nb5 cells

**FIGURE A12.**
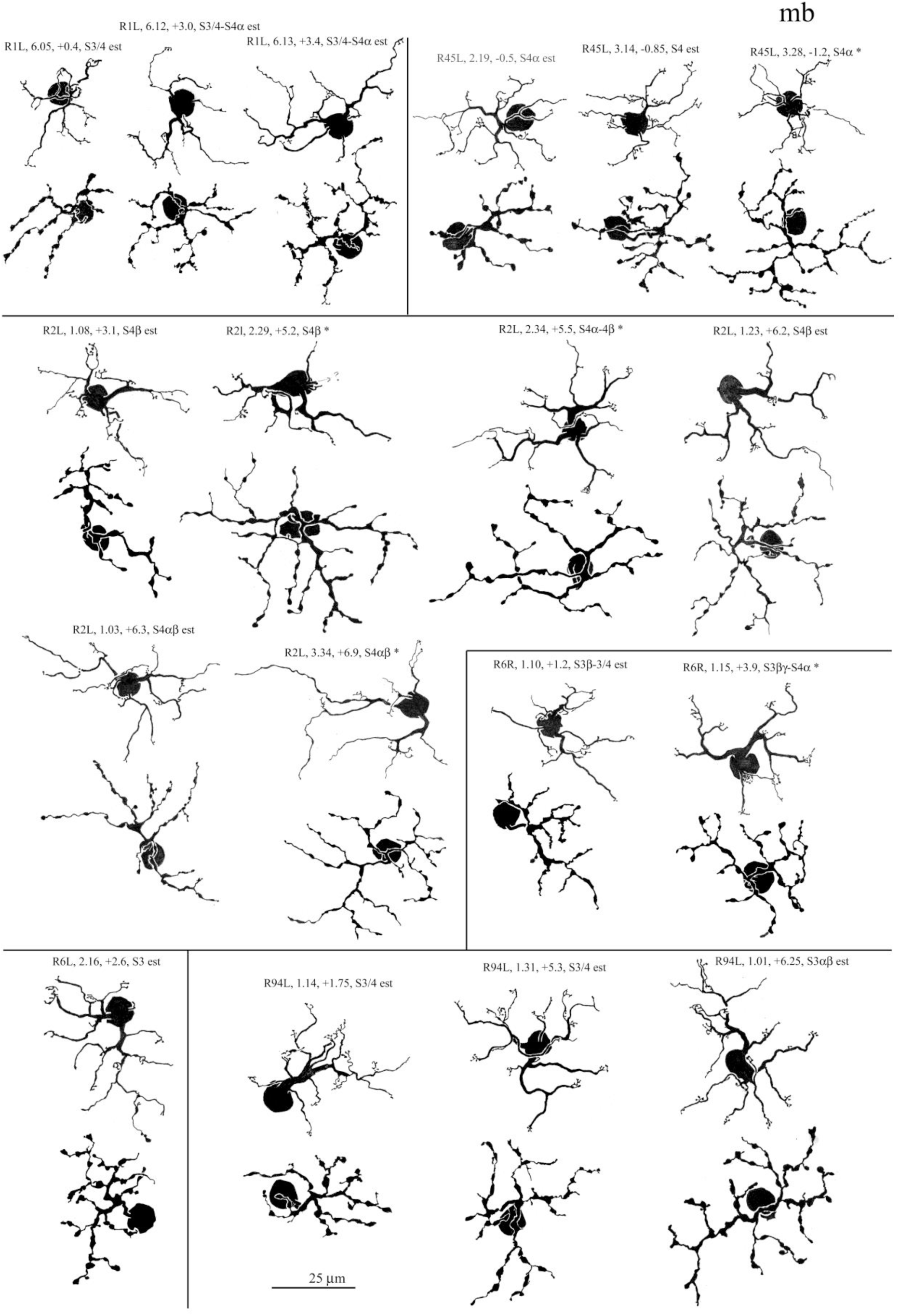
Camera lucida drawings of medium-wide field, type b cone bipolar cells: mb cells

**FIGURE A13.**
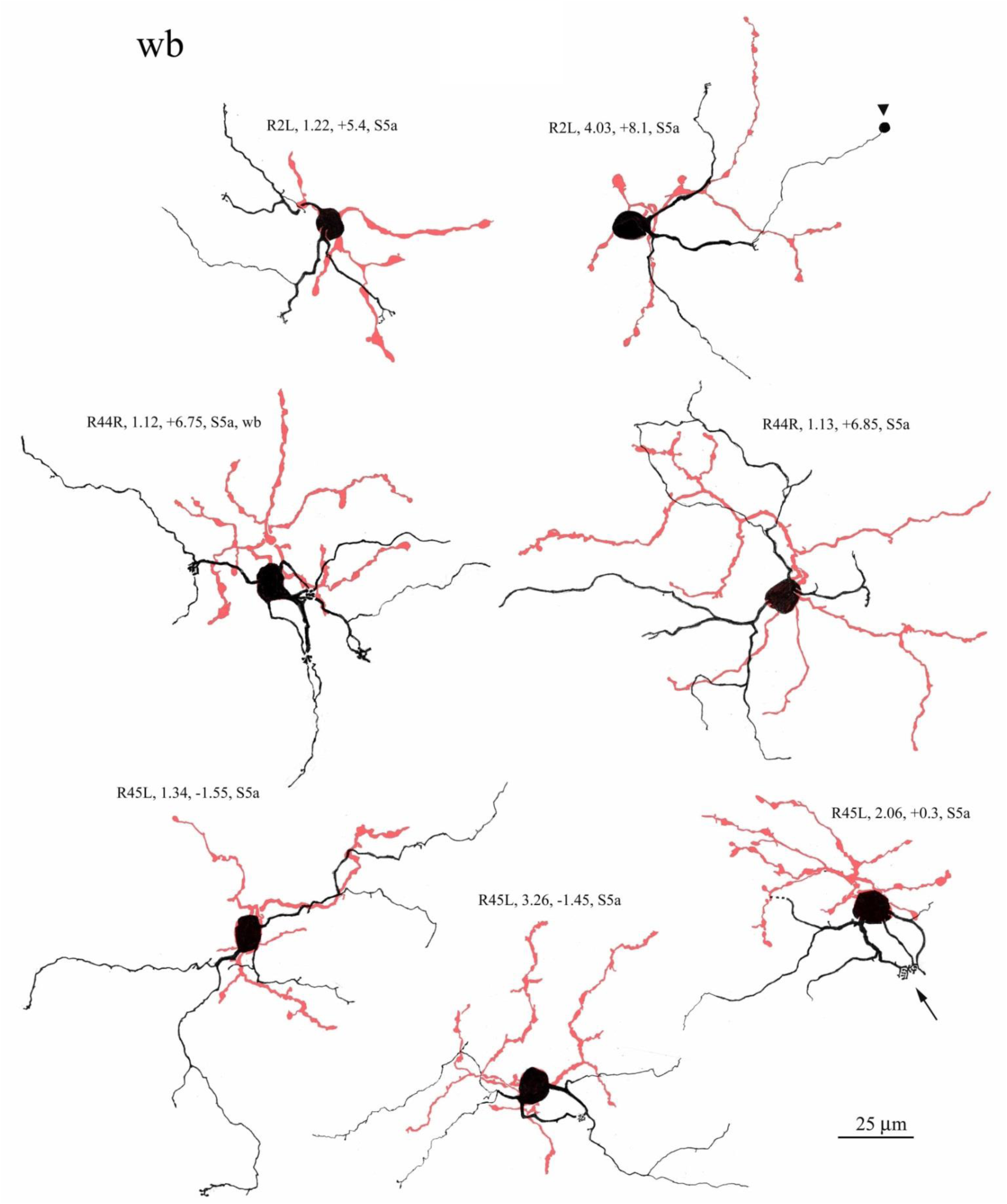
Camera lucida drawings of wide-field, type b cone bipolar cells: wb cells. Dendrites drawn in black, axon terminals drawn in red. Arrowhead indicate small spherule at the end of dendritic branch, probably an artifact of fixation, rather than a rod spherule. Arrowhead indicates dendritic convergence upon a single, short wavelength cone, in a profusion of digitiform terminal appendages, characteristic of wb cells

**FIGURE A14.**
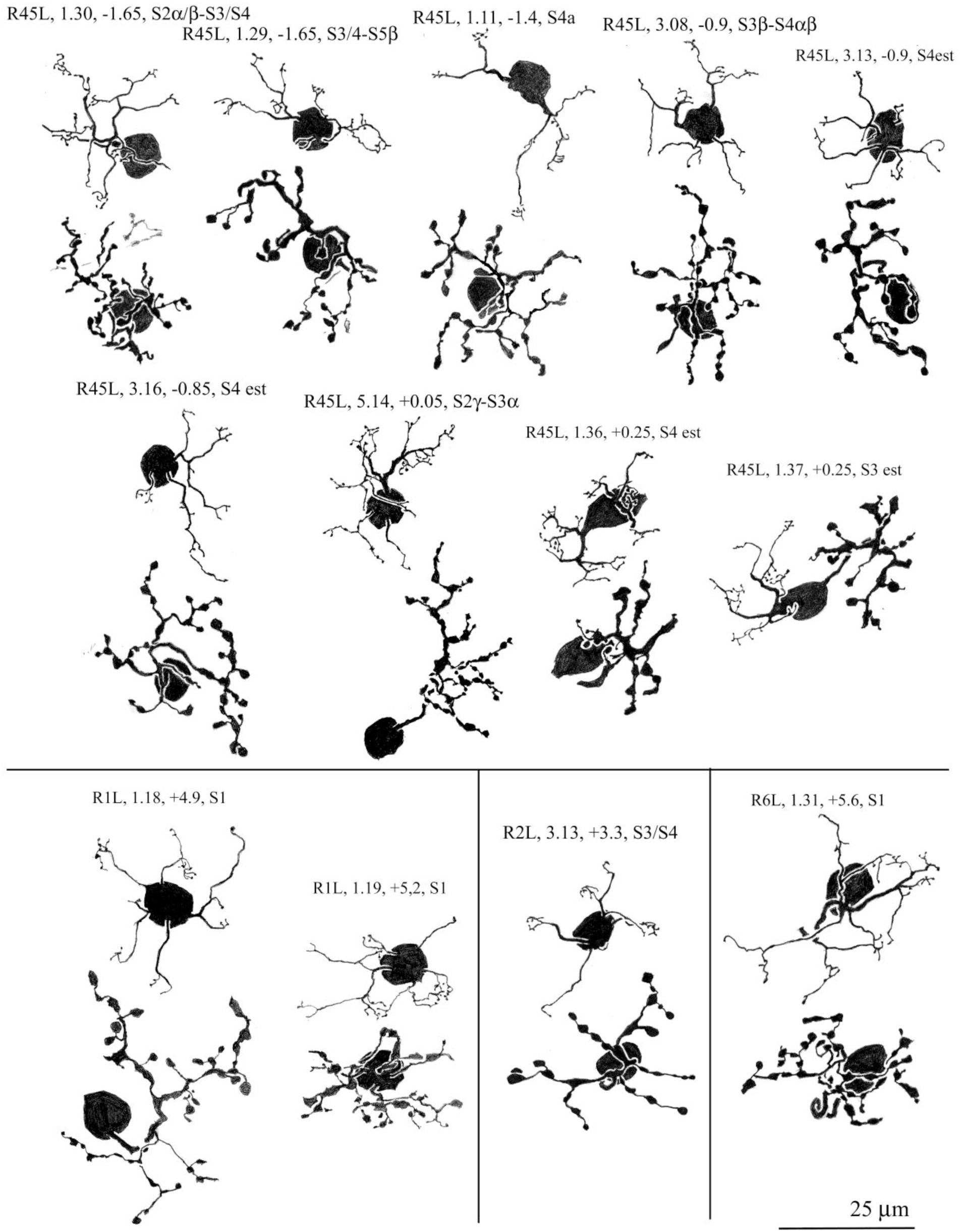
Camera lucida drawings of cone bipolar cells, both type a and type b, characterized by festoons of spinous appendages, along the course of dendrites and that they’re termini. Most have lobulated axon terminal appendages. These could not be classified as a separate type or types, because of insufficient numbers, and location in one retina primarily

**FIGURE A15.**
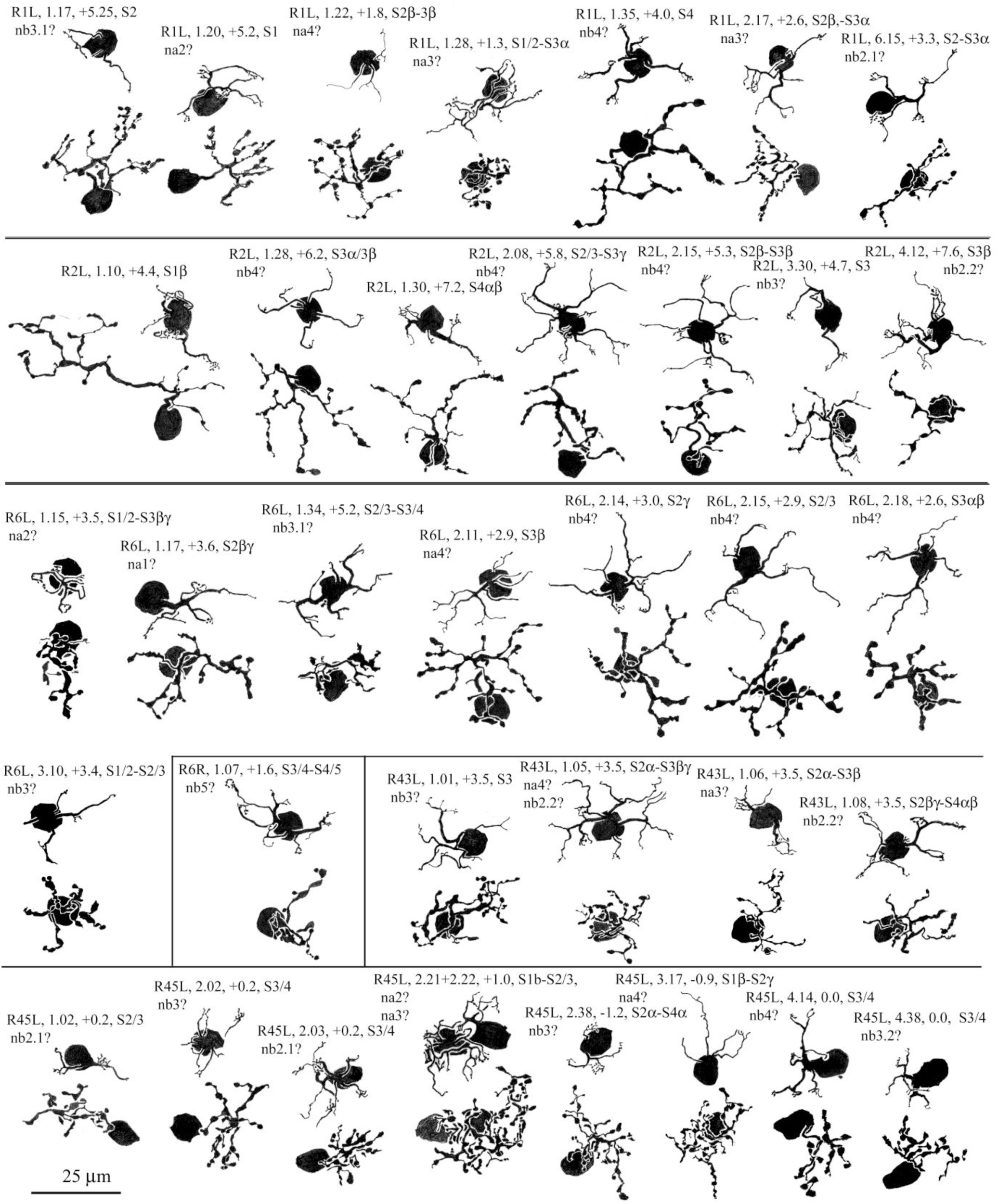
Camera lucida drawings of cone bipolar cells, both type a and type b, that could not be classified, primarily because of their dendritic morphology, intermediate between two or more types

